# Lognormal distributions capture site-specific variability in enteric virus concentrations in wastewater

**DOI:** 10.1101/2025.03.25.645242

**Authors:** Chaojie Li, Tamar Kohn, Shotaro Torii, Htet Kyi Wynn, Alexander J Devaux, Charles Gan, Timothy R. Julian, Émile Sylvestre

## Abstract

As more data on virus concentrations in influent water from wastewater treatment plants (WWTPs) becomes available, establishing best practices for virus measurements, monitoring, and statistical modelling can improve the understanding of virus concentration distributions in wastewater. To support this, we assessed the temporal variability of norovirus, adenovirus, enterovirus, and rotavirus concentrations in influent water across multiple WWTPs in Switzerland, the USA, and Japan. Our findings demonstrate that the lognormal distribution accurately predicts temporal variations in concentrations for all viruses at all sites, outperforming the gamma and Weibull distributions that do not capture high variability. However, important differences in variability and uncertainty were observed across systems, underscoring the need for site-specific assessments. Using lognormal parameters, we identified optimal monitoring frequencies to balance cost-effectiveness and precision. For most sites, weekly monitoring would be sufficient to estimate the annual average concentration of enteric viruses within a 95% confidence interval of 0.5-log10. We examined the mechanistic basis of the lognormal distribution, highlighting processes that drive its prevalence and shape the behavior of its upper tail. By integrating these insights, this study provides a novel statistical foundation for optimizing virus monitoring frameworks and informing public health interventions targeting wastewater systems.

## 1 Introduction

Enteric viruses, such as norovirus, rotavirus, and enterovirus, can enter wastewater systems from fecal discharge, posing public health risks when wastewater is inadequately managed. Infected people shed these viruses into sewage systems via toilets, sinks, and other discharge points. Combined sewers may also receive viruses from stormwater runoff contaminated by human and animal feces. Wastewater treatment plant (WWTP) processes can inactivate or remove viruses before discharging treated water into the environment, but their efficacy varies by treatment processes and virus type (Blatchley III et al., 2007). Conventional treatment plants typically employ primary treatment (physical) and secondary treatment (biological) steps, which typically reduce enteric virus concentrations by 1–3 log10 (Kitajima et al., 2014; Zhang et al., 2016). This reduction may not always be sufficient to manage infection risks for people exposed to treated wastewater (Xagoraraki et al., 2014). Quantitative microbial risk assessment (QMRA) can help inform infection risks to support the design and implementation of risk reduction strategies, but reliable virus concentration data is required to provide accurate risk estimates. Establishing best practices for virus measurements, monitoring, and statistical modelling can improve the understanding of virus concentration distributions and support QMRA in wastewater management.

Although quantitative PCR (qPCR) and digital PCR (dPCR) methods are increasingly used to monitor viruses in wastewater due to their sensitivity and specificity, these molecular methods introduce uncertainties related to the accuracy of quantitative results (Petterson et al., 2015). Factors such as adsorption to particles, PCR inhibition from organic matter, and losses during sample concentration and nucleic acid extraction can affect the reliability of measurements (Li et al., 2010). These uncertainties can be accounted for by correcting measurements based on the recovery rate of spiked surrogate viruses, such as murine hepatitis virus (MHV) (Ye et al., 2016) or non-enveloped viruses like MS2 and PhiX174 (Pecson et al., 2022). However, even though spiked viruses are routinely used as process controls, this data is rarely applied to correct concentration estimates for enteric viruses (Petterson et al., 2015).

In addition, the value of data on virus concentration data depends on the frequency of sample collection. Monitoring intervals, which can range from daily to monthly intervals (Daelman et al., 2013), should be guided by monitoring objectives, such as characterizing temporal variability in virus concentrations over a year or detecting short-term peaks indicative of viral outbreaks. However, high-frequency monitoring can be impractical in many contexts due to the significant costs and expertise required for virus analyses (La Rosa and Muscillo, 2013). Therefore, cost-effective monitoring strategies and statistical models that balance precision and practicality are essential to improve the utility of virus monitoring programs.

Statistical approaches for selecting the most suitable distributions, accounting for analytical recovery rates, and determining optimal monitoring frequencies have been developed for faecal indicator and protozoan pathogens in drinking water sources (Sylvestre et al., 2020; Sylvestre et al., 2021). However, these approaches have not been systematically adapted for enteric viruses in wastewater, where distinct challenges, such as higher variability and the influence of virus-specific shedding patterns, must be addressed. By leveraging these methods and tailoring them to enteric viruses, this study aims to fill this gap and provide a stronger statistical foundation for monitoring enteric viruses in wastewater.

In this study, we investigated temporal variations in enteric virus concentrations in influent wastewater to advance monitoring strategies. Using original data sets from two municipal WWTPs in Switzerland, as well as literature data reported for five WWTPs in the USA (Pecson et al., 2022) and one in Japan (Kazama et al., 2017), we characterized temporal variability for multiple virus types in diverse geographic contexts. We proposed candidate parametric distributions to model the data and applied information criteria to evaluate their suitability. Additionally, we assessed the impact of incorporating sample-specific analytical recovery rates and varying monitoring frequencies on model fidelity.

## 2 Material and methods

### 2.1 Rationale for sites selection and virus monitoring strategies

Data sets from eight wastewater treatment plants (WWTPs) were selected for this study. Two WWTPs in Switzerland—the STEP Vidy WWTP in Lausanne and the ARA Werdhölzli WWTP in Zurich—provided original data, while previously published data were obtained from Matsushima, Japan (Kazama et al., 2017), and five WWTPs in California, USA [Los Angeles County Sanitation Districts (LACSD), City of Los Angeles Sanitation and Environment (LASAN), Orange County Sanitation District (OCSD), City of San Diego (SD), and San Francisco Public Utilities Commission (SFPUC)] (Pecson et al., 2022). These sites were chosen to represent various geographical locations and population sizes (Table 1).

**Table 1.**
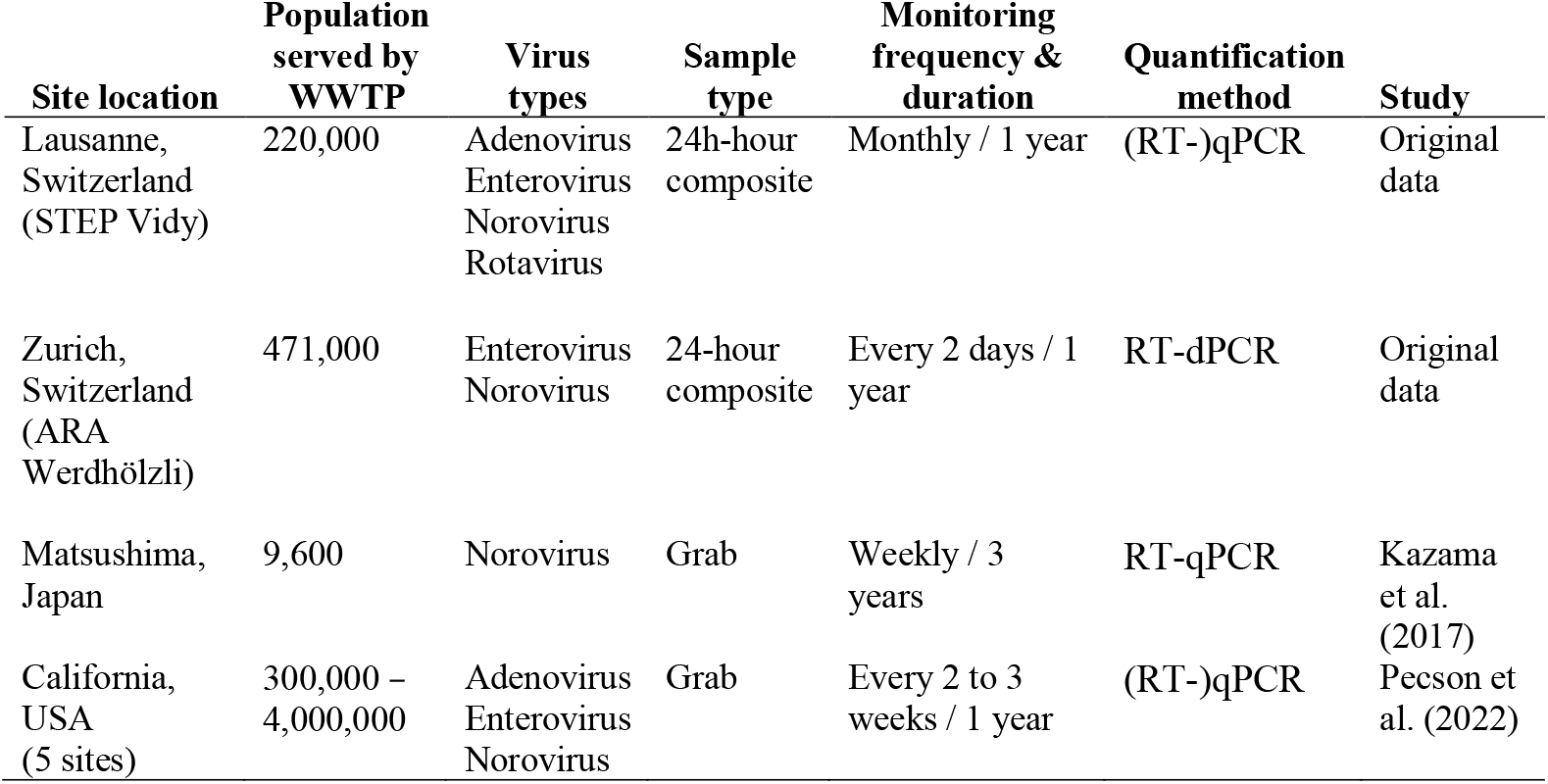
Locations of municipal wastewater treatment plants (WWTPs) and corresponding virus types analyzed in influent wastewater at these sites.

| Site location | Population served by WWTP | Virus types | Sample type | Monitoring frequency & duration | Quantification method | Study |
| --- | --- | --- | --- | --- | --- | --- |
| Lausanne, Switzerland (STEP Vidy) | 220,000 | Adenovirus<br>Enterovirus<br>Norovirus<br>Rotavirus | 24h-hour composite | Monthly / 1 year | (RT-)qPCR | Original data |
| Zurich, Switzerland (ARA Werdhölzli) | 471,000 | Enterovirus<br>Norovirus | 24-hour composite | Every 2 days / 1 year | RT-dPCR | Original data |
| Matsushima, Japan | 9,600 | Norovirus | Grab | Weekly / 3 years | RT-qPCR | Kazama et al. (2017) |
| California, USA (5 sites) | 300,000 – 4,000,000 | Adenovirus<br>Enterovirus<br>Norovirus | Grab | Every 2 to 3 weeks / 1 year | (RT-)qPCR | Pecson et al. (2022) |

Only data quantified by molecular methods—(RT-)qPCR or RT-dPCR—with documented process controls and recoveries were included to maintain methodological consistency. All data sets also fulfilled the following criteria: (i) ≥12 months of routine sampling, (ii) consistent quantification of at least one target enteric virus, and (iii) sufficient sample size (≥10 positive detections) for parametric modelling. The viruses measured included adenovirus, enterovirus, norovirus, and rotavirus at Lausanne; enterovirus and norovirus at Zurich; norovirus at Matsushima; and adenovirus, enterovirus, and norovirus at the five WWTPs in California, USA. Certain studies involved extended monitoring campaigns (e.g., the multi-year, weekly sampling in Matsushima), whereas others had higher-frequency monitoring (e.g., the every-two-day sampling at ARA Werdhölzli). Analyzing data under these different monitoring regimes allowed us to examine whether the modelling framework proposed in this work holds across both short-term, high-frequency datasets and longer-term monitoring programs. Consequently, the monitoring recommendations derived in this work should be broadly applicable to various contexts.

### 2.2 Sampling and quantification of enteric viruses

#### 2.2.1 Original data from Swiss WWTPs

For the WWTP in Lausanne, 24-hour composite wastewater influent samples were collected monthly between November 2018 and October 2019. Detailed sampling processing procedures, nucleic acid extraction methods, and (RT-)qPCR quantification are described in Li et al. (2023). In brief, approximately 600 ml of raw sewage was first pre-filtered and further concentrated using a centrifugal filter unit, resulting in a final concentrate of about 2 ml. Nucleic acids were extracted using the QIAamp Viral RNA Mini Kit and stored at –80 °C prior to qPCR/RT-qPCR quantification of adenovirus, enterovirus, norovirus, and rotavirus. All qPCR/RT-qPCR assays were carried out in duplicate, with DNA gBlocks used as standards for absolute quantification. The primer and probe sequences are reported in Li et al. (2023).

For the WWTP in Zurich, 180 24-hour composite influent samples were collected between February 1, 2021, and January 29, 2022. After collection, the wastewater samples were shipped on ice and stored at 4 °C for up to eight days before processing. The collected samples were processed either by Protocol 1 (ultrafiltration followed by RNA extraction using QIAamp viral RNA Mini Kit, used before November 10, 2021) (Huisman et al., 2022) or Protocol 2 (total nucleic acid extraction, samples collected after November 30, 2021) (Nadeau et al., 2023). Samples collected between November 10 and November 30, 2021, were processed by both protocols to establish a conversion factor that corrects for inter-method differences in observed concentration. To adjust for differences in method performance, a correction factor was applied to measurements from Protocol 1. Specifically, data from Protocol 1 were adjusted by multiplying by 1.17 for EV and 2.65 for NoV GII. The extracted RNA was stored at −80 °C for up to 1.5 years before being measured on dPCR.

A duplex RT-dPCR assay for human enterovirus (EV) and norovirus genogroup II (NoV GII) RNA quantification was optimized by adapting the thermal cycling conditions and primer and probe concentrations from previously described RT-qPCR assays (Kageyama et al., 2003; Loisy et al., 2005; Monpoeho et al., 2000; Tsai et al., 1993). Ten-fold diluted extracts were used as RNA templates. The assay was performed in 12 µL reaction mixtures using the QIAcuity OneStep Advanced Probe Kit (QIAGEN) and 8.5k 96-well Nanoplates (QIAGEN). The duplex RT-dPCR mixture contained 3 μL of 4x OneStep Advanced Probe Master Mix, 0.12 µl of OneStep RT Mix, 1.5 µl of GC Enhancer, 1000 nM forward primer (EV: 5’-CCTCCGGCCCCTGAATG -3’, NoV GII: 5’-ATGTTCAGRTGGATGAGRTTCTCWGA -3’), 1000 nM reverse primer (EV: 5’-ACCGGATGGCCAATCCAA -3’, NoV GII: 5’-TCGACGCCATCTTCATTCACA -3’), 500 nM probe (EV: 5’- HEX - CGGAACCGACTACTTTGGGTGTCCGT-BHQ1 -3’, NoV GII: 5’- FAM-AGCACGTGGGAGGGCGATCG-TAMRA -3’), 3 or 4 µl of template RNA, and DNase/RNase free water. All RT-dPCR reactions were performed in duplicate. The nanoplate was loaded onto the QIAcuity One, 2-plex Device (Qiagen). The thermal cycles include an RT step at 50 °C for 60 min, 95 °C for 5 min for enzyme activation, and followed by 45 cycles of denaturation (95 °C for 15 s) and annealing/extension (at 60 °C for 60 s). In each run, a negative control (no template) and a positive control (i.e., synthetic DNA gBlock® containing the target sequence, Integrated DNA Technologies, Coralville, IA, USA) were included. Quantities were expressed as genome copies (GC µL^-1^) per reaction. The RT-dPCR assays were performed using automatic settings for the threshold and baseline. Samples showing >5-fold difference between the duplicated reactions were excluded (3 out of 180 samples) from further analysis. Further quality control data for the duplex RT-dPCR assay is given in the Supporting Information. The completed dMIQE (digital Minimum Information for Publication of Quantitative Digital PCR Experiments) checklist for this RT-dPCR is available in Huisman et al. (2022).

To assess the recovery efficiency for 91 wastewater samples collected at the Zurich WWTP, Murine Hepatitis Virus (MHV) was spiked into replicate aliquots of 50 ml wastewater at a concentration of approximately 1×10^6^ GC per 50 ml^-1^. The preparation of MHV stock solutions is described in Fernandez-Cassi et al. (2021). MHV was determined using the primers and probes described in Fernandez-Cassi et al. (2021), but using a protocol modified for RT-dPCR and a Naica System (Stilla Technologies, Villejuif, France), using the qScript XLT 1-Step RT-PCR Kit (QuantaBio, Beverly, Massachusetts, United States).

#### 2.2.2 Literature data from WWTPs in the USA and Japan

The methods used for sample collection, processing, and quantification are summarized below; full methodological details are provided in the original studies cited.

In Pecson et al. (2022), 4-L grab samples of raw wastewater were collected at each site every two to three weeks between December 2019 and January 2021. Commercial laboratories analyzed samples using standardized methods. Upon receipt, they were refrigerated at 4 ^°^C and processed within 72 hours of collection. Norovirus (genogroups I and II) and enterovirus were quantified using RT-qPCR, and adenovirus was quantified using qPCR. Nucleic acid extraction was performed with the Zymo Quick-DNA/RNA Viral Kit, and assays were run in triplicate with the average gene copies reported. To evaluate analytical recovery rates, MS2 and PhiX174 were spiked into 1000 ml of the sample at approximately 10^8^ plaque-forming units (PFU). A minimum recovery efficiency threshold of 1% was set for all matrix spikes.

In Kazama et al. (2017), 1-L grab samples of raw wastewater were collected weekly from a single WWTP between 2013 and 2016. Samples were transported to the laboratory on ice and stored in a deep freezer on the same day as collection. Norovirus genogroups I and II were quantified using RNA extraction with the QIAamp Viral RNA Mini Kit, followed by RT-qPCR. Murine norovirus (MNV) was spiked in each sample as a whole-process control and quantified by qPCR. A minimum recovery efficiency threshold of 1% was applied to all matrix spikes.

For all data sets, concentrations of norovirus genogroups I and II were summed to analyze and compare the overall distribution of norovirus in wastewater across locations.

### 2.3 Statistical modelling of temporal variations in virus concentrations

#### 2.3.1 Model formulation

To model the variability and uncertainty in virus concentrations, parametric probability distributions were fitted to the monitoring data. The analysis assumed stationarity, meaning the mean, variance, and autocorrelation of concentrations remain constant over time. This approach estimates the annual distribution of enteric virus concentrations, as this is typically required to calculate annual infection risks in QMRA. If seasonal or outbreak-driven variability were to be explicitly considered, separate distributions for specific periods, such as winter and summer, would be necessary.

Two mixed Poisson distributions, the Poisson lognormal distribution (PLN) and the Poisson gamma distribution (PGA), were used to model concentration variability using virus count and processed water volume data. In this framework, the Poisson distribution accounts for the uncertainty associated with the spatial (random) dispersion of viruses in the water sample, and the continuous distribution (gamma or lognormal) characterizes temporal variation in virus concentrations.

Virus concentrations in wastewater are expected to follow a lognormal distribution due to multiplicative processes such as shedding, decay, aggregation, and disaggregation, which, according to the Central Limit Theorem, result in a normal distribution for the logarithm of the concentrations. The probability function of the Poisson lognormal distribution for a virus count *x* is:

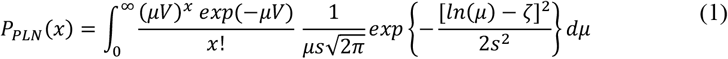

where *V* is the sample volume, *μ* is the mean virus concentration in the sample, *ζ* is the mean of the natural logarithm of the virus concentration and *s* is the standard deviation of the natural logarithm of the virus concentration.

The gamma distribution, on the other hand, as a thinner upper tail compared to the lognormal, reflecting a more rapid decline in probability for high concentrations. This makes it a suitable candidate for modelling virus concentrations in systems where high concentrations are constrained by physical limits, such as dilution in wastewater systems, or a biological limit, such as shedding saturation. The probability function of the Poisson gamma distribution for a virus count *x*is given by:

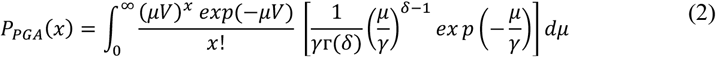

where *γ* and *δ* are parameters of a gamma distribution which has a mean of *γδ* and a variance of *δγ*^2^.

In cases where original observations were not available, three continuous probability distributions (lognormal, gamma, Weibull) were employed to describe the temporal variations in reported concentrations. The probability function of the Weibull distribution for a reported virus concentration z is:

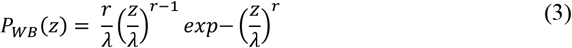

where *r* is the shape parameter and *λ* is the scale parameter.

#### 2.3.2 Model implementation

For statistical inference, a frequentist method (maximum likelihood estimator) and Bayesian methods (Markov chain Monte Carlo) were applied to estimate parameters of the lognormal, gamma, and Weibull distribution, while only the Bayesian method was used to estimate parameters of the mixed Poisson models. Bayesian inference was carried out as described in (Sylvestre et al., 2021). The priors selected for the lognormal distribution parameters were: a diffuse uniform prior for the mean with a uniform distribution with a lower of −10 and upper of 10 and a vague exponential prior for the standard deviation with an exponential distribution of 0.1. Three Markov chains were run for 10^6^ iterations after a burn-in phase of 10^4^ iterations. All statistical computations were performed using R (v4.1.2).

#### 2.3.3 Model comparison

Temporal variations in sewage virus concentrations were represented with complementary cumulative distribution functions (CCDF) curves as these curves help visualize the behavior of the upper tail of the distributions. Goodness-of-fit criteria were specified to compare the performance of each model. Continuous distributions were compared using the deviance information criterion (DIC) and the Akaike information criterion (AIC). The two mixed Poisson models were compared using the marginal deviance information criterion (DICm) (Quintero and Lesaffre, 2018). A difference in DIC or AIC or DICm of at least three points was considered meaningful (Gelman et al., 2013).

### 2.4 Incorporation of analytical recovery rates into the model

Analytical recovery rates of other viruses that were spiked into the wastewater were also evaluated for their ability to inform inter-sample variation in recovery rates. For datasets from Pecson et al. (2022), we divided sample-specific enteric virus concentrations by the average of the sample-specific recovery rates of MS2 and PhiX174, following the approach used by Pecson et al. (2022). To evaluate the impact of the recovery rates on the distributional form, the PLN model was fitted to data with and without sample-specific correction for the recovery rate.

### 2.5 Influence of virus monitoring frequencies on model predictions

Using the Zurich dataset, we evaluated how the monitoring frequency affects predictions of virus concentrations modelled by the PLN distribution over a period of a year. To achieve this, we randomly sub-sampled the dataset to represent different monitoring intervals: monthly, every two weeks, and weekly measurements. We compared the CCDF curves for each sub-sampled dataset with those derived from the original dataset.

Furthermore, we investigated the impact of the monitoring frequency on the uncertainty of the arithmetic mean virus concentration predicted by the PLN. This analysis was conducted using the method of Olsson (2005), which estimates the confidence interval (CI) of the arithmetic mean of a lognormal distribution. This approach takes into account the sample size and the sigma (*s*) parameter values of the lognormal distribution. To quantify uncertainty, we calculated the width of the 95% CI on the arithmetic mean on the log scale.

## 3 Results

### 3.1 Variation in concentrations of viruses across locations

The time series data for enteric viruses from the eight WWTPs are shown in Figure 1. Norovirus shows a clear temporal pattern with higher concentrations in winter (blue-shaded areas) in Zurich (Panel B), and Matsushima, Japan (Panel D). In contrast, the WWTP in Lausanne (Panel A) shows relatively consistent concentrations throughout the year, while the five WWTPs in California, USA (Panel C) display high variability without clear temporal trends. In winter 2019, rotavirus peaks in Lausanne, and adenovirus peaks in California, USA. Adenovirus and enterovirus concentrations are more sporadic in Lausanne and California, showing occasional peaks without a seasonal trend.

**Fig. 1.**
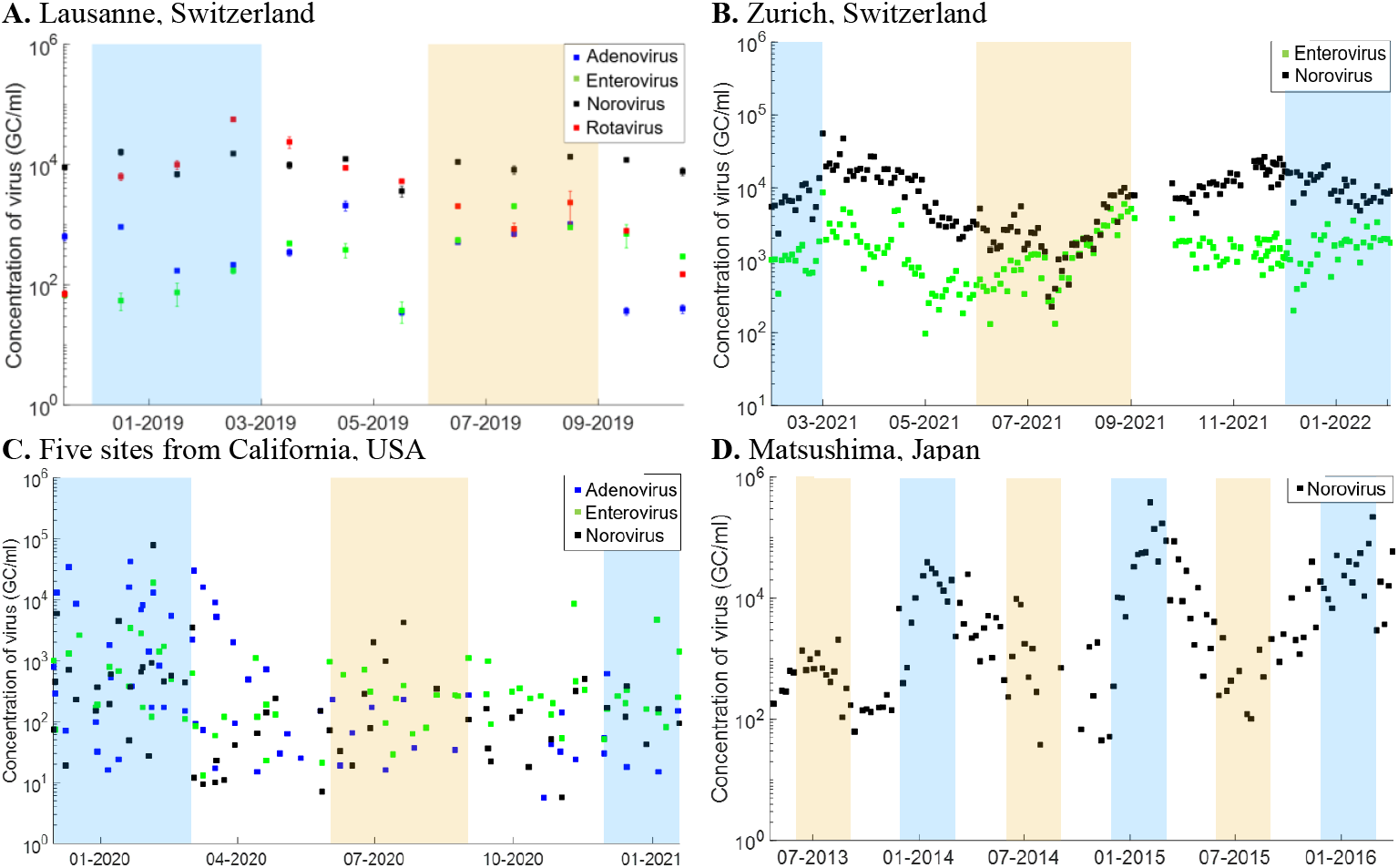
Concentrations of enteric viruses in influent water from wastewater treatment plants (WWTPs) in genome copies (GC) per millilitre (ml) wastewater in Switzerland, USA, and Japan. Blue shades indicate winter seasons, and orange shades indicate summer seasons.

Table 2 presents the arithmetic mean concentration and coefficient of variation (CV) for each virus type across Switzerland, Japan, and the USA. Arithmetic mean norovirus concentrations in Japan and Switzerland (approx. 1.0 × 10^4^ GC ml^-1^) are 10 to 100 higher than in the USA. CVs below 1.0 for Swiss sites indicate relatively stable concentrations, while CVs higher than 2.5 in Japan and the USA suggest much greater variability. Adenovirus shows high CVs across all USA sites (up to 2.1), indicating substantial fluctuations compared to Lausanne (CV of 1.0).

**Table 2.** Sample arithmetic mean concentration in genome copies (GC) per milliliter (ml) and the coefficient of variation of adenovirus (AD), enterovirus (ET), norovirus (NR), and rotavirus (RT) in influent water from wastewater treatment plants located in Switzerland, Japan, and USA.

| Virus | Location of wastewater treatment plants | Sample size | Sample mean concentration (GC ml<sup>-1</sup>) | Coefficient of variation | Lognormal parameter ($\sigma$) |
| --- | --- | --- | --- | --- | --- |
| Adenovirus | Lausanne, Switzerland | 12 | $5.6 \times 10^2$ | 1.0 | 1.5 |
| | California, USA – LACSD | 14 | $5.1 \times 10^2$ | 1.9 | 2.2 |
| | California, USA – LASAN | 14 | $6.7 \times 10^2$ | 1.8 | 1.9 |
| | California, USA – OCSD | 11 | $2.1 \times 10^3$ | 1.8 | 2.4 |
| | California, USA – SD | 11 | $9.5 \times 10^2$ | 2.1 | 2.2 |
| | California, USA – SFPUC | 13 | $1.6 \times 10^3$ | 2.1 | 2.6 |
| Enterovirus | Lausanne, Switzerland | 12 | $4.8 \times 10^2$ | 1.1 | 1.4 |
| | Zurich, Switzerland | 180 | $1.5 \times 10^3$ | 0.8 | 0.8 |
| | California, USA – LACSD | 10 | $1.8 \times 10^2$ | 0.6 | 1.5 |
| | California, USA – LASAN | 17 | $2.1 \times 10^2$ | 1.0 | 0.6 |
| | California, USA – OCSD | 13 | $5.1 \times 10^2$ | 2.6 | 1.6 |
| | California, USA – SD | 14 | $2.2 \times 10^2$ | 2.0 | 1.2 |
| | California, USA – SFPUC | 17 | $3.5 \times 10^2$ | 0.9 | 1.2 |
| Norovirus | Lausanne, Switzerland | 12 | $1.0 \times 10^4$ | 0.3 | 0.4 |
| | Zurich, Switzerland | 180 | $1.0 \times 10^4$ | 0.8 | 0.9 |
| | Matsushima, Japan | 131 | $1.8 \times 10^4$ | 2.5 | 2.1 |
| | California, USA – LACSD | 8 | $2.9 \times 10^2$ | 1.5 | 1.6 |
| | California, USA – LASAN | 14 | $1.1 \times 10^2$ | 1.3 | 1.0 |
| | California, USA – OCSD | 11 | $2.0 \times 10^3$ | 3.1 | 2.4 |
| | California, USA – SD | 15 | $3.7 \times 10^2$ | 1.6 | 2.0 |
| | California, USA – SFPUC | 13 | $3.4 \times 10^2$ | 2.5 | 1.7 |
| Rotavirus | Lausanne, Switzerland | 12 | $9.7 \times 10^3$ | 1.6 | 2.2 |

### 3.2 Performance of parametric distributions in predicting temporal variations

Figure 2 presents complementary cumulative distribution functions (CCDFs) for norovirus concentrations at four WWTPs, with mixed Poisson distributions fitted to count and volume data (Panel A) and continuous distributions fitted to reported concentrations (Panel B). The complete CCDF curves for all datasets are shown separately in Figure S3 (discrete mixed Poisson distributions) and Figure S4 (continuous distributions).

**Fig. 2.**
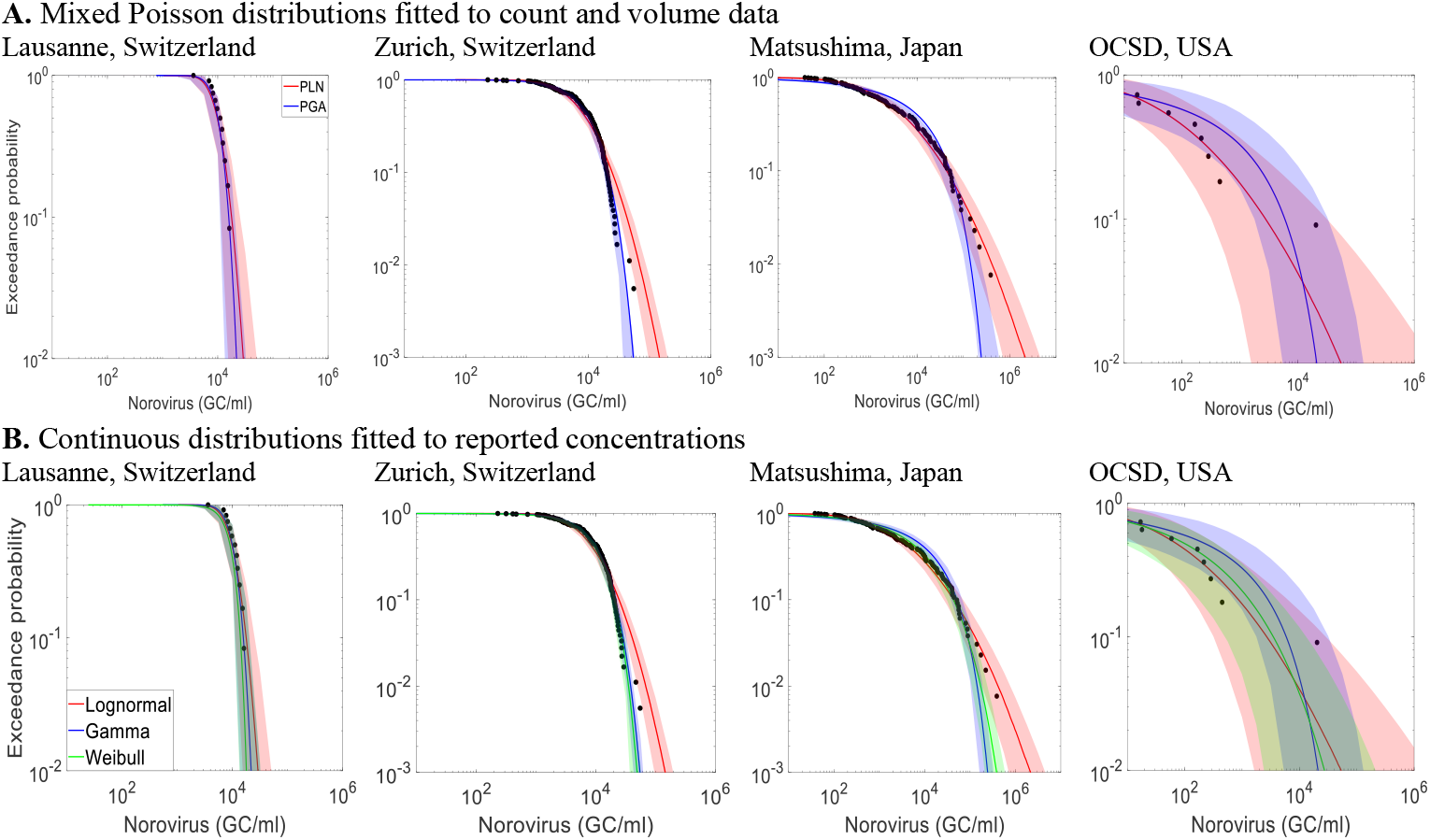
Complementary cumulative distribution functions (CCDFs) of the mixed Poisson distributions and the continuous distributions of norovirus concentrations in four wastewater treatment plants. In the upper row, red curves represent PLN distributions, and blue curves represent PGA distributions; in the lower row, red curves represent lognormal distributions, blue curves represent gamma distributions, and green curves represent Weibull distributions.

In Panel A, the PLN and PGA distributions are compared across norovirus datasets. The PLN distribution closely fits concentrations across all locations. In contrast, the PGA distribution underestimates peak concentrations in the Japan (Matsushima) and USA (OCSD) datasets, though it performs well for the Lausanne and Zürich datasets. The USA (OCSD) dataset highlights a limitation of the PGA when an extreme outlier are present—in this case, a maximum measured concentration 1.8-log10 higher than the second-highest value. The fitting process forces the gamma distribution to adjust its parameters to accommodate this extreme value. However, due to the thinner tail of the gamma compared to the lognormal, this adjustment distorts the overall fit, causing deviations in how the model represents lower concentrations.

Panel B shows the CCDFs of three continuous distribution lognormal, gamma, and Weibull distributions fitted to reported concentrations. As for the mixed Poisson distributions, the lognormal distribution generally provides a good fit across all datasets, while the gamma and Weibull distributions underestimate peak concentrations in Japan and the USA.

Those visual comparisons are supported by statistical comparisons using information criteria (Tables S1 and S2). The PLN and lognormal distributions yield lower DICm and AIC values, respectively, indicating better overall performance than PGA, gamma, and Weibull distributions.

### 3.3 Impact of sample-specific recovery rates on distributional form

Figure 3 illustrates recovery rate distributions and CCDFs of virus concentrations (adenovirus, norovirus, and enterovirus) at selected USA WWTPs, with and without sample-specific recovery corrections (using MS2 or PhiX174 as controls). The CCDFs highlight how recovery corrections impact the variability and uncertainty in predicted virus concentrations. The full set of recovery-adjusted CCDFs is presented in Figure S5.

**Fig. 3.**
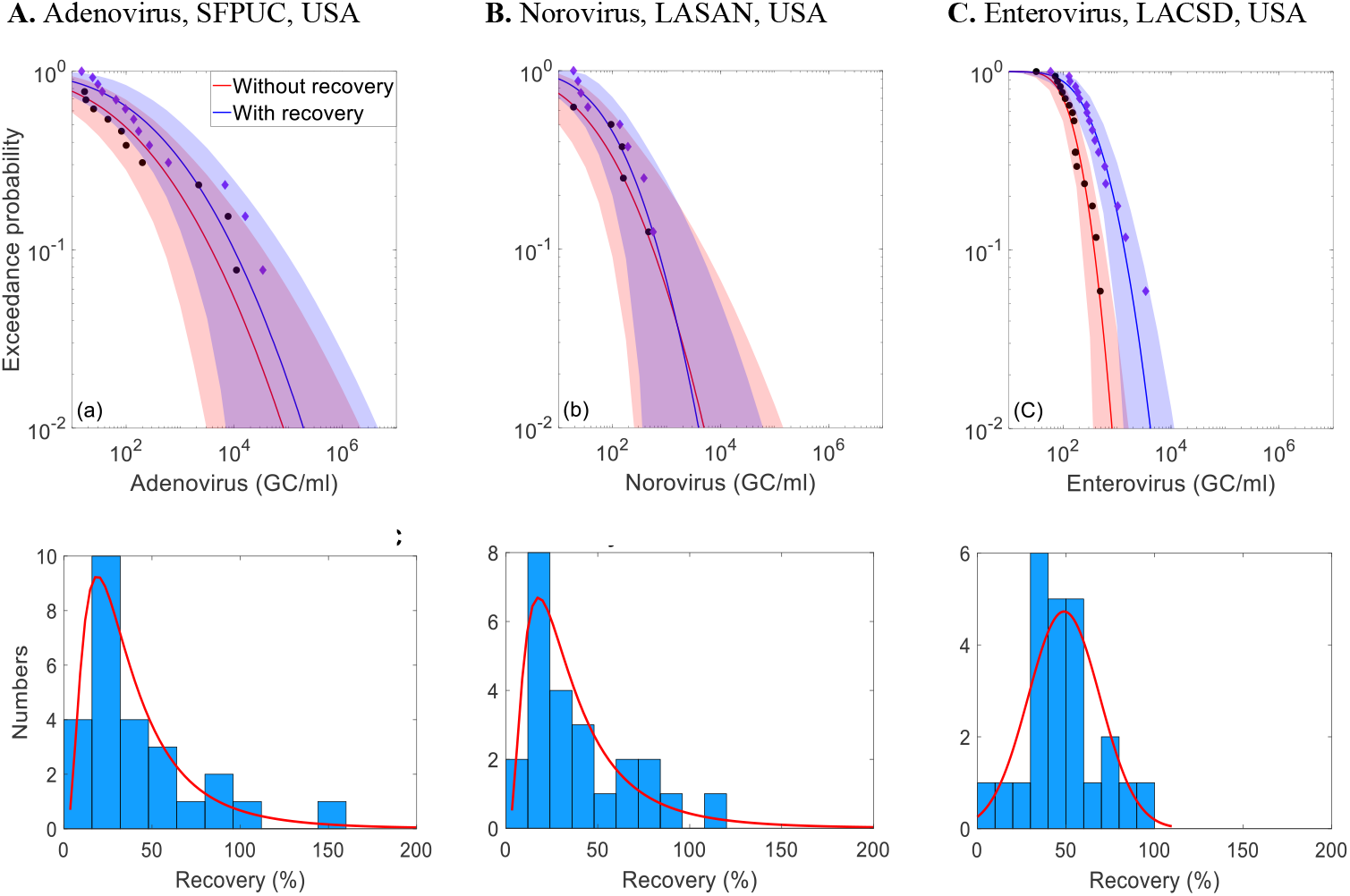
Complementary cumulative distribution functions (CCDFs) of enteric virus concentrations with and without correcting for sample-specific recovery rates for three wastewater treatment plants (WWTPs) from California, USA. Histograms illustrate the average analytical recovery rates of MS2 and PhiX174 measured in wastewater samples collected for enteric virus measurements for the three WWTPs from California, USA.

The impact of recovery rate correction on concentration distributions can depend on the distribution of the recovery data. For LACSD, where the recovery rate distribution is symmetric, with a mode of 50% and a mean of 48%, applying the recovery correction results in a horizontal shift in the CCDF, increasing concentrations without changing variability. The SFPUC and LASAN exhibit right-skewed recovery rate distributions, with modes around 20% and means around 40%. For SFPUC, recovery adjustment also leads to a horizontal shift in the upper tail of the distribution without affecting variability and uncertainty. However, for LASAN, the recovery correction does not produce a horizontal shift in the upper tail; instead, it reduces variability and uncertainty by compensating for previously underestimated low concentrations.

For the Zurich data set, recovery rates estimated using MHV process control data (Fig. 4) had a mean of 6%, ranging from 2% to 20%. These low recovery rates, when used to adjust enteric virus concentrations, increased the estimated concentration by up to two orders of magnitude. Applying the MHV recovery correction also results in a horizontal shift in the CCDF, increasing concentrations without changing variability.

**Fig. 4.**
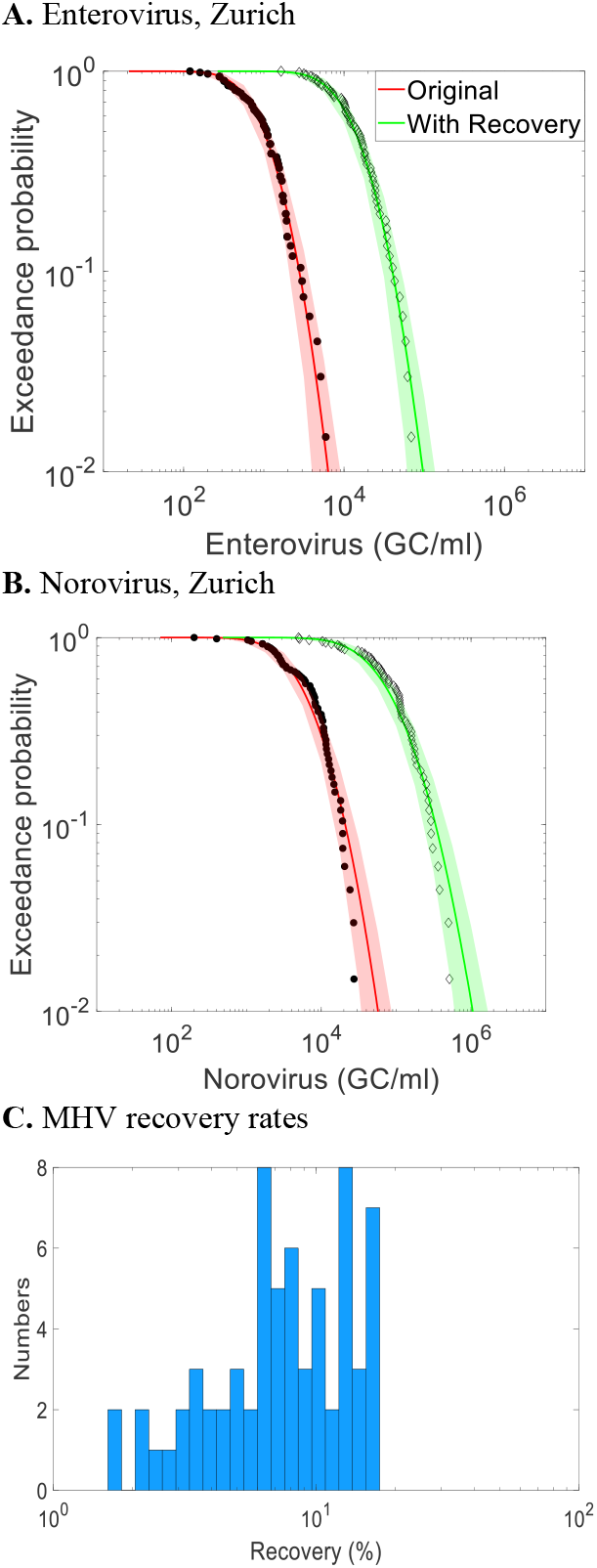
Complementary cumulative distribution functions (CCDFs) of enterovirus and norovirus with raw measurement data (red curves) for ARA Werdhölzli WWTP in Zurich, sample-specific correction with sample-specific correction with MHV recovery rates (green curves). Histogram of analytical recovery rates of MHV process control data measured in wastewater samples collected for enteric virus measurements at this WWTP.

### 3.4 Influence of virus monitoring frequency on distributional fit

Figure 5 presents CCDFs of enterovirus and norovirus concentrations from sub-sampled data at the ARA Werdhölzli WWTP in Zurich, with monitoring frequencies set to monthly, every two weeks, and weekly. Increasing the monitoring frequency from monthly to every two weeks substantially reduces 95% uncertainty intervals for both viruses. This reduction results in an accurate and less uncertain prediction of norovirus peak concentrations but an underestimation of enterovirus peak concentrations. Further increasing the frequency from bi-weekly to weekly provides a minor additional reduction in uncertainty but peak concentrations were accurately predicted for both viruses. It can also be noted that, as sampling frequency increases, the PLN and PGA distributions converge in their ability to predict the variability.

**Fig. 5.**
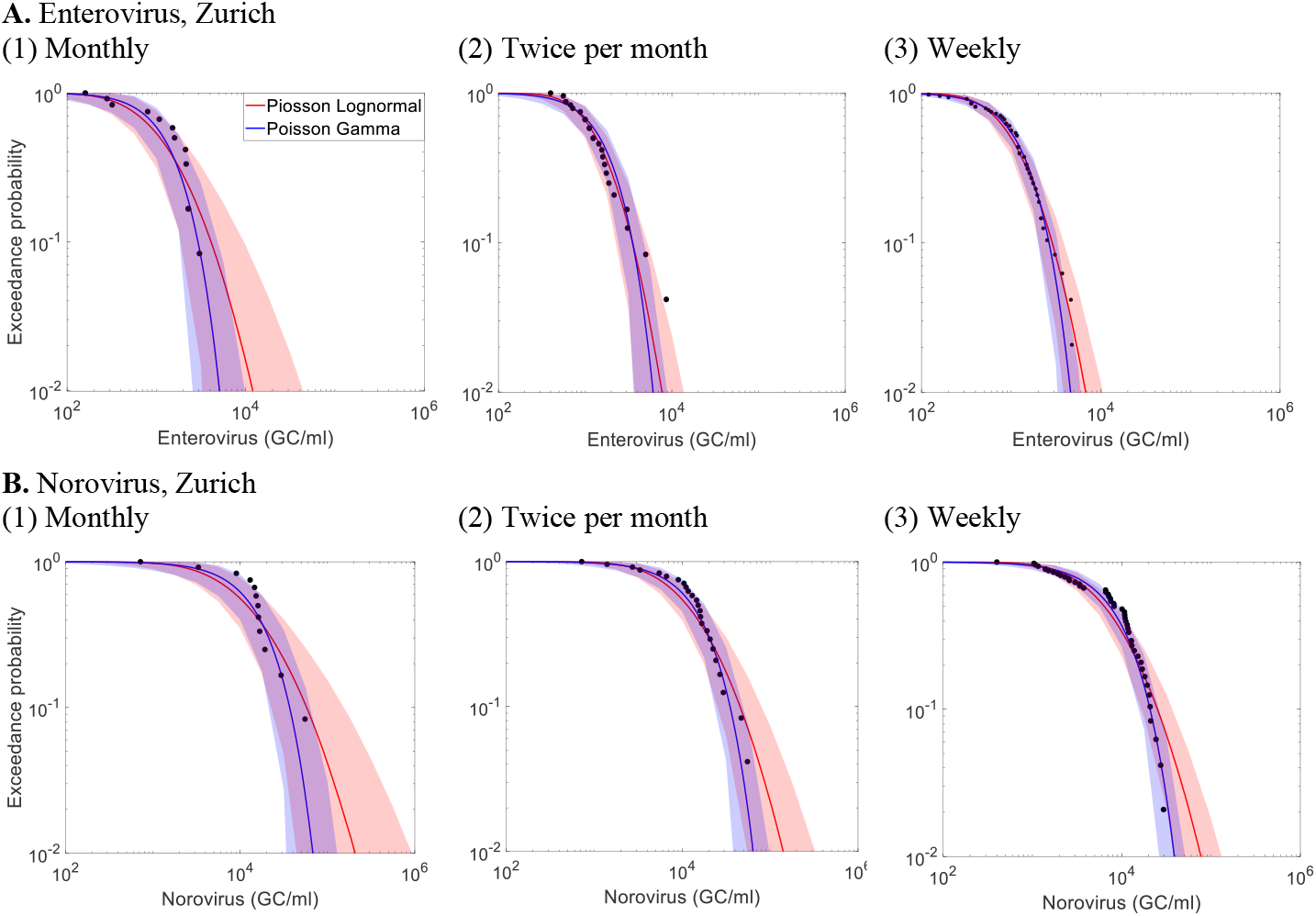
Complementary cumulative distribution function (CCDFs) of sub-sampled (top) enterovirus and (bottom) norovirus measurement data from ARA Werdhölzli WWTP in Zurich, Switzerland.

Figure 6 illustrates the relationship between sample size (i.e., the number of samples collected at a specific monitoring frequency over a set duration) and the uncertainty of the arithmetic mean, represented by the width of the 95% CI in log scale, for different standard deviation (*σ*) of the lognormal distribution. At a *σ* of 1.5, approximately 16 samples are required to reduce the 95% CI width on the arithmetic mean to below 1.0 log10. For the Zurich WWTP dataset, where *σ* is 0.97 and the sample size is 180, the 95% CI width of the mean is well below 0.3 log10. The values of *σ* for other WWTPs range from 0.7 to 2.6 (Table 2), reflecting varying levels of concentration variability across sites. This indicates that WWTPs with higher *σ* values, reflecting greater variability in virus concentrations, require more frequent sampling to achieve a precise mean estimate. For sites with lower *σ* values, less frequent sampling can still provide reliable estimates, optimizing monitoring resources.

**Fig. 6.**
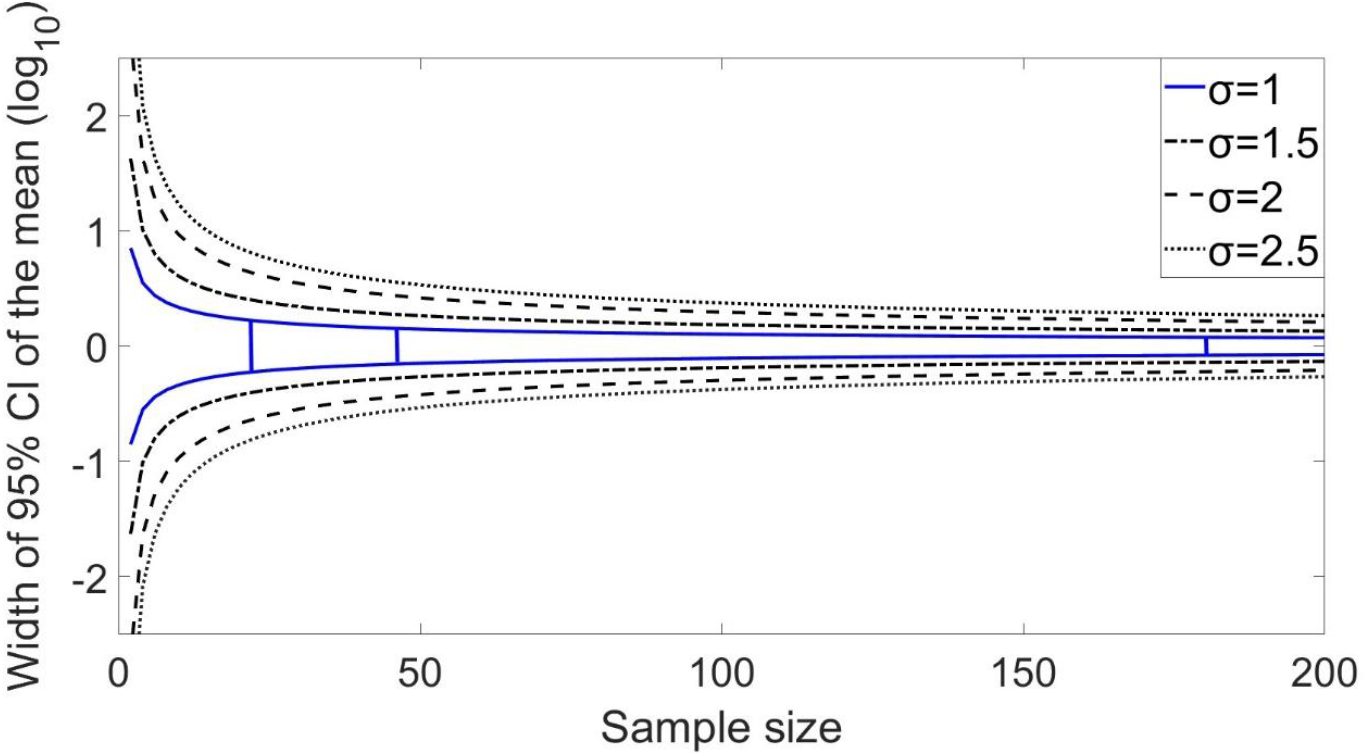
Relationship between sample size (i.e., the number of samples collected at a specific monitoring frequency for a specific duration and the width of 95% the confidence interval of the arithmetic mean virus concentration in log_10_ scale for distinct values of the *σ* parameter of the lognormal distribution. The blue curves represent the relationship for the *σ* value of 1.0 estimated for norovirus and enterovirus at the ARA Werdhölzli WWTP in Zurich. The vertical lines show the actual sample size (*n*=180), a sample size four times smaller (*n*=45, similar to weekly monitoring), and a sample size eight times smaller (*n*=23; similar to monitoring twice a month).

## 4 Discussion

In this study, we assessed the temporal variability of enteric virus concentrations in influent wastewater from multiple WWTPs across Switzerland, the USA, and Japan. The data set and associated models support models reliant on estimates of distributions of pathogen concentrations in wastewater influent, for example exposure assessment models within a QMRA framework. By evaluating original and literature-derived datasets, we observed clear temporal patterns in norovirus concentrations, with fluctuations ranging up to five orders of magnitude over a year. An apparent increase in norovirus was present in winter months in Zurich, Switzerland, and Matsushima, Japan, consistent with findings from other studies (Farkas et al., 2018; Prevost et al., 2015). While adenovirus and rotavirus also peaked in winter in Lausanne, Switzerland, and the USA, more extensive datasets would be necessary to investigate seasonality at these locations. Our study revealed that although fluctuations in enteric virus concentrations may appear sporadic in time series analyses, they can be governed by common underlying generative processes described with simple parametric distributions. These distributions must be carefully chosen to accurately predict peak concentrations, as these peaks play a critical role in applications like QMRA (Smeets, 2010; Sylvestre, 2020).

Our findings demonstrate that the lognormal distribution is well-suited for modelling the variability of enteric virus concentration across different locations and virus types. Posterior predictive checks and model comparison with deviance information criteria supported this selection. In contrast, gamma and Weibull distributions often fail to capture the high variability observed, especially at CV values above 2.5. This limitation is expected because these distributions lack the necessary skewness to model extreme variability, as their skewness plateaus with increasing CV, unlike the lognormal distribution (Vargo et al., 2017).

The prevalence of the lognormal distribution indicates that multiplicative processes shape the variability in virus concentrations and suggests the absence of strict constraints (e.g., physical or biological limits) on maximum virus concentrations under monitored conditions. The variability likely arises from the way enteric viruses are shed by infected people within communities. Enteric virus densities in the stools of infected individuals vary widely; for example, ranging from 10^5^ to 10^11^ genome copies per gram for norovirus (Atmar et al., 2008; Teunis et al., 2015). When aggregated, these highly variable virus loads result in wastewater concentrations that follow lognormal distributions. The upper tail of the distribution may be shaped by rare individuals or clusters shedding exceptionally high virus loads. Localized outbreaks may further impact the upper tail by causing sudden increases in virus shedding in communities. Peak concentrations may be amplified in smaller WWTPs with lower dilution capacities. In our study, norovirus peaks were higher at the WWTP in Matsushima, Japan (serving ∼9,600 people) compared to the ARA Werdhölzli WWTP in Zurich, Switzerland (serving ∼471,000 people). It is worth highlighting that not all microorganisms in wastewater follow a lognormal distribution. The concentrations of *Escherichia coli* in influent wastewater from Swiss WWTPs follow a gamma distribution (Conforti et al., 2024). This distinction is likely due to the lower variability of *E. coli* densities in stools, which typically range from 10^6^ to 10^9^ colony-forming units (CFU) per gram (Harwood et al., 2019), combined with relatively constant shedding rates.

Accurately estimating the distribution of virus concentrations requires acknowledging that observations are subject to various measurement uncertainties, including sampling uncertainty, detection limits, and analytical recovery rates. Discrete mixed Poisson distributions are better candidate models than continuous distributions, as they can incorporate uncertainty associated with non-detects into parametric analyses (Chik et al., 2018). In addition, discrete mixture distributions—which directly model virus counts within a sample—could be further developed to address uncertainties related to virus aggregation in wastewater.

When adjusting concentrations for recovery, the reliability of surrogate viruses depends on how closely their morphology and behavior in wastewater align with those of the target virus. In our study, the model enveloped virus MHV and the non-enveloped bacteriophages MS2 and PhiX174 had variable recovery estimates, ranging from 1 to 100%. On average, MHV recovery rates were around 1.0-log10 lower than those of MS2 and PhiX174, likely due to the enveloped structure of MHV. Enveloped viruses have unique adsorption, aggregation, and recovery behaviors in raw wastewater, with studies indicating that they are more strongly associated with solids and are more susceptible to inactivation during recovery processes compared to non-enveloped viruses (Ye et al., 2016). To improve surrogate selection, evaluating correlations between the recovery rates of these surrogates and those of naturally occurring enteric viruses could provide insights into which surrogates are most reliable for assessing virus concentration distributions.

Our analysis of high-frequency monitoring data from Zurich WWTP demonstrated that increasing the virus monitoring frequency from monthly to bi-weekly significantly reduced the uncertainty of the modelled distribution of concentrations. Further increasing to weekly sampling, however, provided minimal additional reductions in uncertainty but improved the prediction of peaks for enterovirus. For lognormally distributed concentrations, the optimal monitoring frequency varies based on the standard deviation of the natural logarithm of the

concentration (*σ*). For the Zurich WWTP, *σ* is approximately 1.0 for norovirus and enterovirus, but for other WWTPs in this study, *σ* values up to 2.6 were found. In these cases, more frequent monitoring may be needed to adequately estimate the distribution for QMRA. Developing adaptive monitoring strategies that start with high-frequency sampling (e.g., weekly) to assess concentration variability (*σ*) and then adjusting the frequency as the estimated *σ* stabilizes could make monitoring efforts more cost-efficient and tailored to site-specific conditions.

Our findings highlight that site-specific differences in mean concentrations, variability, and distribution shapes of enteric viruses can be substantial. Local shedding patterns, population size, or outbreak dynamics may shape the underlying generative processes at each site. At the same time, methodological factors, such as sampling intervals, sample volumes, and analytical recovery rates, also influence measured concentrations. Although all viruses here were measured using PCR-based methods with process controls and recovery spikes, caution is warranted when comparing results across different WWTPs, as these factors may also impact reported variability. This underscores the importance of carefully modelling site-specific distributions, which can complicate meta-analysis. Although pooling data from multiple sites into one single distribution, as proposed by Darby et al. (2023), can offer insights for developing risk-based pathogen treatment requirements in potable reuse; this approach can mask site-specific uncertainty by treating variability as random noise around a single mean. Consequently, sites with distinctly higher (or lower) concentrations may be missed, and peak concentrations, often the key driver in risk assessment, may be underestimated. Random-effects meta-analysis addresses this limitation by allowing each data set to have its own distribution nested within a broader population distribution, preserving site-specific variability (Higgins et al., 2009). Further development of meta-analytic models for microbial risk assessment would help refine how we estimate and manage virus concentration variability across diverse WWTPs.

## 5 Conclusions

Based on our analysis of enteric virus datasets from eight WWTPs across Switzerland, Japan, and the USA, we conclude:

- Enteric virus concentrations in wastewater influent vary widely by virus type and location, with average norovirus concentrations ranging from 1.1 × 10^2^ to 1.8 × 10^4^ genome copies per liter across sites, emphasizing the need to preserve site-specific variability rather than relying solely on aggregated estimates.
- The lognormal distribution accurately predicted peak virus concentrations and outperformed gamma and Weibull distributions.
- Recovery-corrected concentrations using MS2, PhiX174, and MHV matrix spike data were successfully incorporated into parametric models.
- Weekly over a year should be sufficient to estimate the annual average concentration within a 95% confidence interval of 0.5-log10 at most sites. High variability sites (*σ* > 2) may need more frequent monitoring to achieve accurate estimates of distributions.

## 6 Data statement

Enteric viruses, MHV data as well as R codes will be made available on Zenodo upon acceptance of the manuscript.

## 7 Acknowledgements

This work was supported by the Swiss National Science Foundation (grant no. 31003A_182468). Data from ARA Werdhölzli in Zurich, Switzerland, were obtained with support from the Swiss Federal Office of Public Health. ES was funded by postdoctoral fellowships from the Natural Sciences and Engineering Research Council of Canada (558161-2021) and the Fonds de Recherche du Québec Nature et Technologies (303866). Thanks to Franziska Böni, Xavier Fernandez-Cassi, Anina Kull, Elyse Stachler, and Blanche Wies for supporting the sample collection and processing.

## Notes

### Competing Interest Statement

The authors have declared no competing interest.

## References

Atmar, R.L., Opekun, A.R., Gilger, M.A., Estes, M.K., Crawford, S.E., Neill, F.H. and Graham, D.Y. 2008. Norwalk virus shedding after experimental human infection. Emerging infectious diseases 14(10), 1553.

Blatchley III, E.R., Gong, W.L., Alleman, J.E., Rose, J.B., Huffman, D.E., Otaki, M. and Lisle, J.T. 2007. Effects of wastewater disinfection on waterborne bacteria and viruses. Water environment research 79(1), 81–92.

Chik, A.H.S., Schmidt, P.J. and Emelko, M.B. 2018. Learning something from nothing: the critical importance of rethinking microbial non-detects. Frontiers in Microbiology 9, 2304.

Conforti, S., Holschneider, A., Sylvestre, É. and Julian, T.R. 2024. Monitoring ESBL-Escherichia coli in Swiss wastewater between November 2021 and November 2022: insights into population carriage. Msphere 9(5), e00760–00723.

Daelman, M.R., De Baets, B., van Loosdrecht, M.C. and Volcke, E.I. 2013. Influence of sampling strategies on the estimated nitrous oxide emission from wastewater treatment plants. Water research 47(9), 3120–3130.

Darby, E., Olivieri, A., Haas, C., Di Giovanni, G., Jakubowski, W., Leddy, M., Nelson, K.L., Rock, C., Slifko, T. and Pecson, B.M. 2023. Identifying and aggregating high-quality pathogen data: a new approach for potable reuse regulatory development. Environmental Science: Water Research & Technology 9(6), 1646–1653.

Farkas, K., Marshall, M., Cooper, D., McDonald, J.E., Malham, S.K., Peters, D.E., Maloney, J.D. and Jones, D.L. 2018. Seasonal and diurnal surveillance of treated and untreated wastewater for human enteric viruses. Environmental Science and Pollution Research 25, 33391–33401.

Fernandez-Cassi, X., Scheidegger, A., Bänziger, C., Cariti, F., Corzon, A.T., Ganesanandamoorthy, P., Lemaitre, J.C., Ort, C., Julian, T.R. and Kohn, T. 2021. Wastewater monitoring outperforms case numbers as a tool to track COVID-19 incidence dynamics when test positivity rates are high. Water research 200, 117252.

Gelman, A., Carlin, J.B., Stern, H.S., Dunson, D.B., Vehtari, A. and Rubin, D.B. 2013. Bayesian Data Analysis.

Higgins, J.P., Thompson, S.G. and Spiegelhalter, D.J. 2009. A re-evaluation of random-effects meta-analysis. Journal of the Royal Statistical Society Series A: Statistics in Society 172(1), 137–159.

Huisman, J.S., Scire, J., Caduff, L., Fernandez-Cassi, X., Ganesanandamoorthy, P., Kull, A., Scheidegger, A., Stachler, E., Boehm, A.B. and Hughes, B. 2022. Wastewater-based estimation of the effective reproductive number of SARS-CoV-2. Environmental Health Perspectives 130(5), 057011.

Kageyama, T., Kojima, S., Shinohara, M., Uchida, K., Fukushi, S., Hoshino, F.B., Takeda, N. and Katayama, K. 2003. Broadly reactive and highly sensitive assay for Norwalk-like viruses based on real-time quantitative reverse transcription-PCR. Journal of clinical microbiology 41(4), 1548–1557.

Kazama, S., Miura, T., Masago, Y., Konta, Y., Tohma, K., Manaka, T., Liu, X., Nakayama, D., Tanno, T. and Saito, M. 2017. Environmental surveillance of norovirus genogroups I and II for sensitive detection of epidemic variants. Applied and environmental microbiology 83(9), e03406–03416.

Kitajima, M., Iker, B.C., Pepper, I.L. and Gerba, C.P. 2014. Relative abundance and treatment reduction of viruses during wastewater treatment processes—identification of potential viral indicators. Science of the Total Environment 488, 290–296.

La Rosa, G. and Muscillo, M. (2013) Viruses in food and water, pp.97–125, Elsevier.

Li, C., Sylvestre, É., Fernandez-Cassi, X., Julian, T.R. and Kohn, T. 2023. Waterborne virus transport and the associated risks in a large lake. Water research 229, 119437.

Li, D., Shi, H.-c. and Jiang, S.C. 2010. Concentration of viruses from environmental waters using nanoalumina fiber filters. Journal of Microbiological Methods 81(1), 33–38.

Loisy, F., Atmar, R., Guillon, P., Le Cann, P., Pommepuy, M. and Le Guyader, F. 2005. Real-time RT-PCR for norovirus screening in shellfish. Journal of virological methods 123(1), 1–7.

Monpoeho, S., Dehee, A., Mignotte, B., Schwartzbrod, L., Marechal, V., Nicolas, J.-C., Billaudel, S. and Ferre, V. 2000. Quantification of enterovirus RNA in sludge samples using single tube real-time RT-PCR. Biotechniques 29(1), 88–93.

Nadeau, S., Devaux, A.J., Bagutti, C., Alt, M., Hampe, E.I., Kraus, M., Würfel, E., Koch, K.N., Fuchs, S. and Tschudin-Sutter, S. 2023. Influenza transmission dynamics quantified from wastewater. medRxiv, 2023.2001. 2023.23284894.

Olsson, U. 2005. Confidence intervals for the mean of a log-normal distribution. Journal of Statistics Education 13(1).

Pecson, B.M., Darby, E., Danielson, R., Dearborn, Y., Di Giovanni, G., Jakubowski, W., Leddy, M., Lukasik, G., Mull, B. and Nelson, K.L. 2022. Distributions of waterborne pathogens in raw wastewater based on a 14-month, multi-site monitoring campaign. Water Research 213, 118170.

Petterson, S., Grøndahl-Rosado, R., Nilsen, V., Myrmel, M. and Robertson, L.J. 2015. Variability in the recovery of a virus concentration procedure in water: implications for QMRA. Water research 87, 79–86.

Prevost, B., Lucas, F.S., Goncalves, A., Richard, F., Moulin, L. and Wurtzer, S. 2015. Large scale survey of enteric viruses in river and waste water underlines the health status of the local population. Environment international 79, 42–50.

Quintero, A. and Lesaffre, E. 2018. Comparing hierarchical models via the marginalized deviance information criterion. Statistics in Medicine 37(16), 2440–2454.

Smeets, P.W. (2010) Stochastic modelling of drinking water treatment in quantitative microbial risk assessment, IWA Publishing.

Sylvestre, É. (2020) Systematic assessment of microbial risks associated with hydrometeorological events for drinking water safety management, Ecole Polytechnique, Montreal (Canada).

Sylvestre, É., Burnet, J.-B., Smeets, P., Medema, G., Prévost, M. and Dorner, S. 2020. Can routine monitoring of E. coli fully account for peak event concentrations at drinking water intakes in agricultural and urban rivers? Water research 170, 115369.

Sylvestre, É., Prévost, M., Smeets, P., Medema, G., Burnet, J.B., Cantin, P., Villion, M., Robert, C. and Dorner, S. 2021. Importance of Distributional Forms for the Assessment of Protozoan Pathogens Concentrations in Drinking-Water Sources. Risk Analysis 41(8), 1396–1412.

Teunis, P., Sukhrie, F., Vennema, H., Bogerman, J., Beersma, M. and Koopmans, M. 2015. Shedding of norovirus in symptomatic and asymptomatic infections. Epidemiology & Infection 143(8), 1710–1717.

Tsai, Y.-L., Sobsey, M., Sangermano, L. and Palmer, C. 1993. Simple method of concentrating enteroviruses and hepatitis A virus from sewage and ocean water for rapid detection by reverse transcriptase-polymerase chain reaction. Applied and Environmental Microbiology 59(10), 3488–3491.

Vargo, E., Pasupathy, R. and Leemis, L.M. 2017. Moment-ratio diagrams for univariate distributions. Computational Probability Applications, 149–164.

Xagoraraki, I., Yin, Z. and Svambayev, Z. 2014. Fate of viruses in water systems. Journal of Environmental Engineering 140(7), 04014020.

Ye, Y., Ellenberg, R.M., Graham, K.E. and Wigginton, K.R. 2016. Survivability, partitioning, and recovery of enveloped viruses in untreated municipal wastewater. Environmental science & technology 50(10), 5077–5085.

Zhang, C.-M., Xu, L.-M., Xu, P.-C. and Wang, X.C. 2016. Elimination of viruses from domestic wastewater: requirements and technologies. World Journal of Microbiology and Biotechnology 32, 1–9.

