## Supplementary material for "Lognormal distributions capture site-specific variability in enteric virus concentrations in wastewater": Supplementary_information_March2025_Preprint.docx

**Supplementary Information Contents:**

**S1 Quality control for the RT-dPCR assay**

Quality control results for the enterovirus and norovirus duplex RT-dPCR assay.

**S2 Additional results**

Additional figures and tables not included in the main paper.

**S1 Quality control for the EV / NoV duplex RT-dPCR assay**

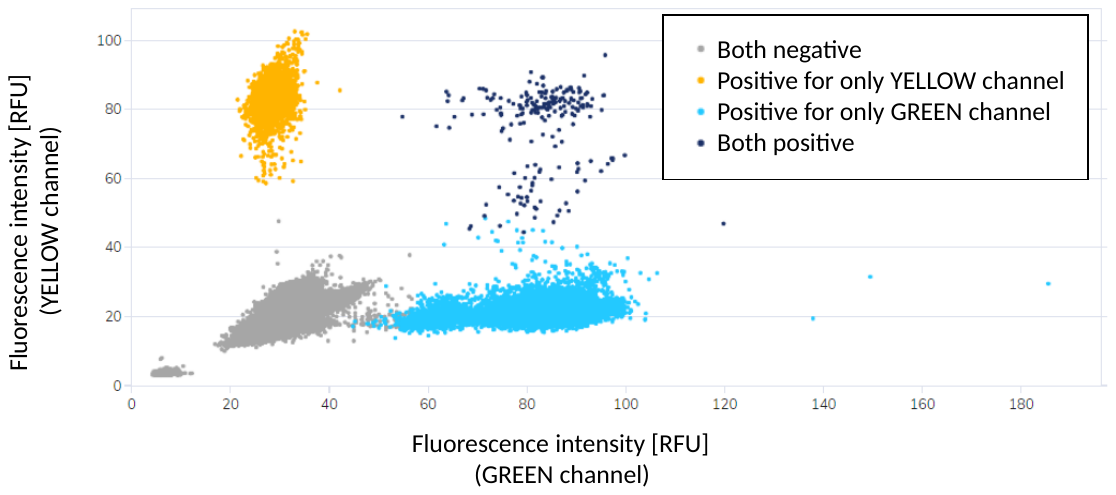

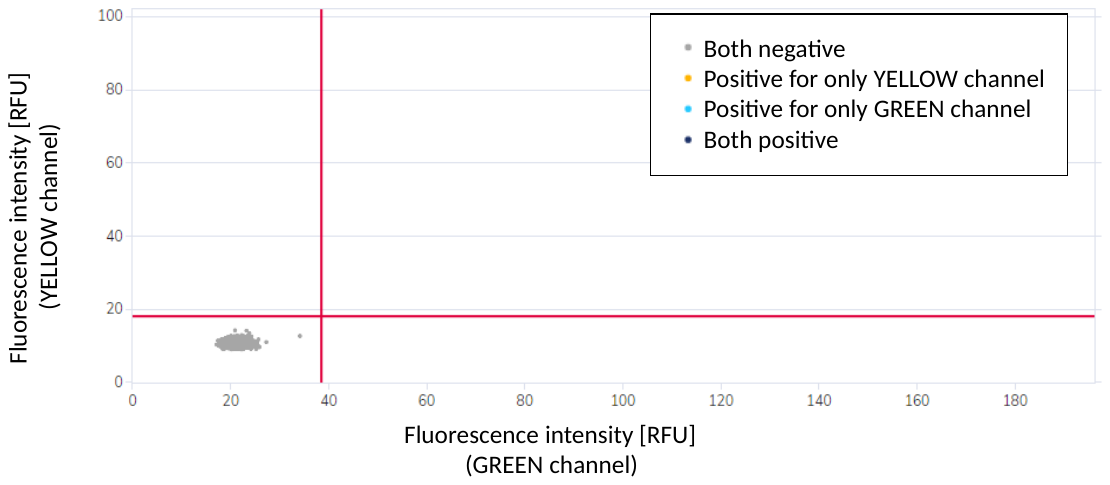

**Fig. S1**. Example fluorescence plots from the EV / NoV GII duplex RT-dPCR assay measured on the QIACuity One 2-plex device. EV is measured in the yellow channel, NoV GII is measured in the green channel. The plots include analysed wastewater samples collected between February 1 and May 2, 2021 (top) and those of negative controls (bottom). Red lines indicate automatically generated thresholds.

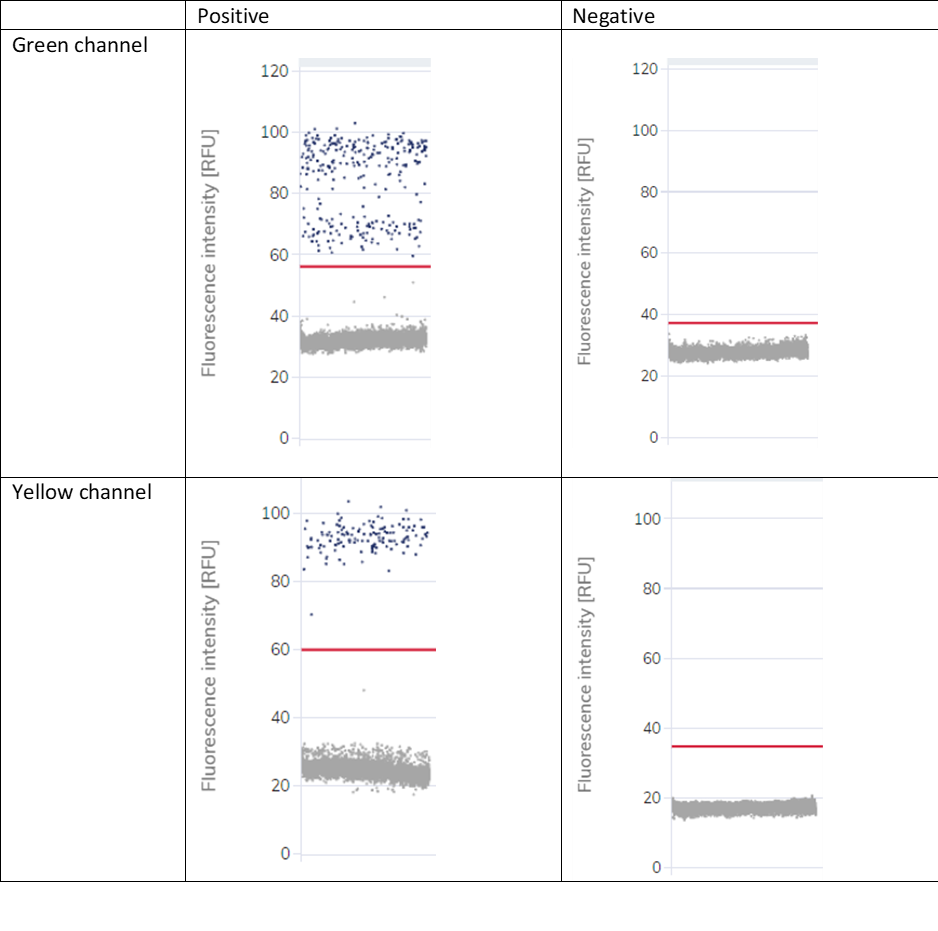

**Fig. S2**. Example outputs generated by the QIAcuity One 2-plex Device for a sample containing both EV (yellow channel) and NoV GII (green channel) and a negative control. Red lines show automatically generated thresholds between positive and negative partitions.

**S2 Additional results**

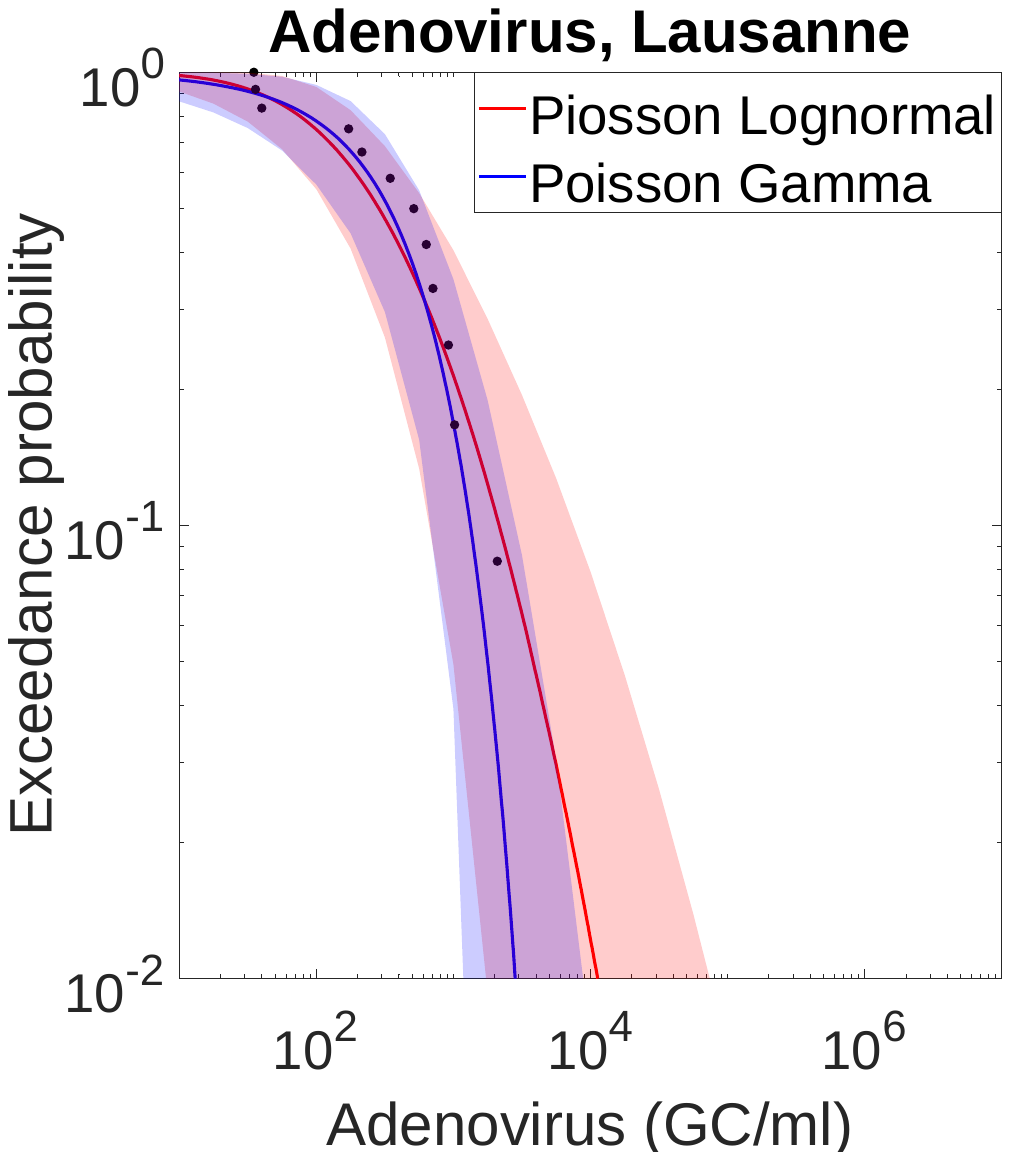

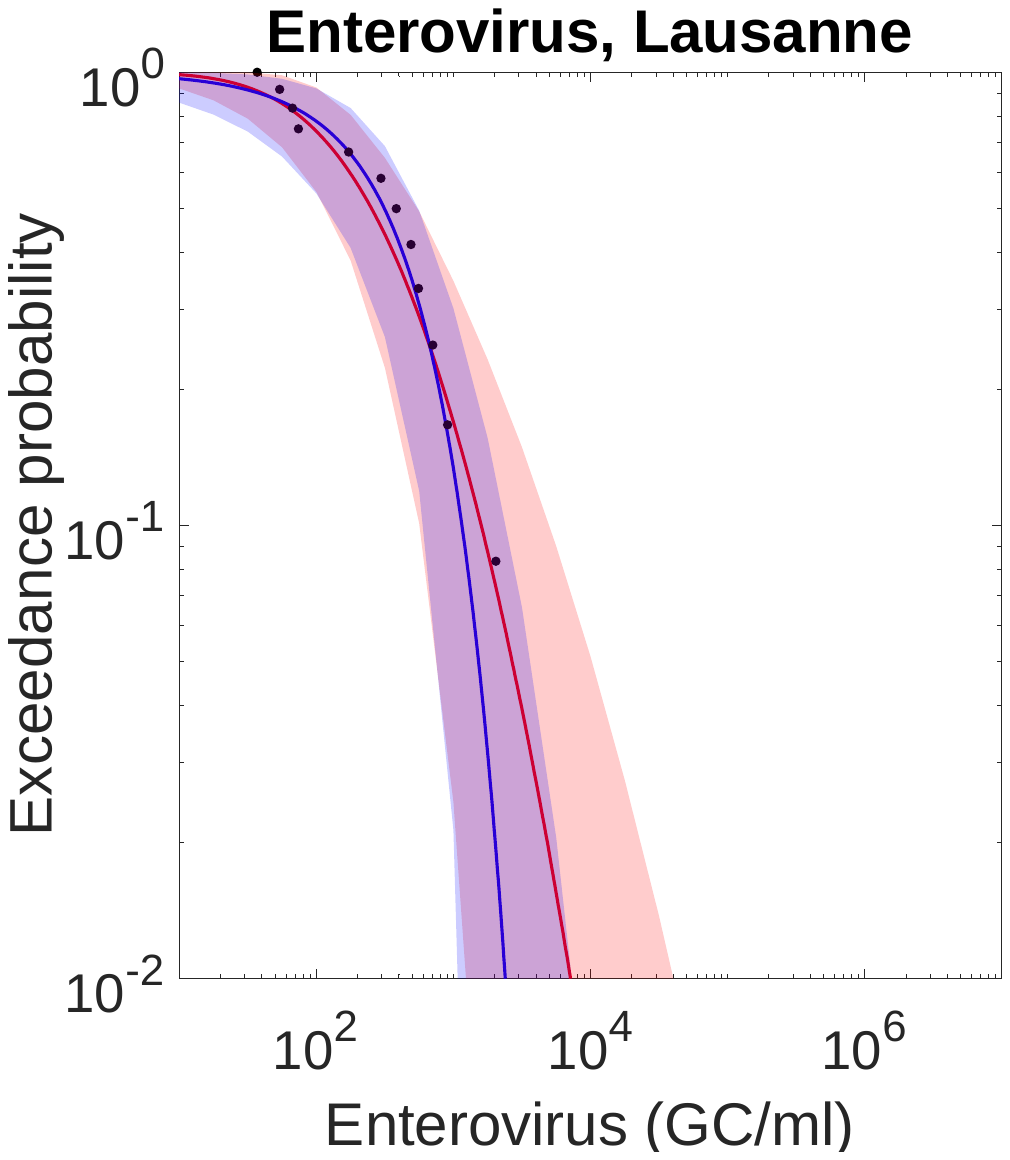

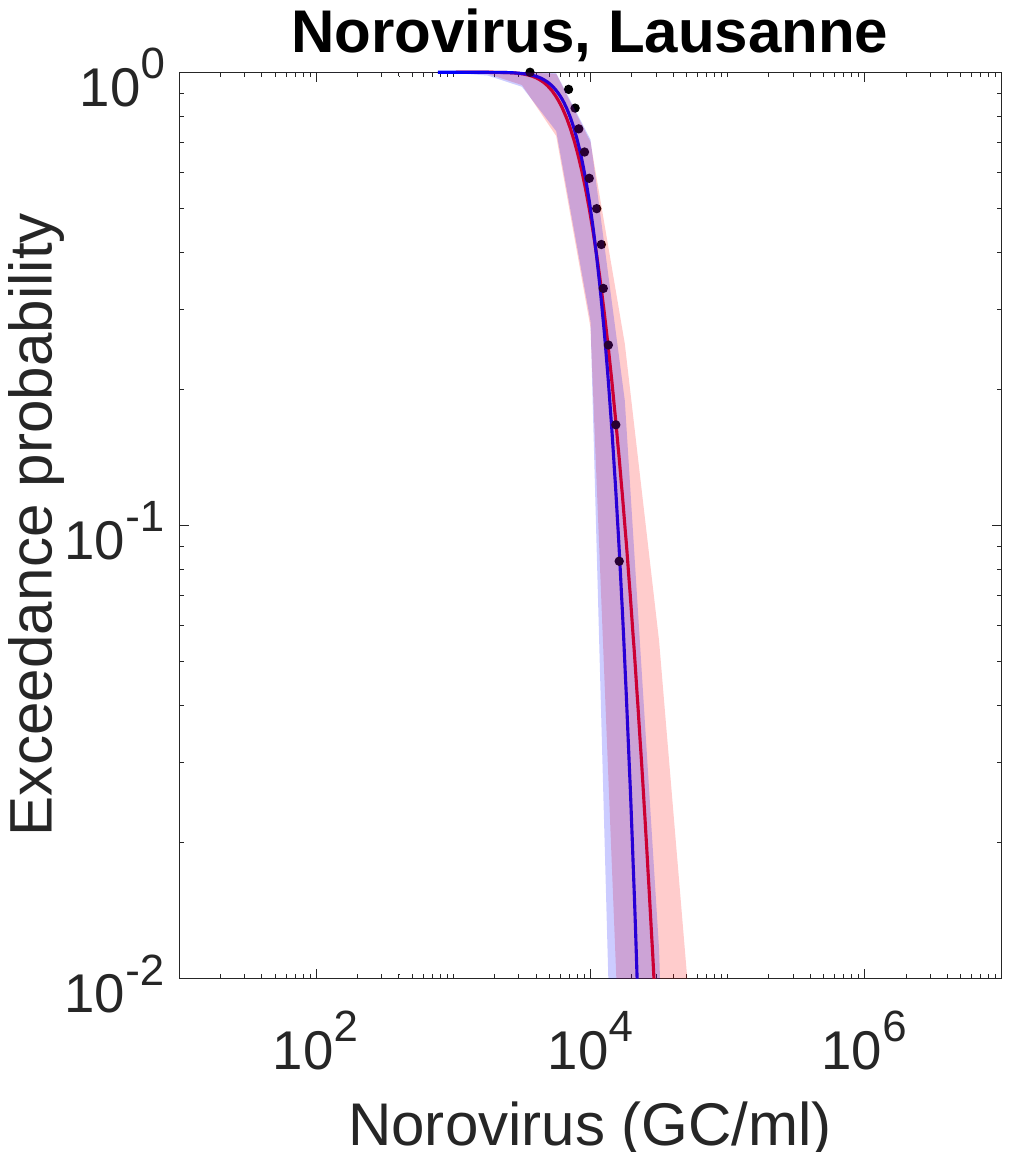

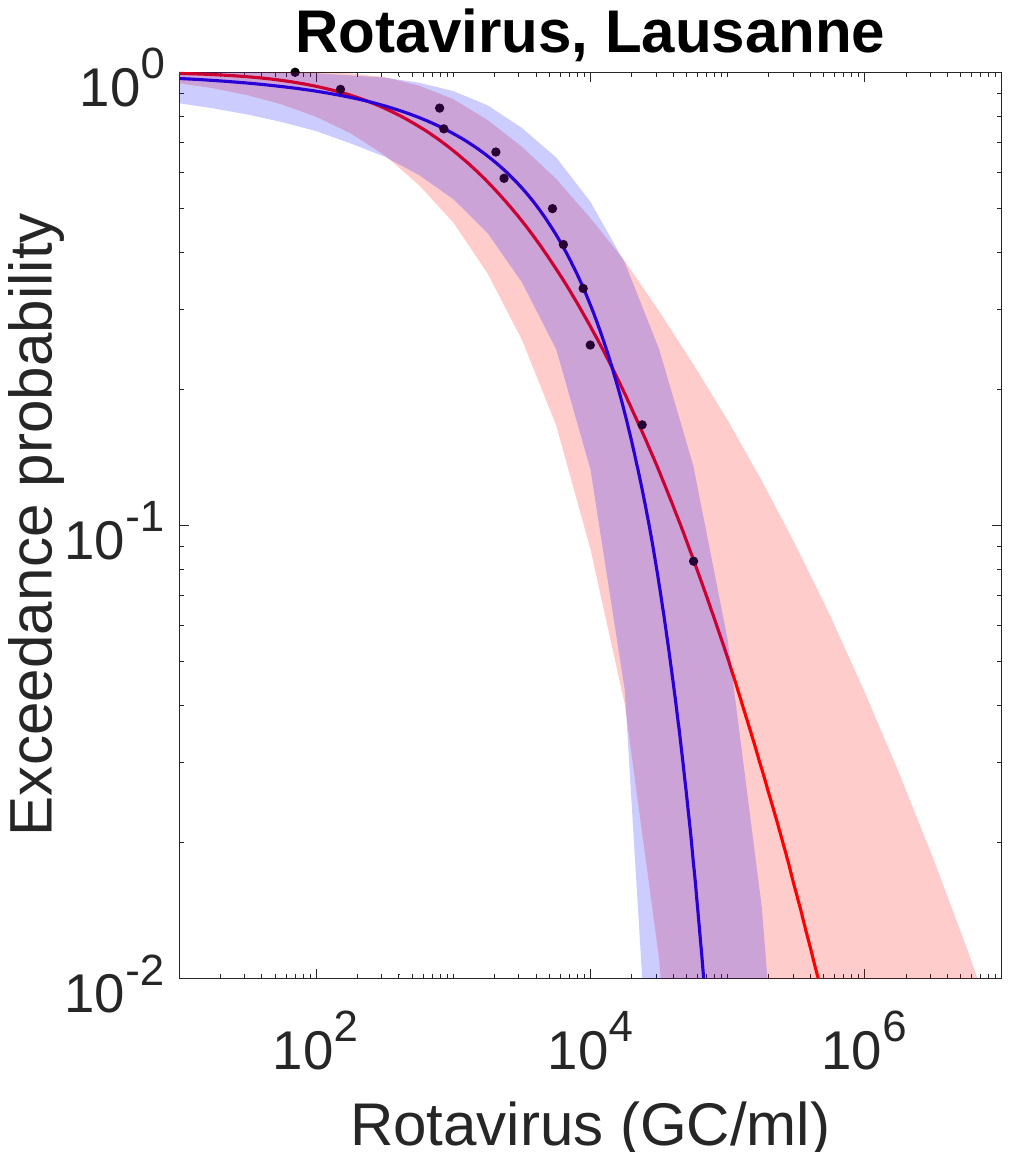

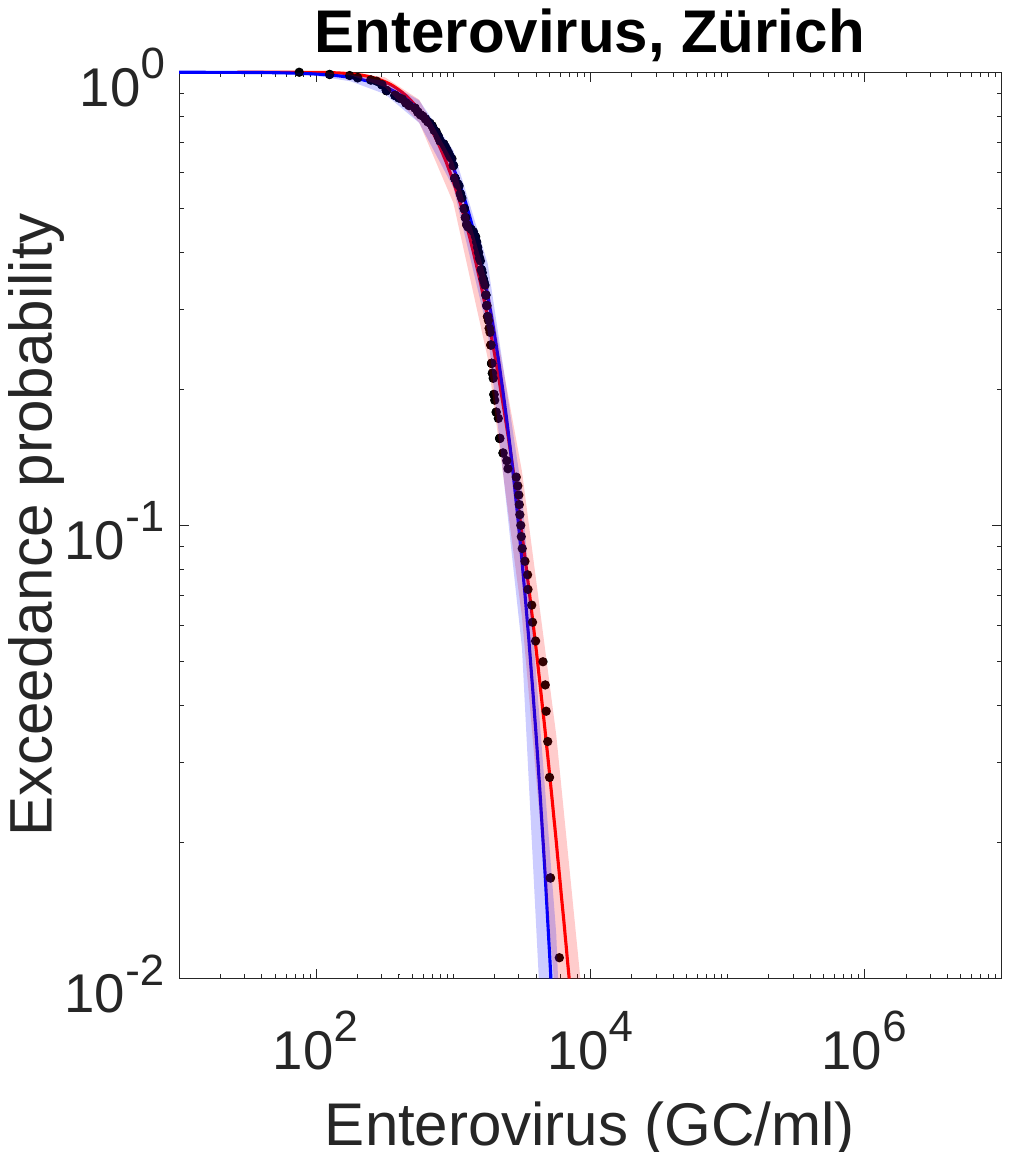

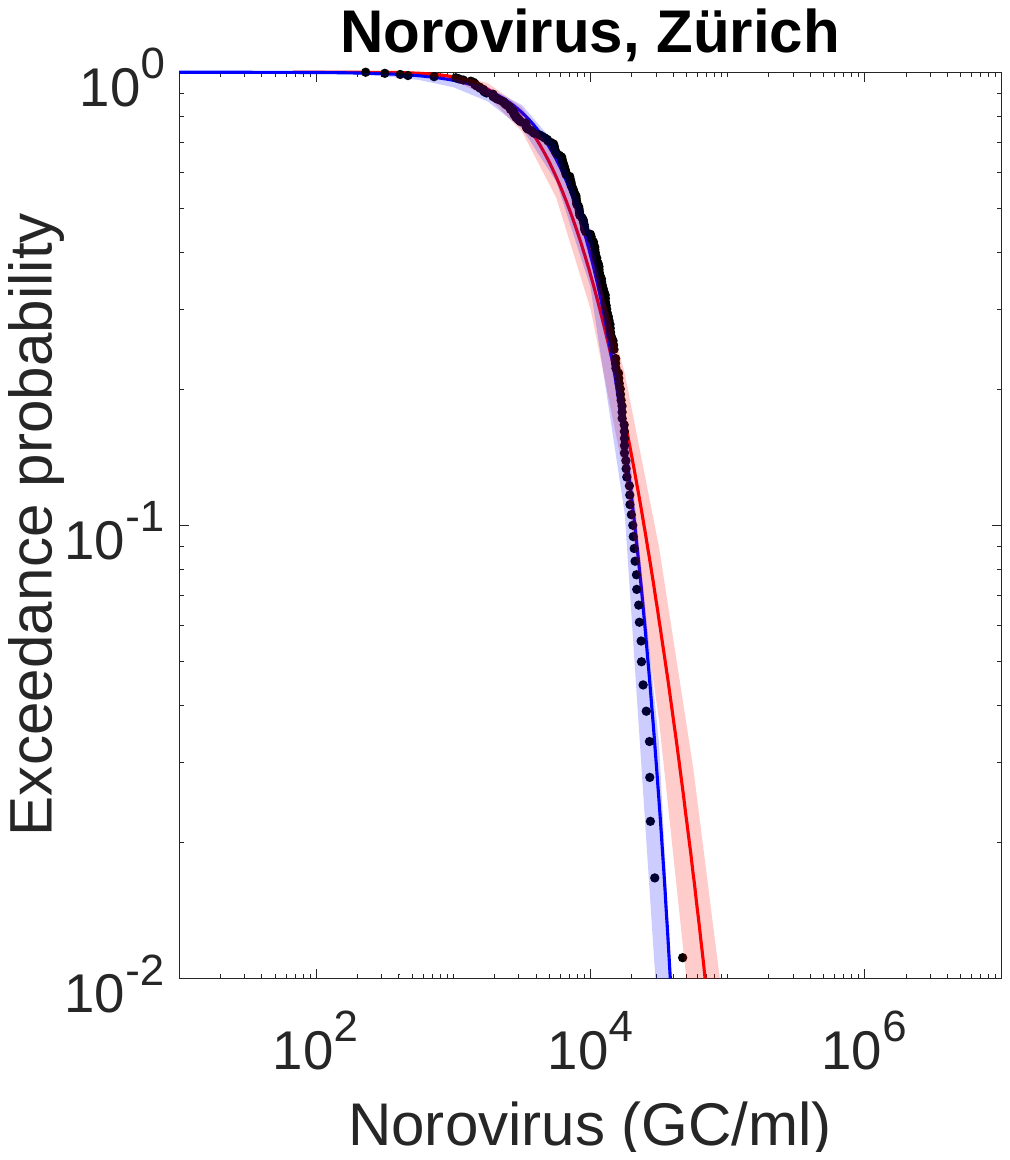

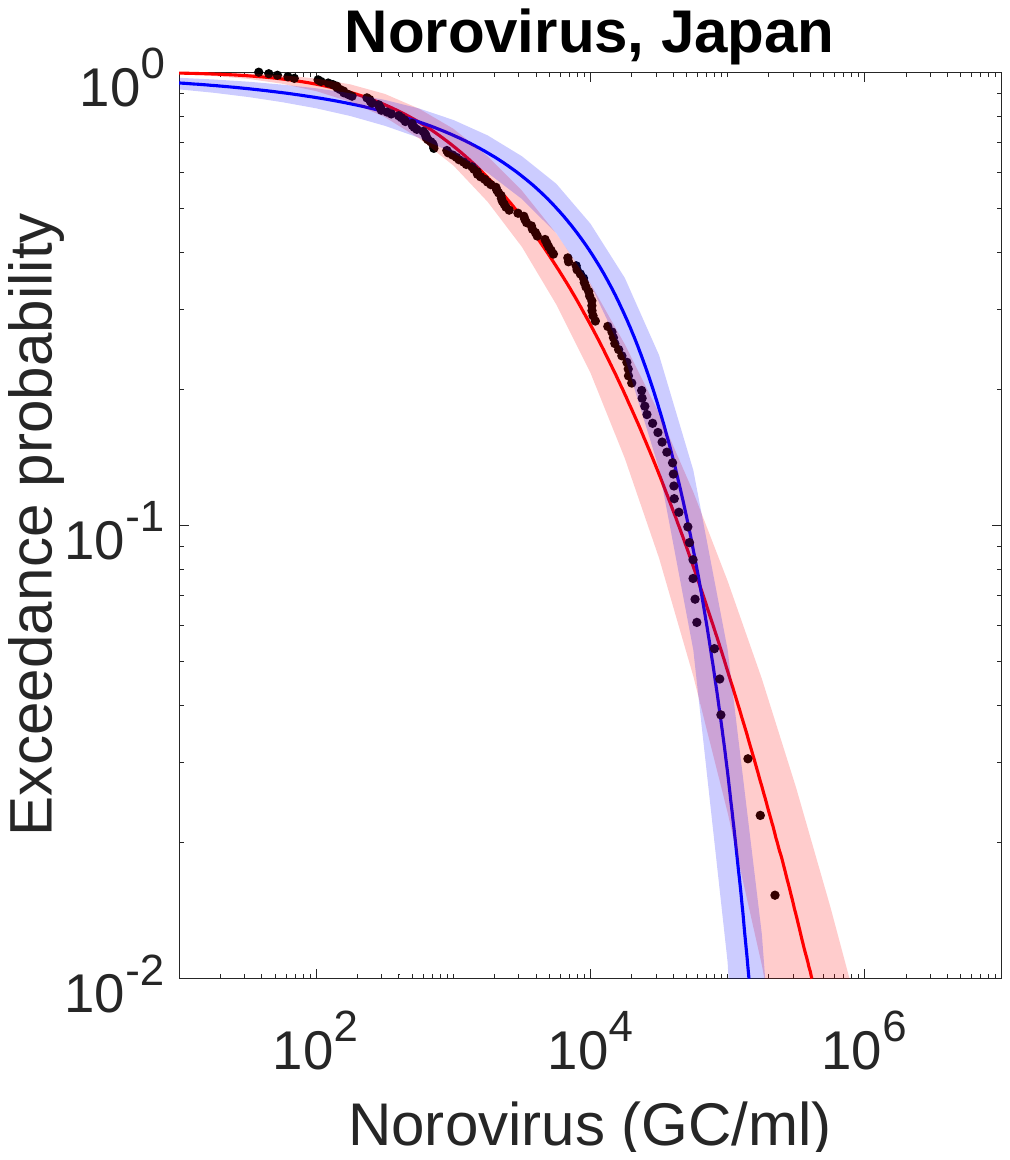

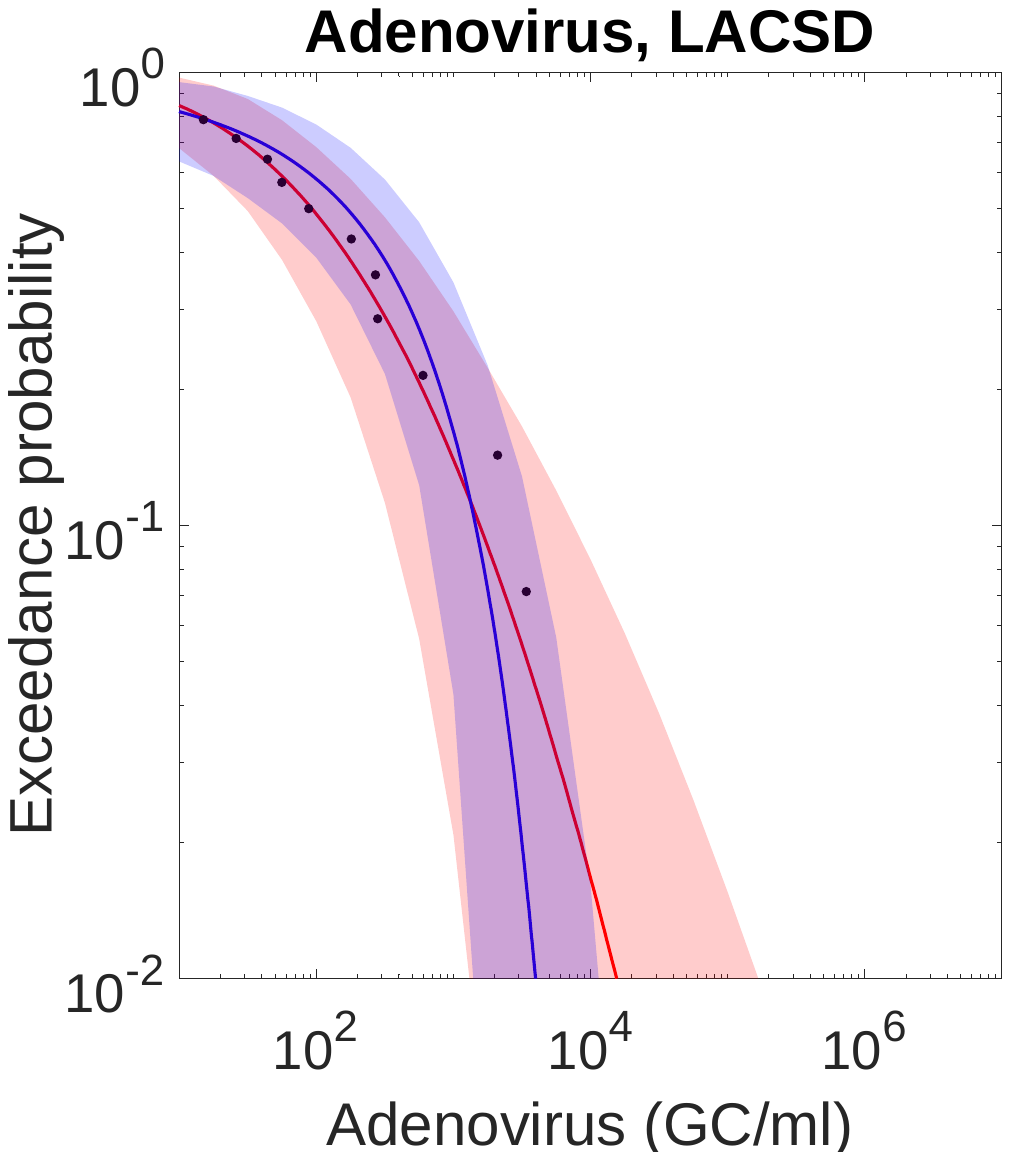

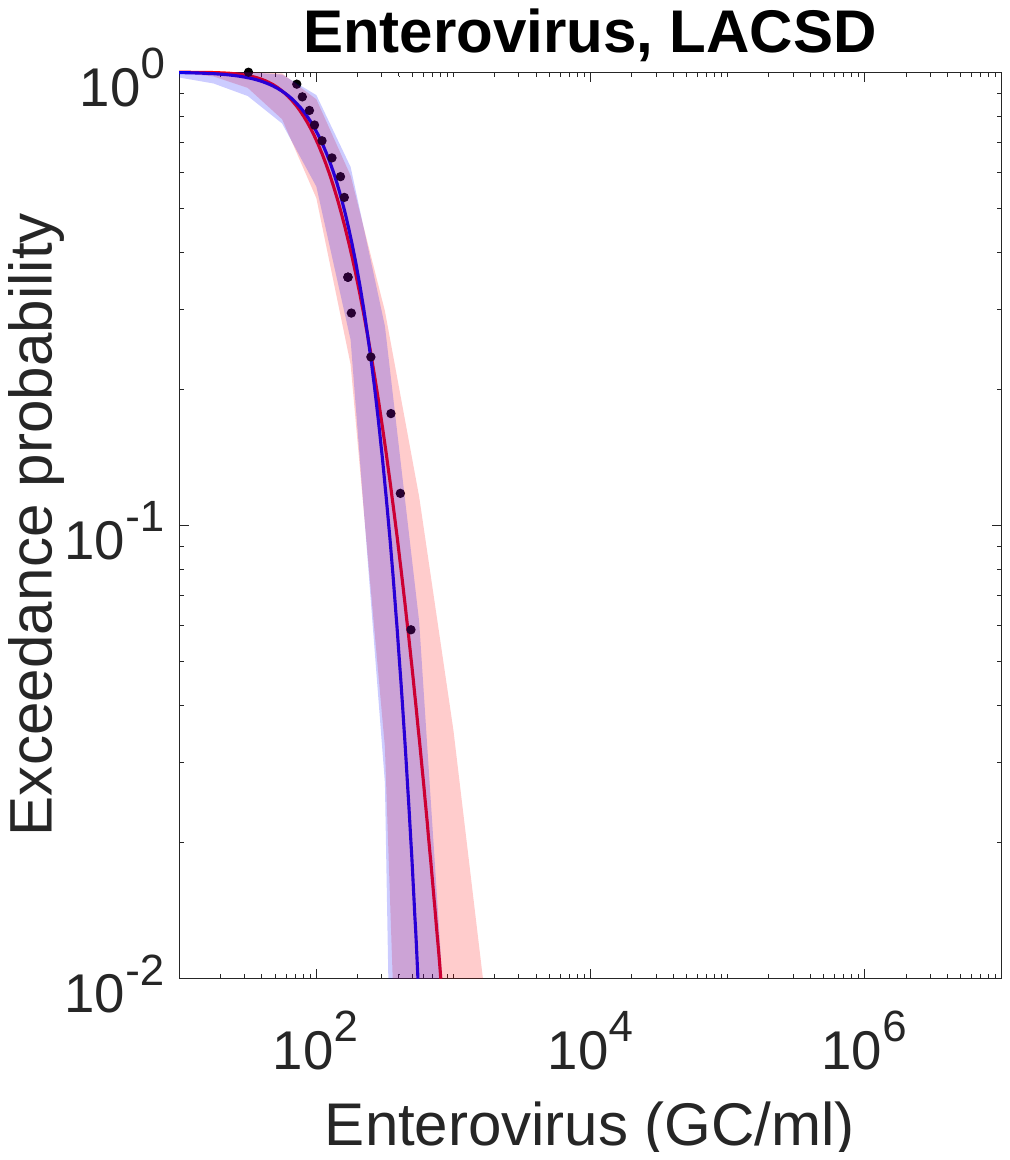

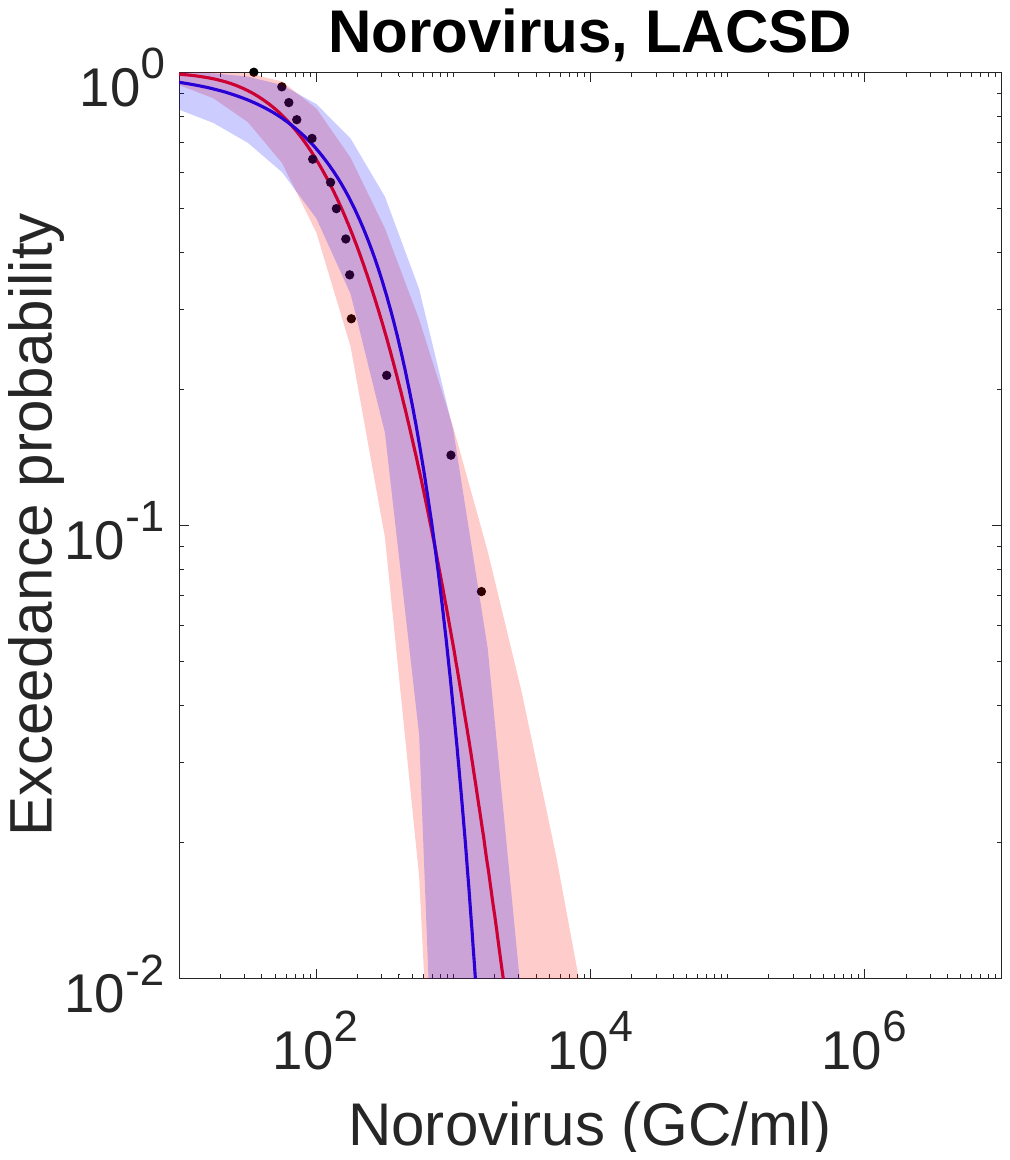

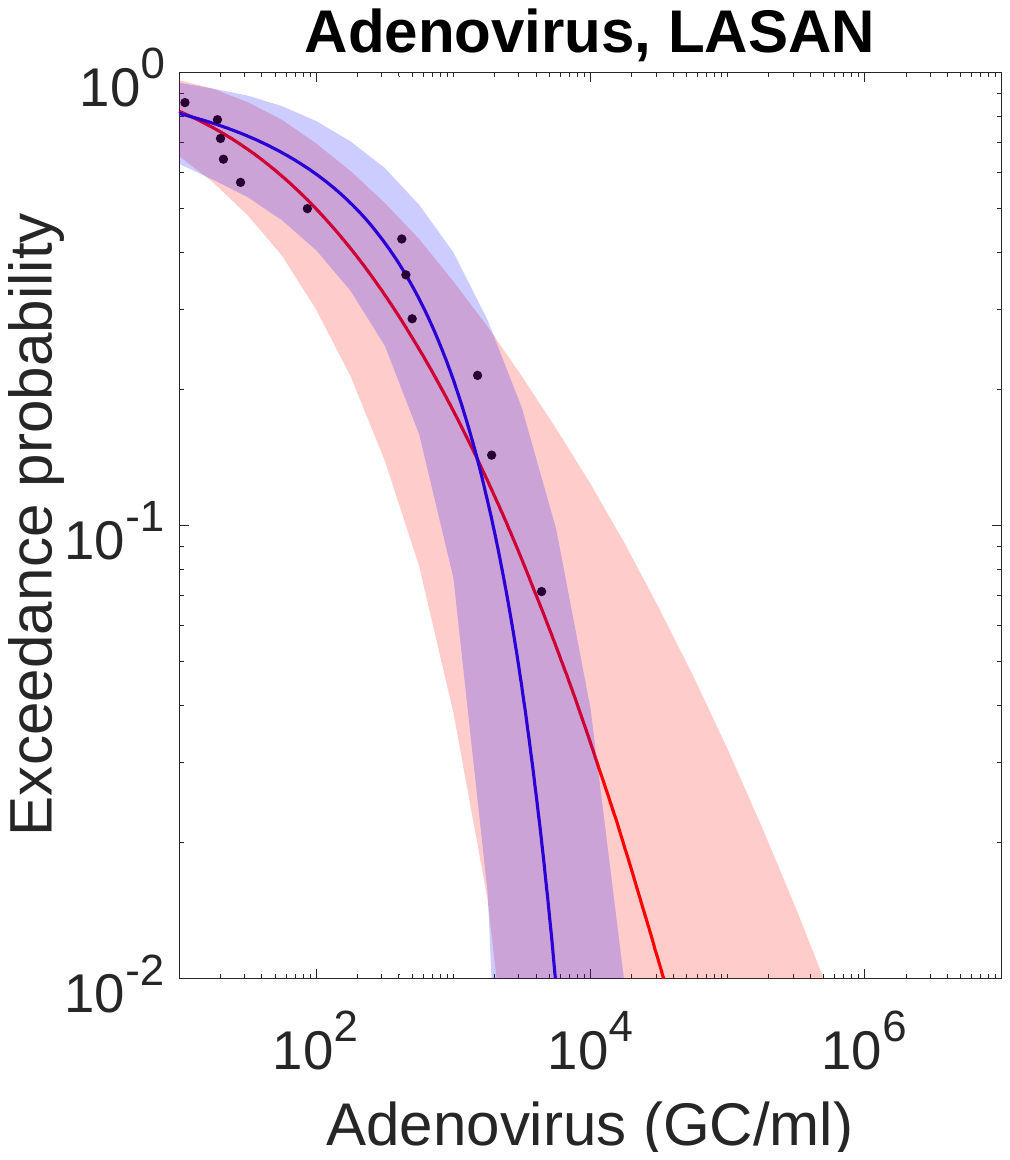

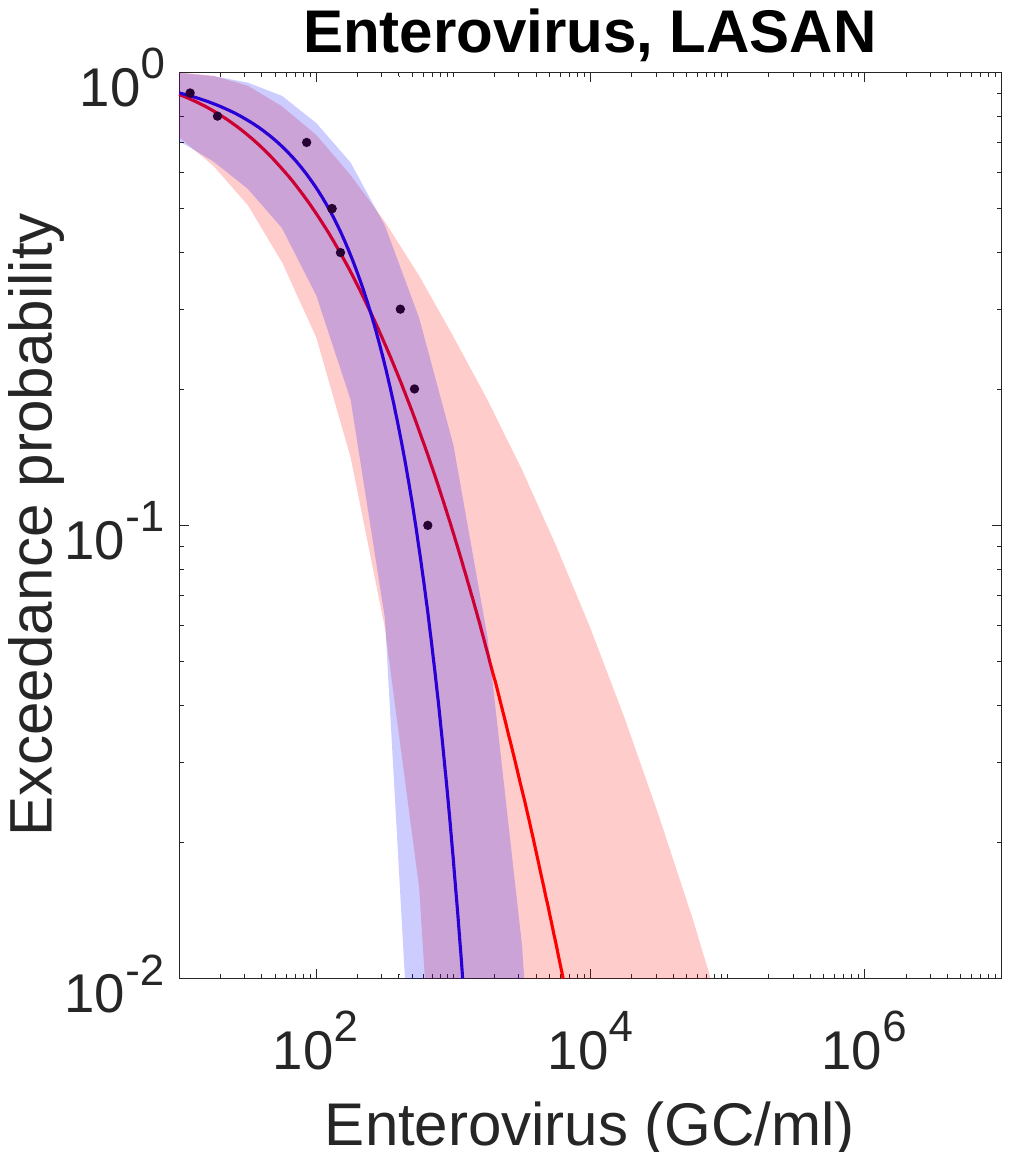

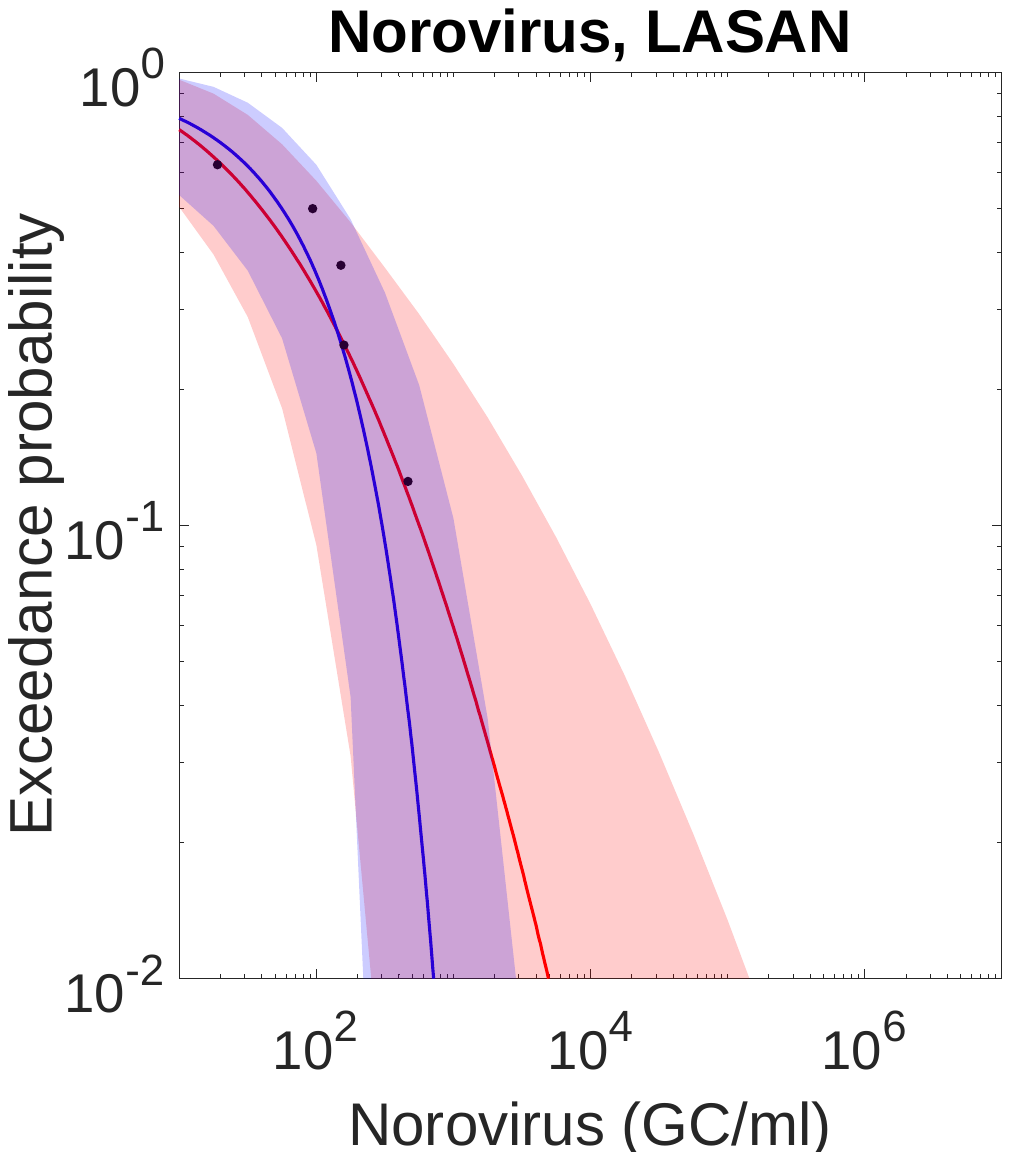

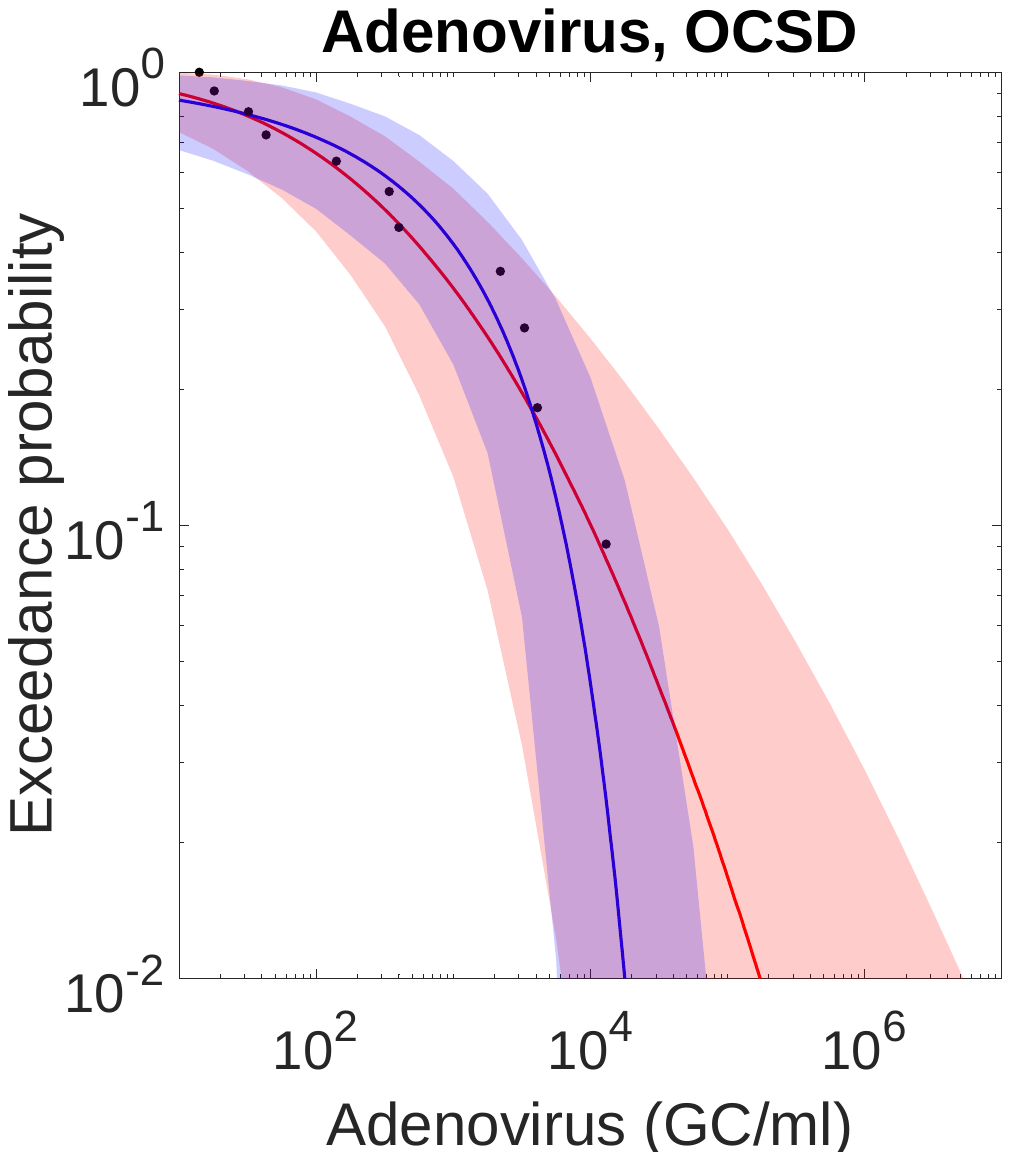

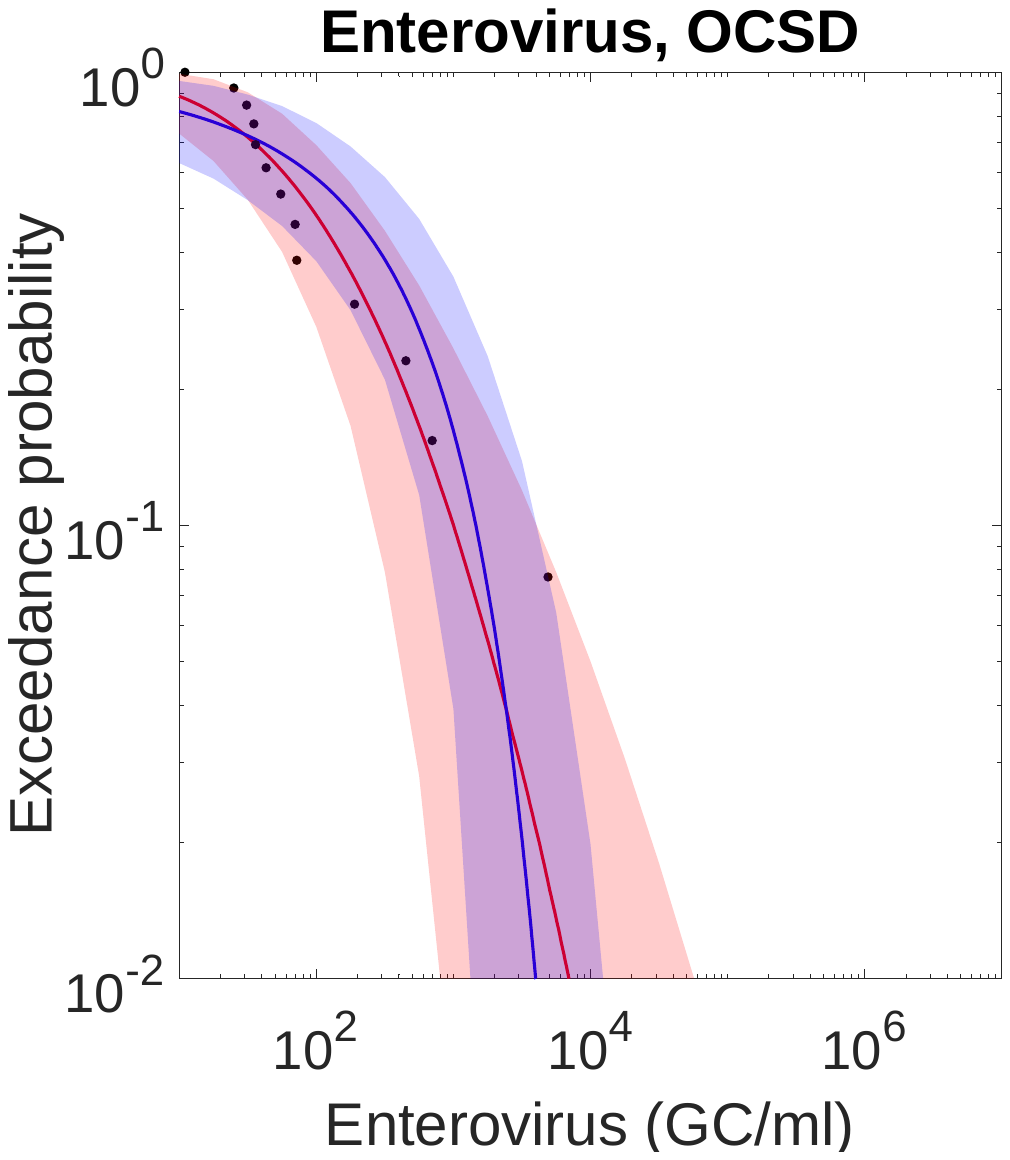

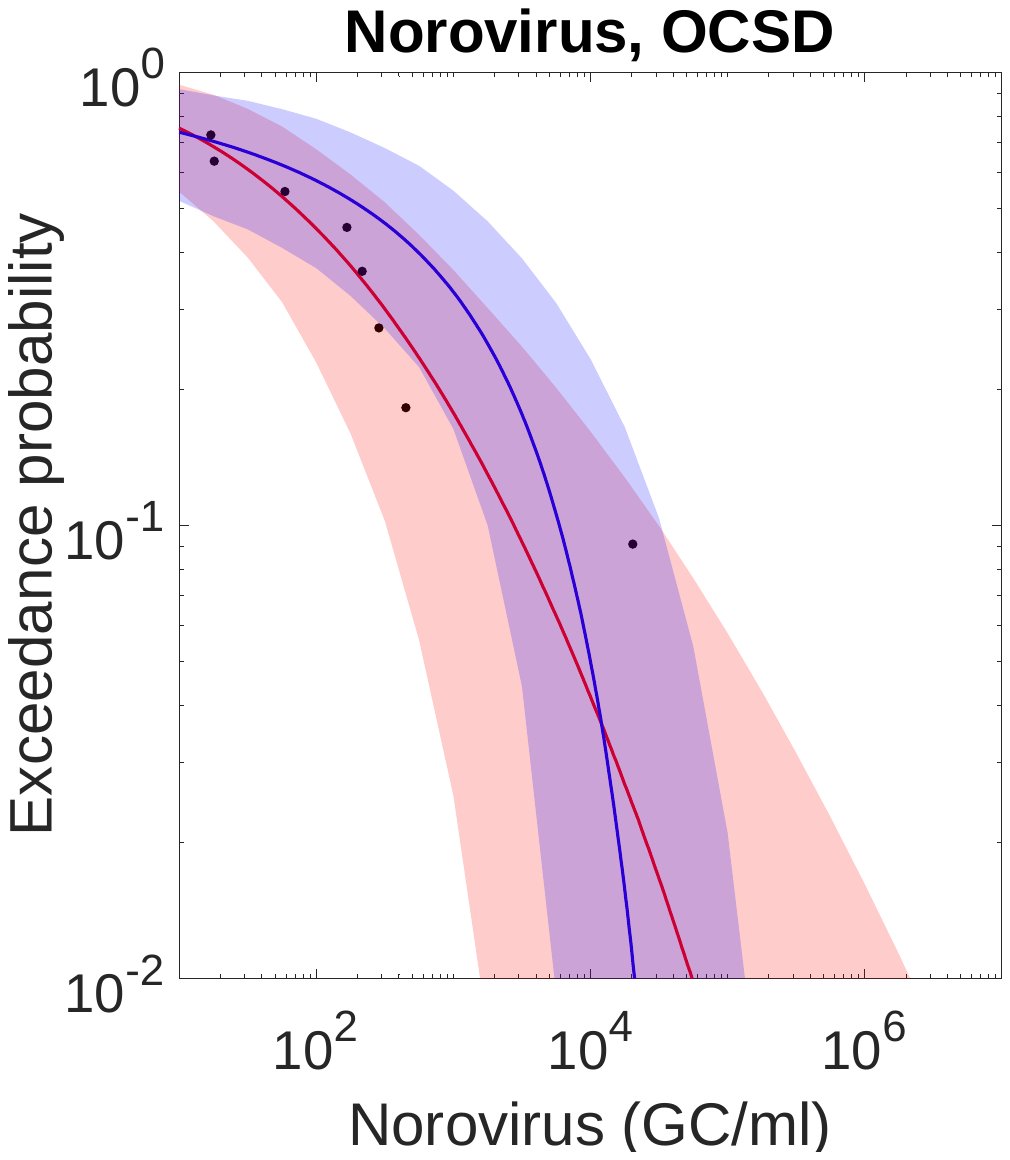

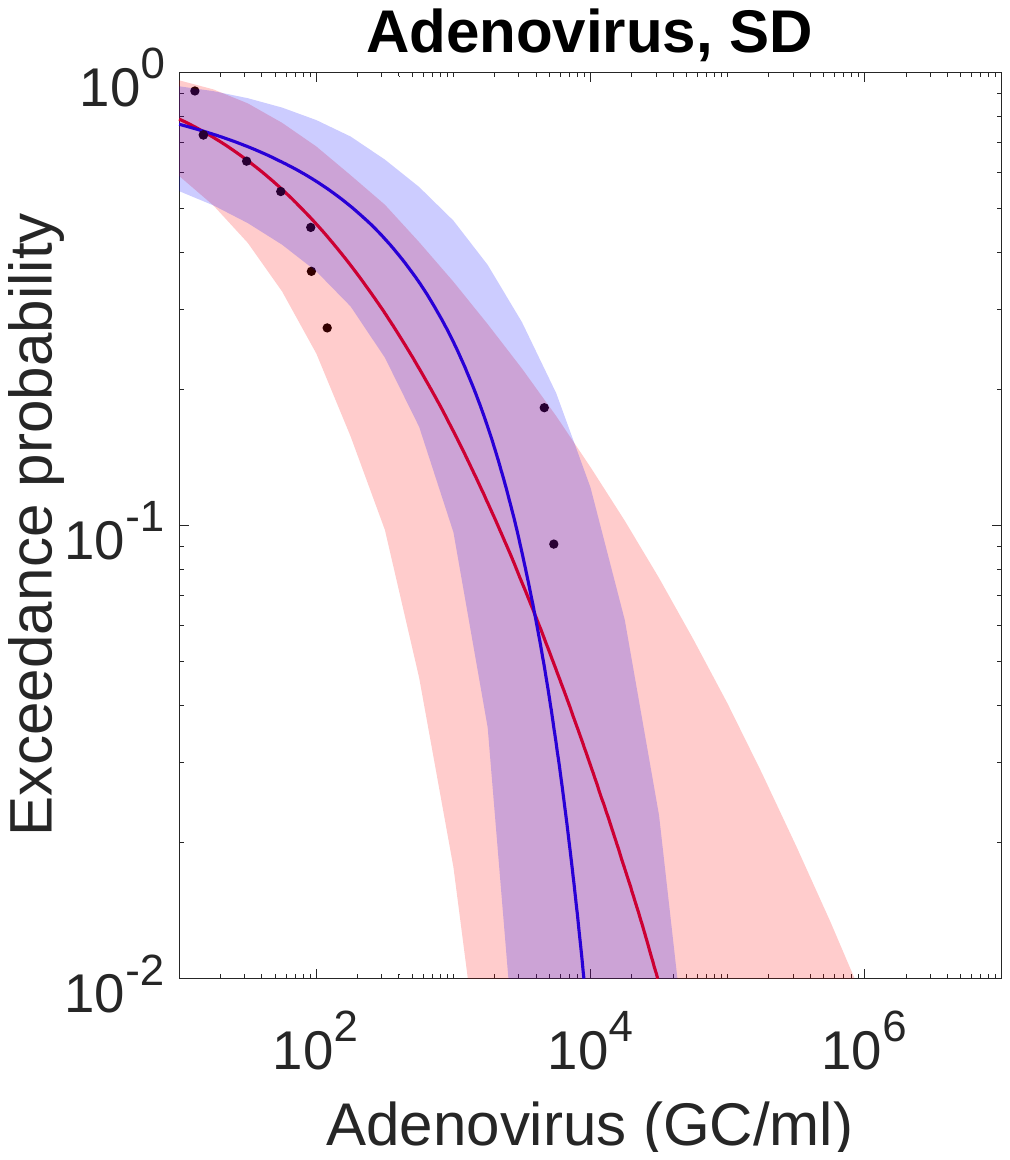

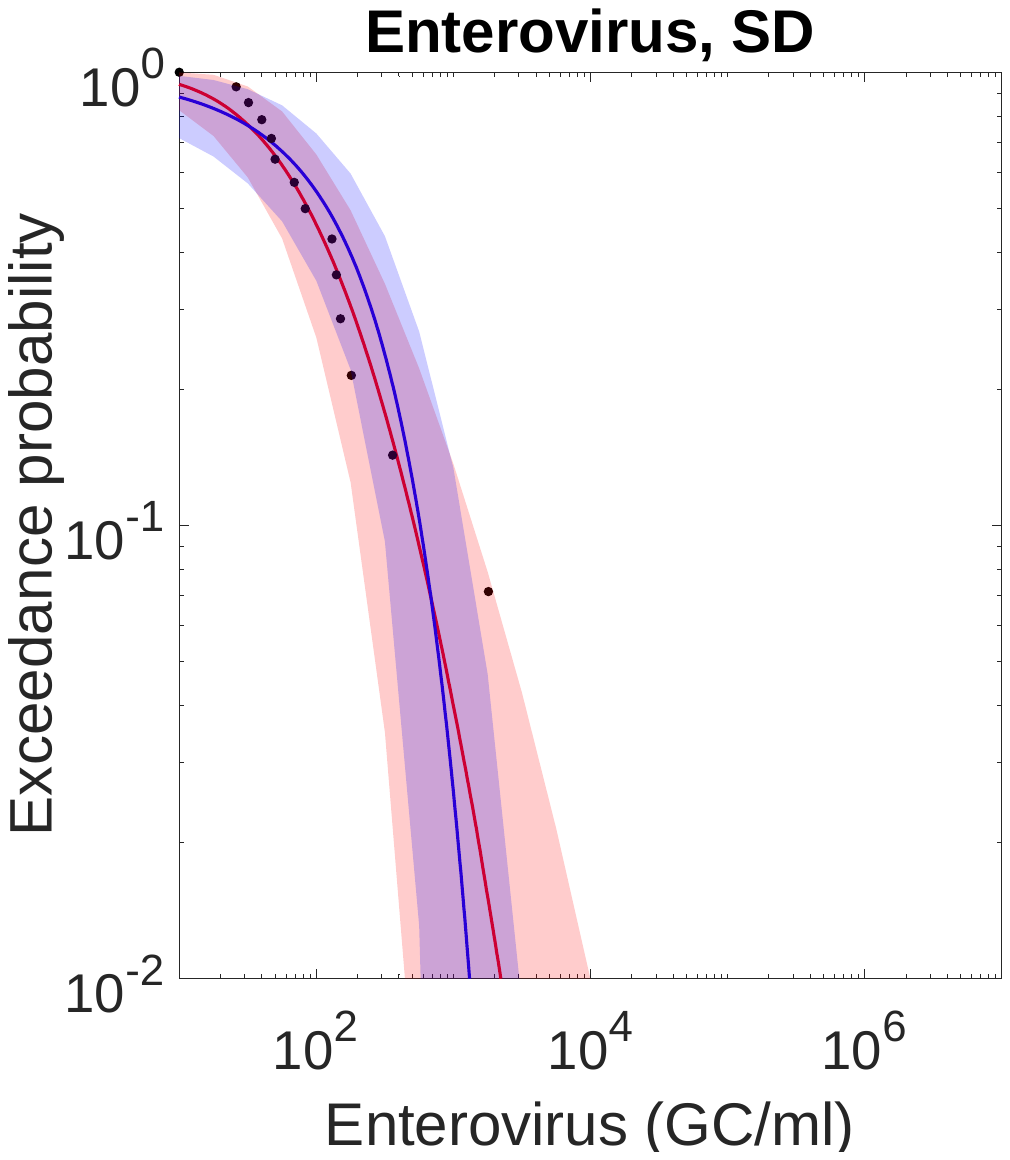

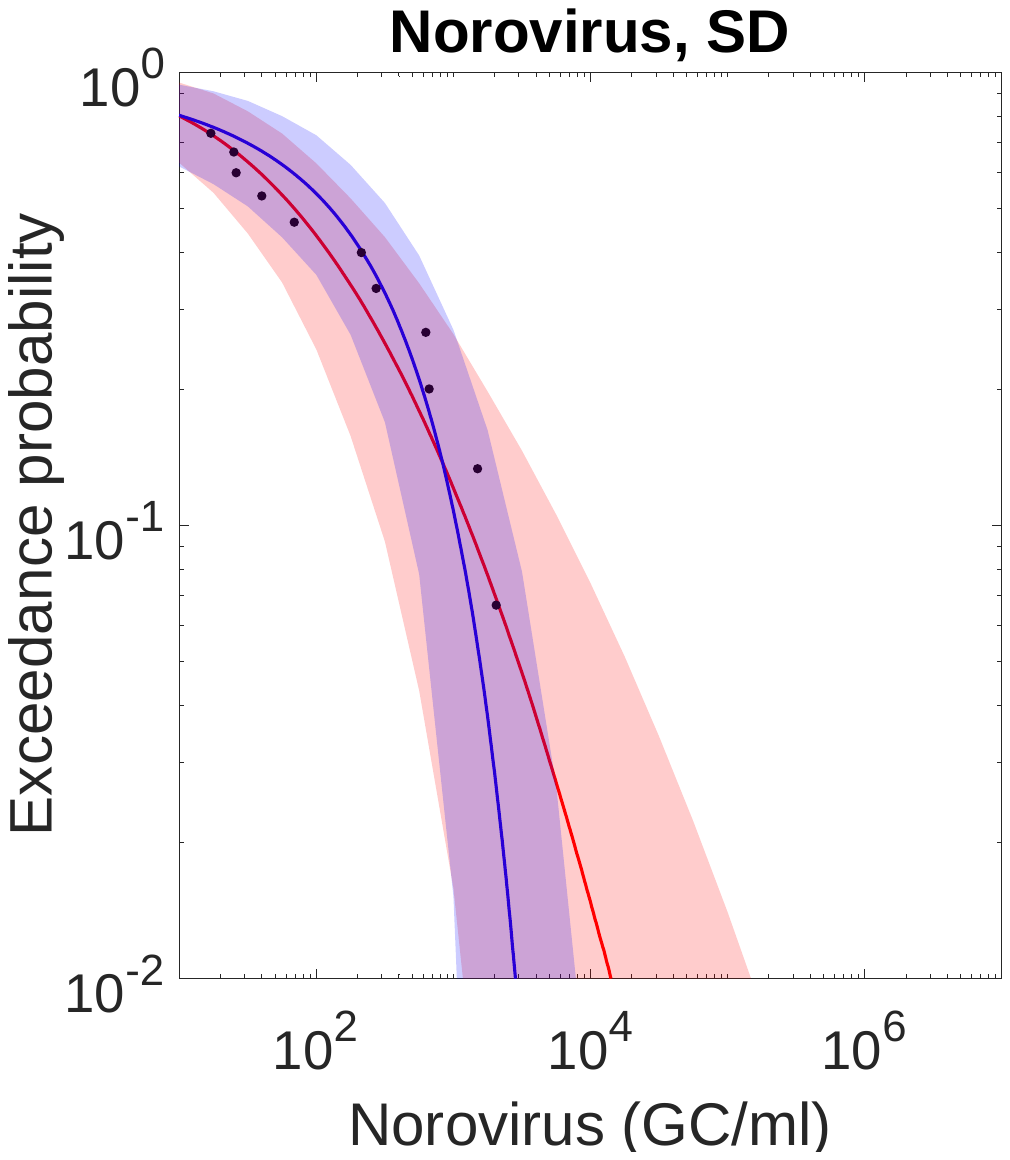

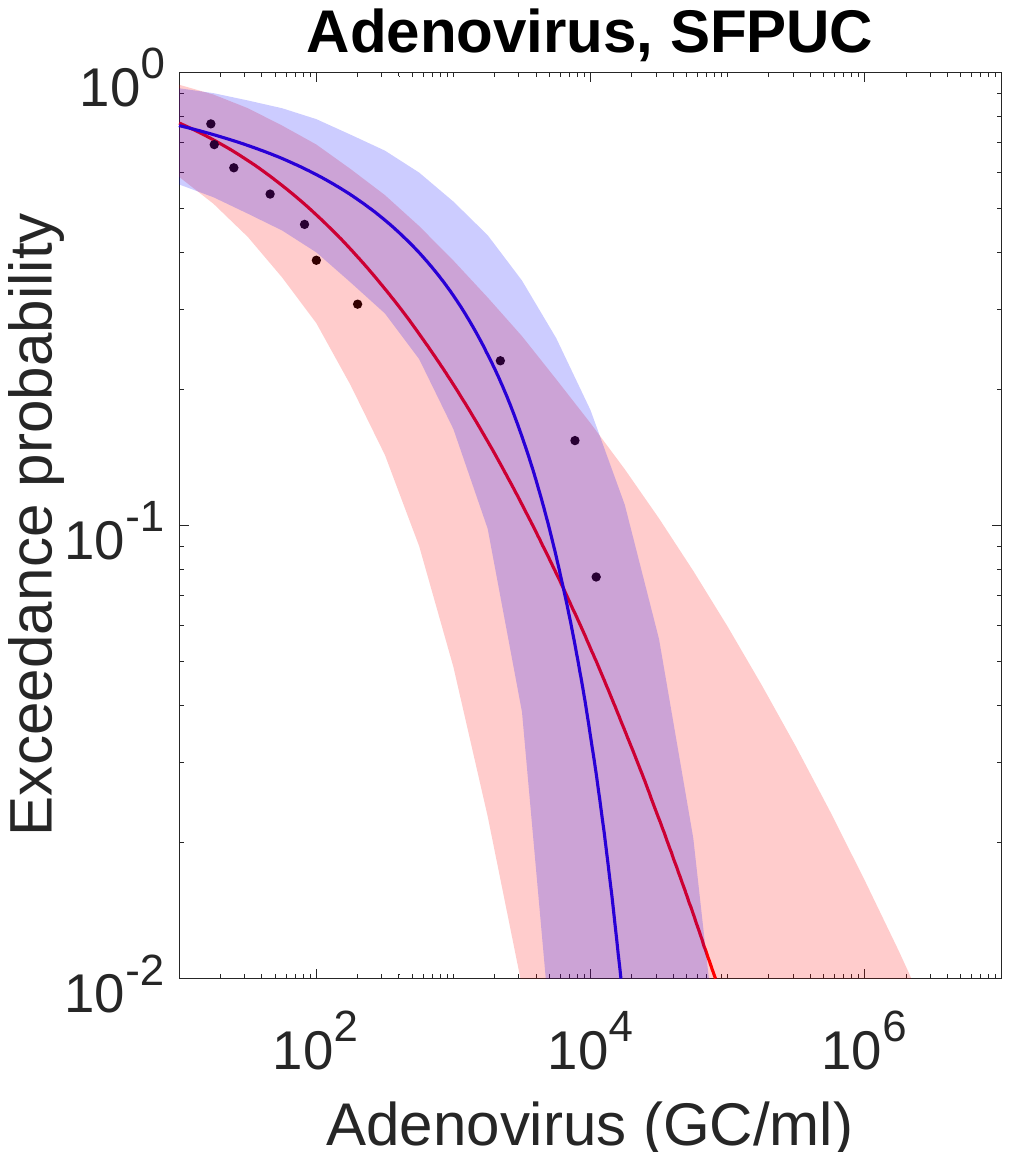

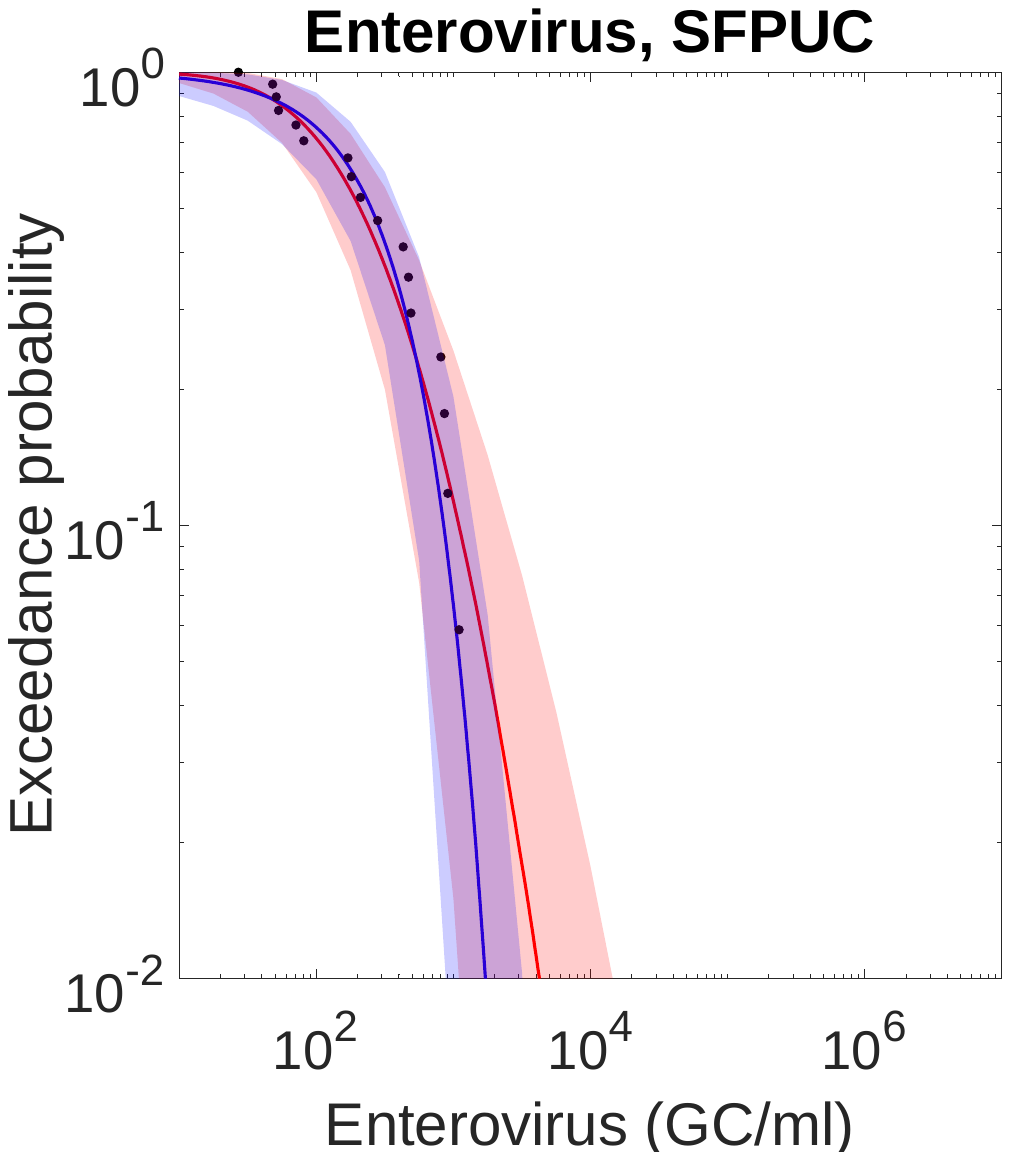

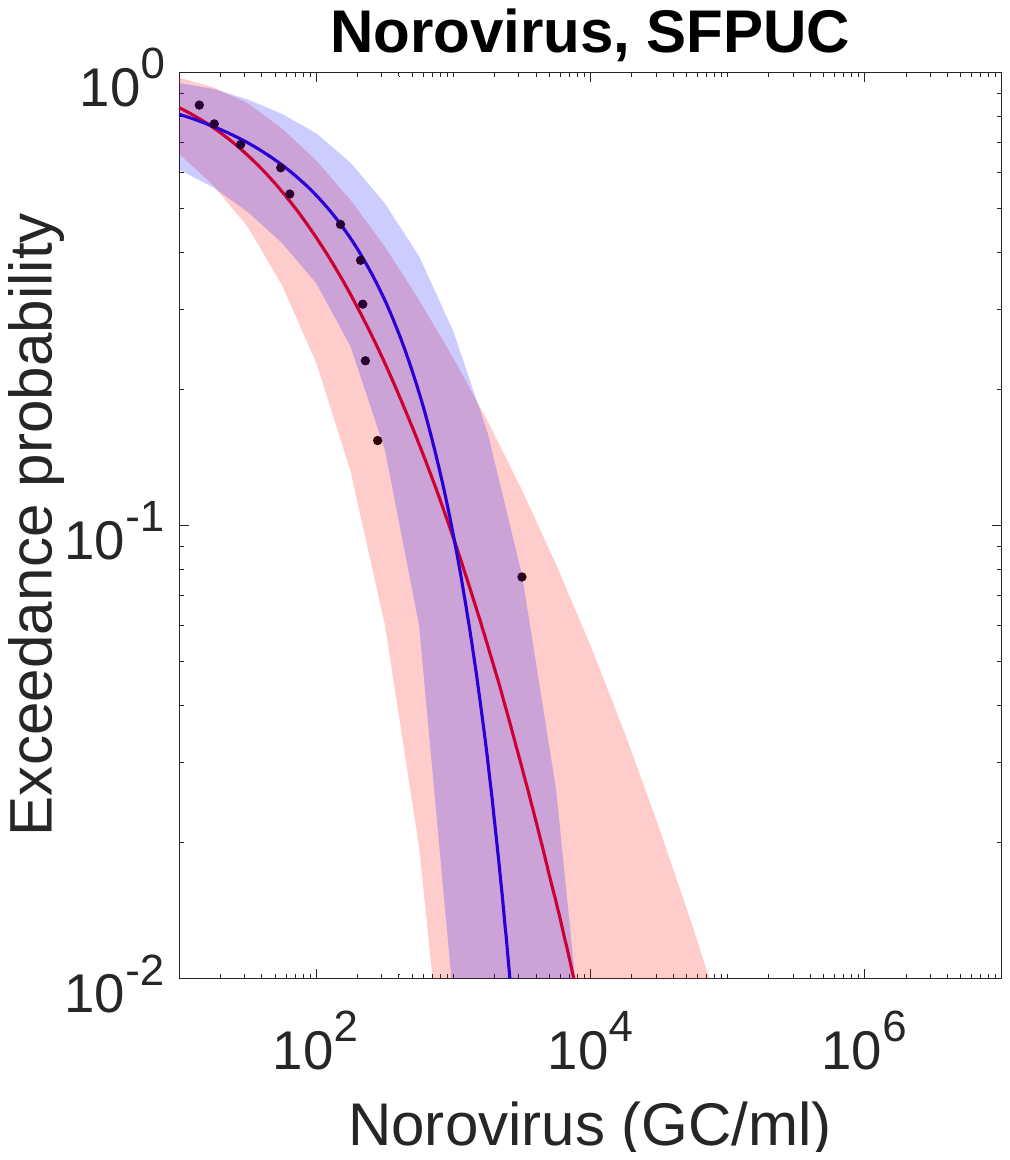

**Fig. S3.** Complementary cumulative distribution functions (CCDFs) of the mixed Poisson distributions of enteric virus concentrations for the eight wastewater treatment plants. Red curves represent PLN distributions and blue curves represent PGA distributions.

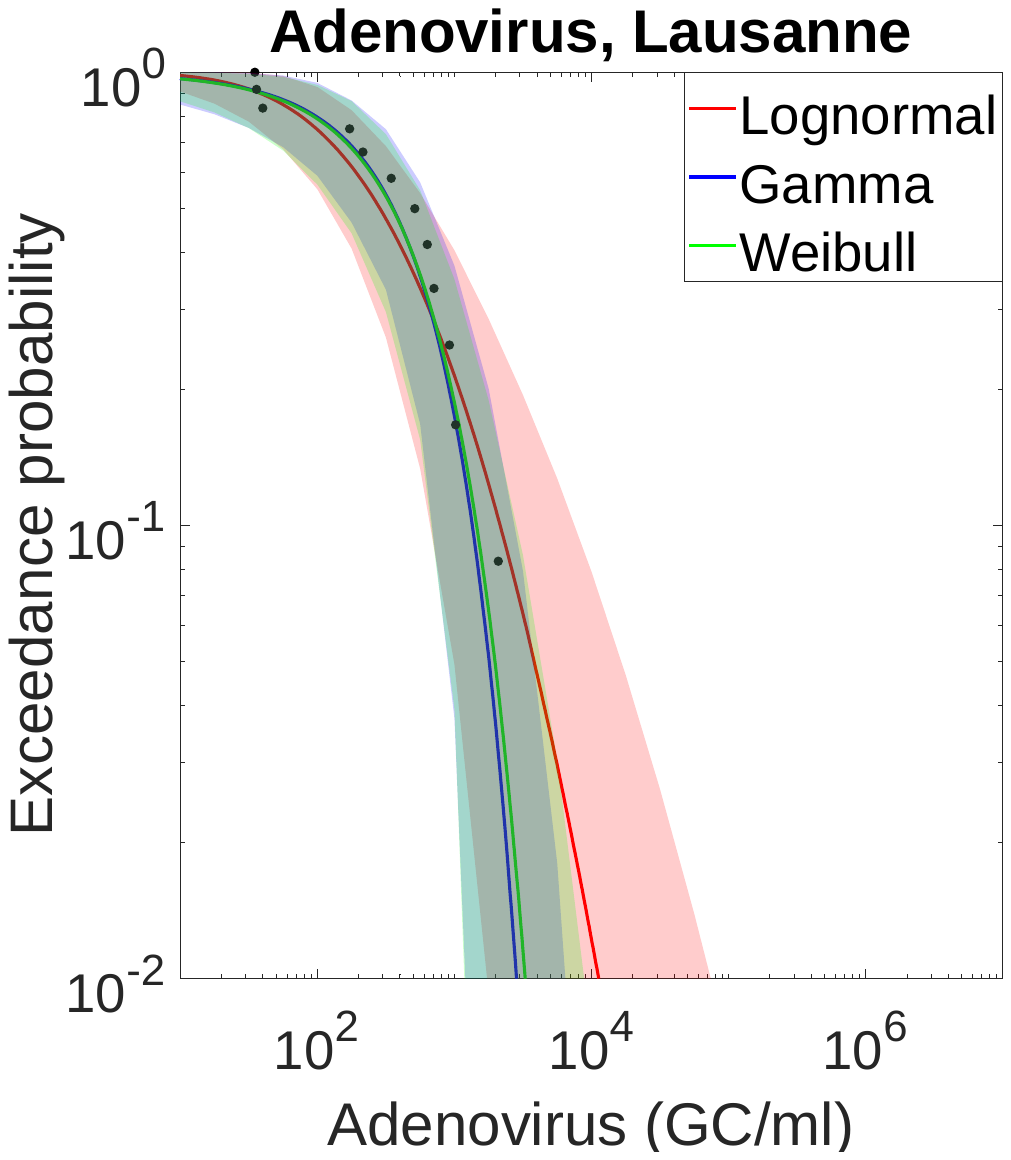

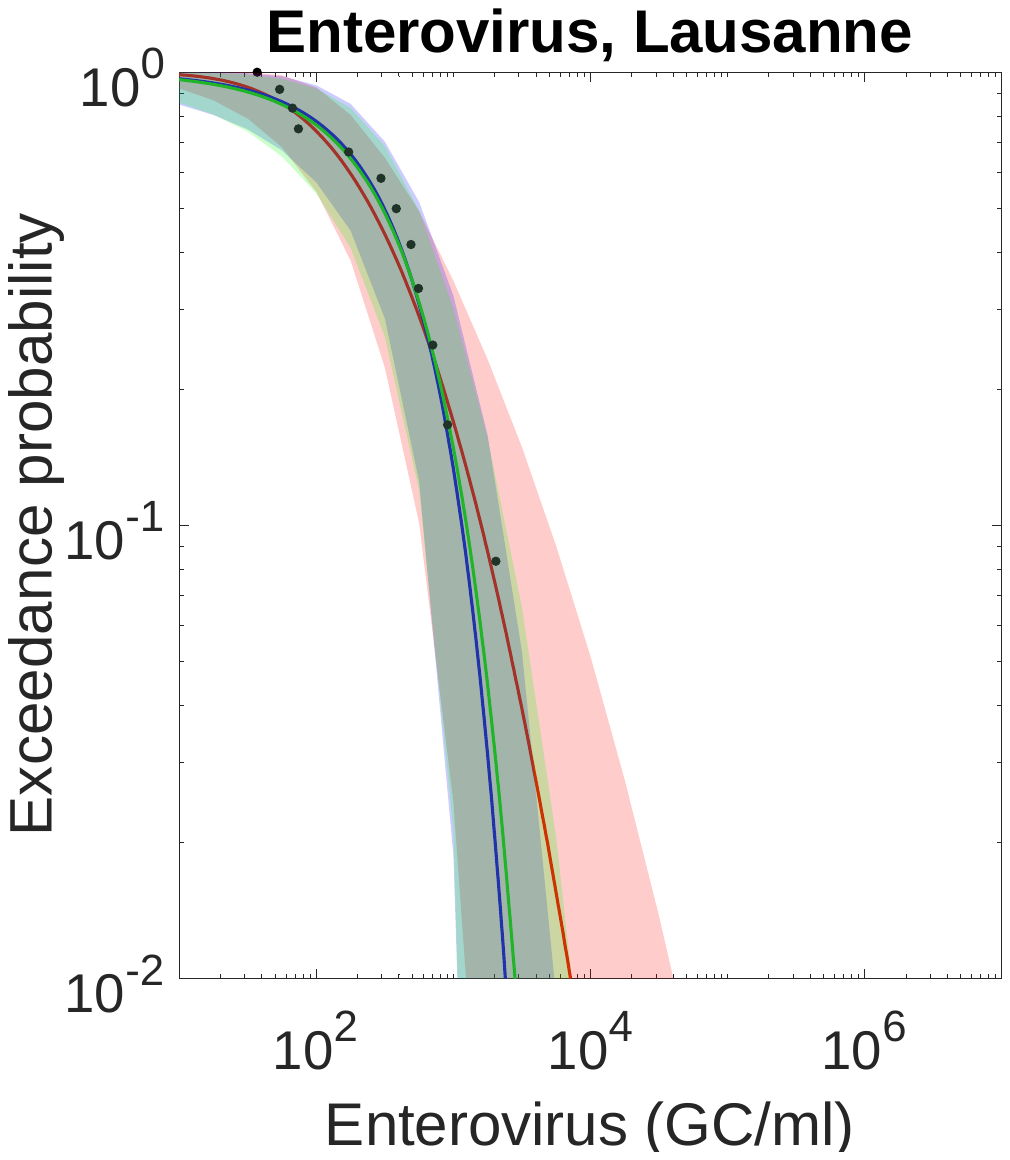

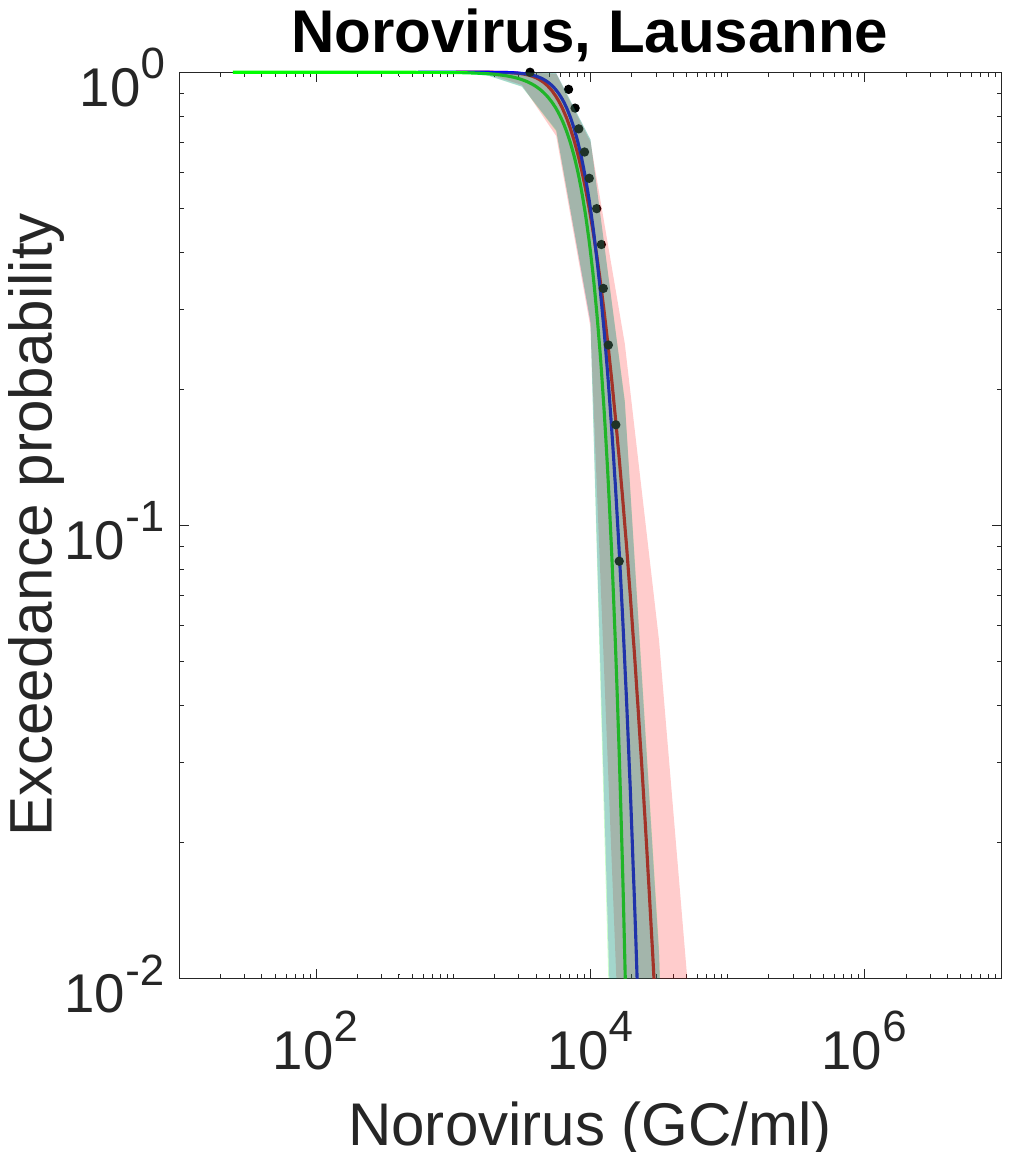

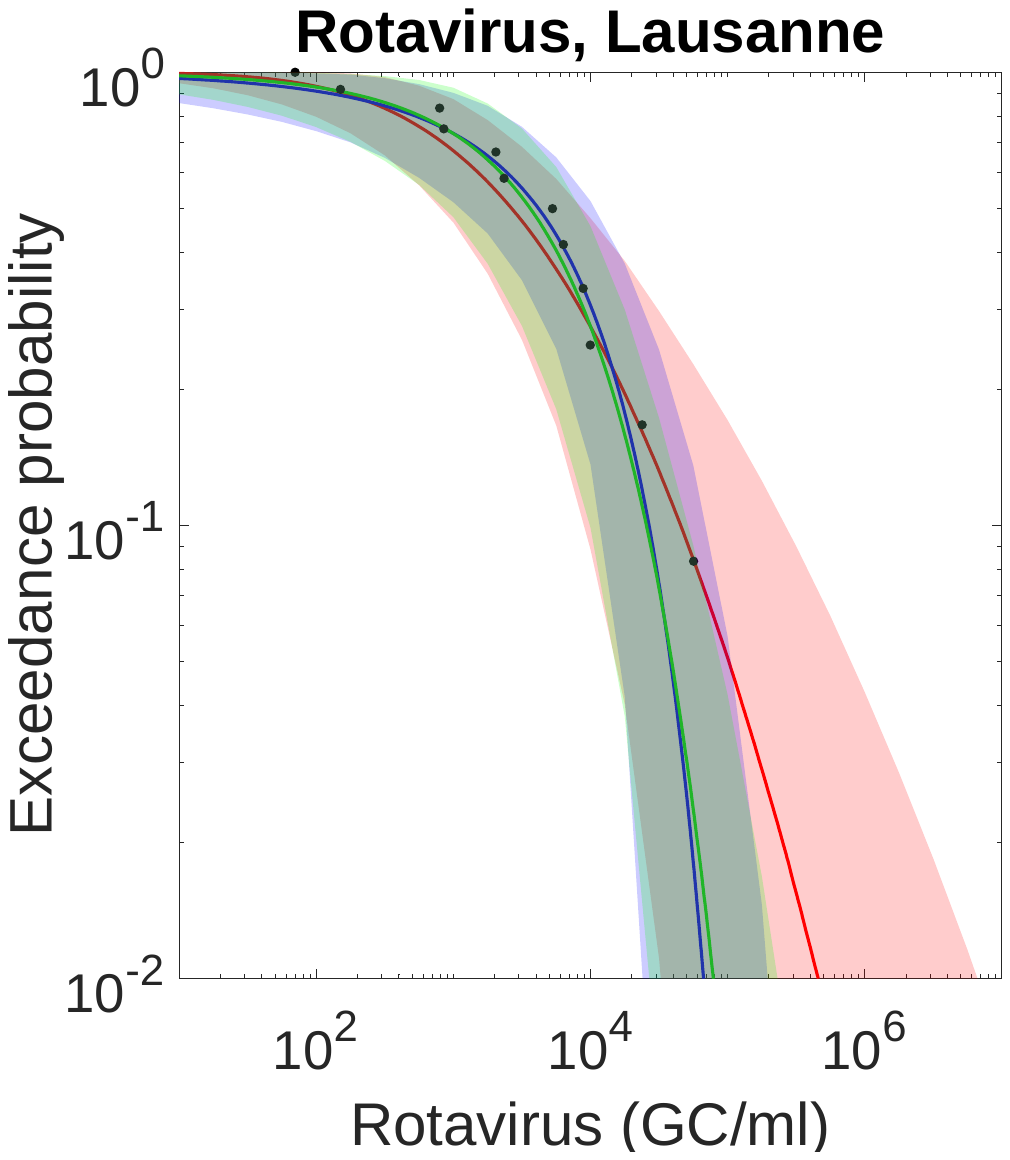

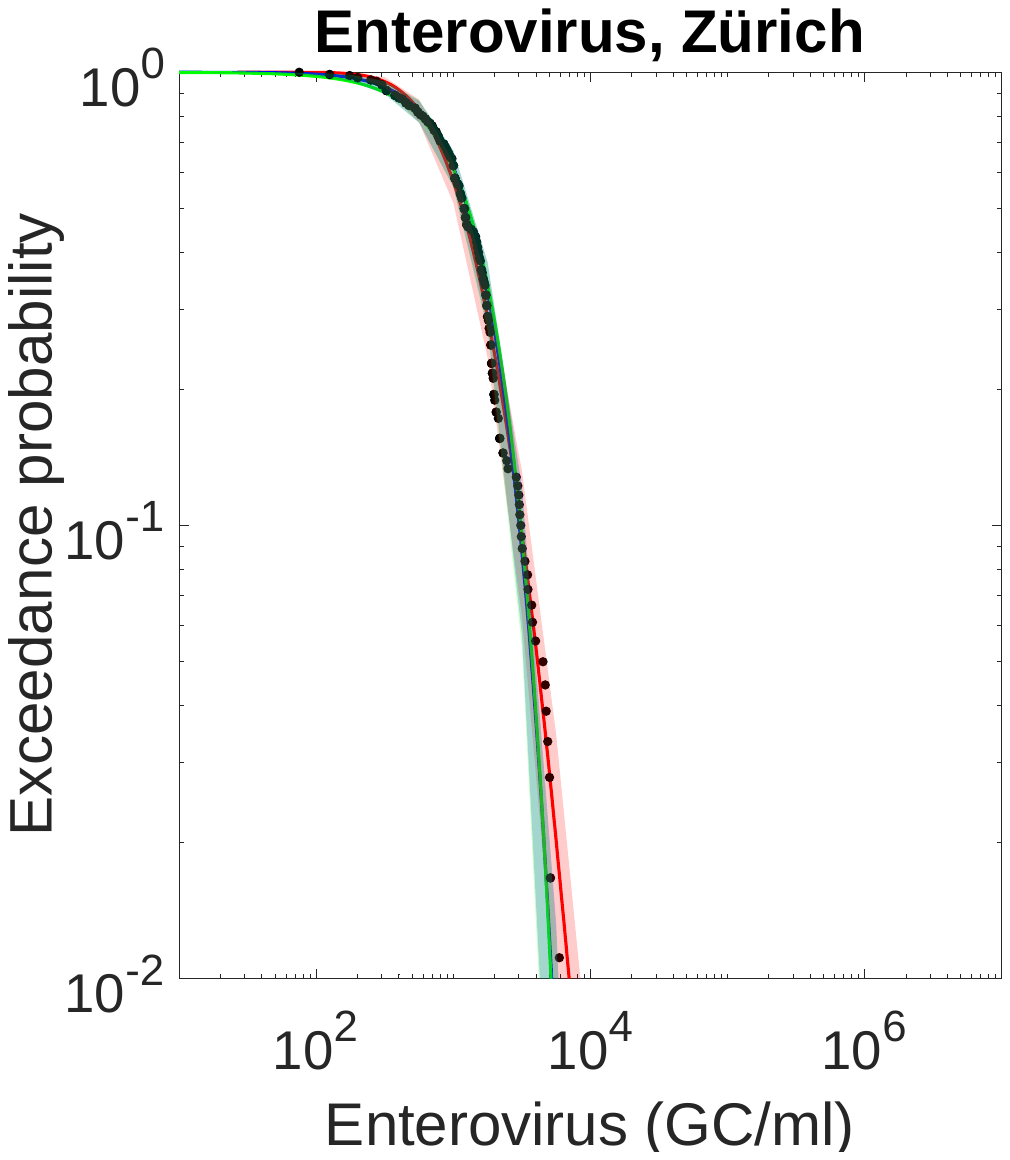

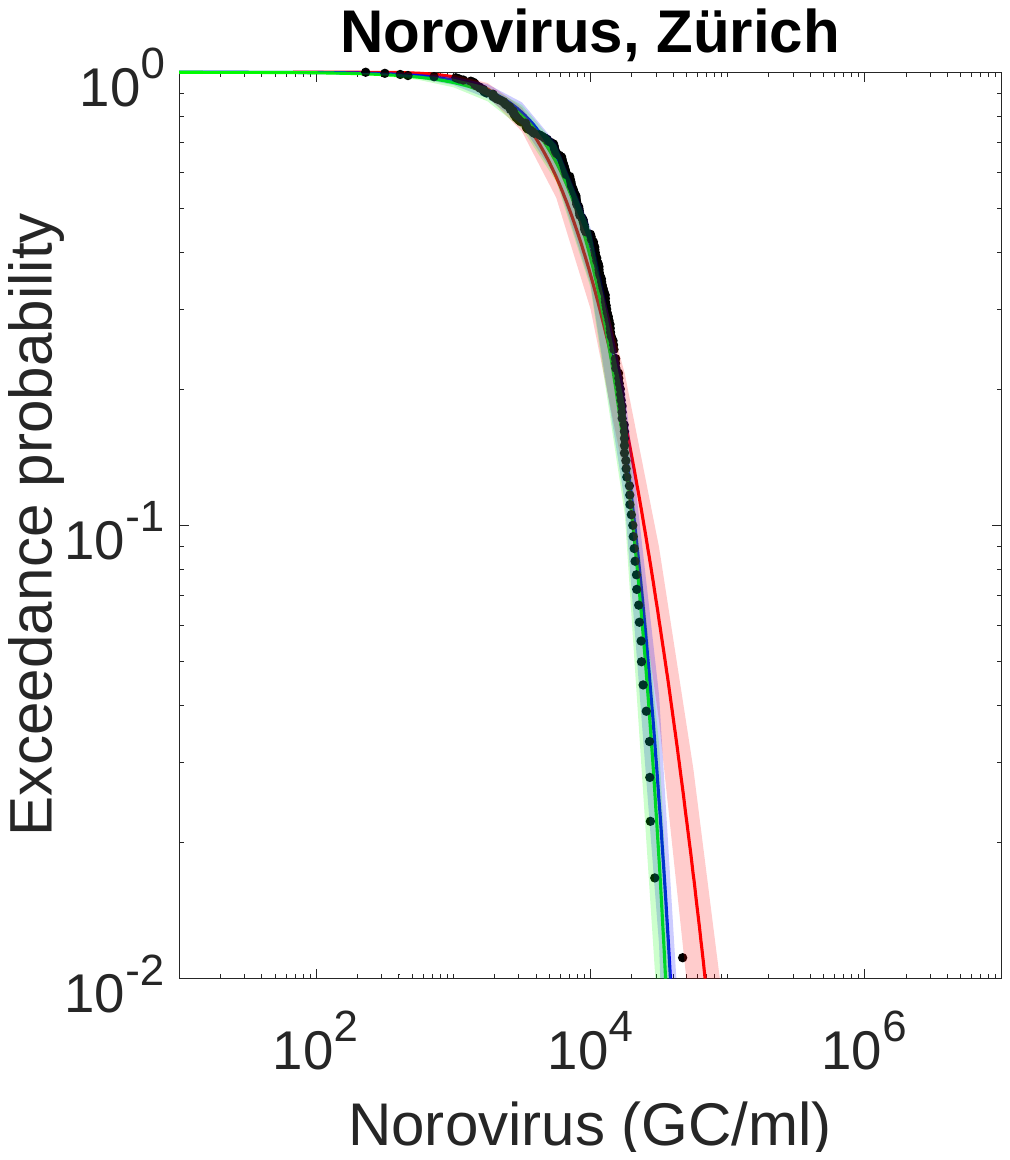

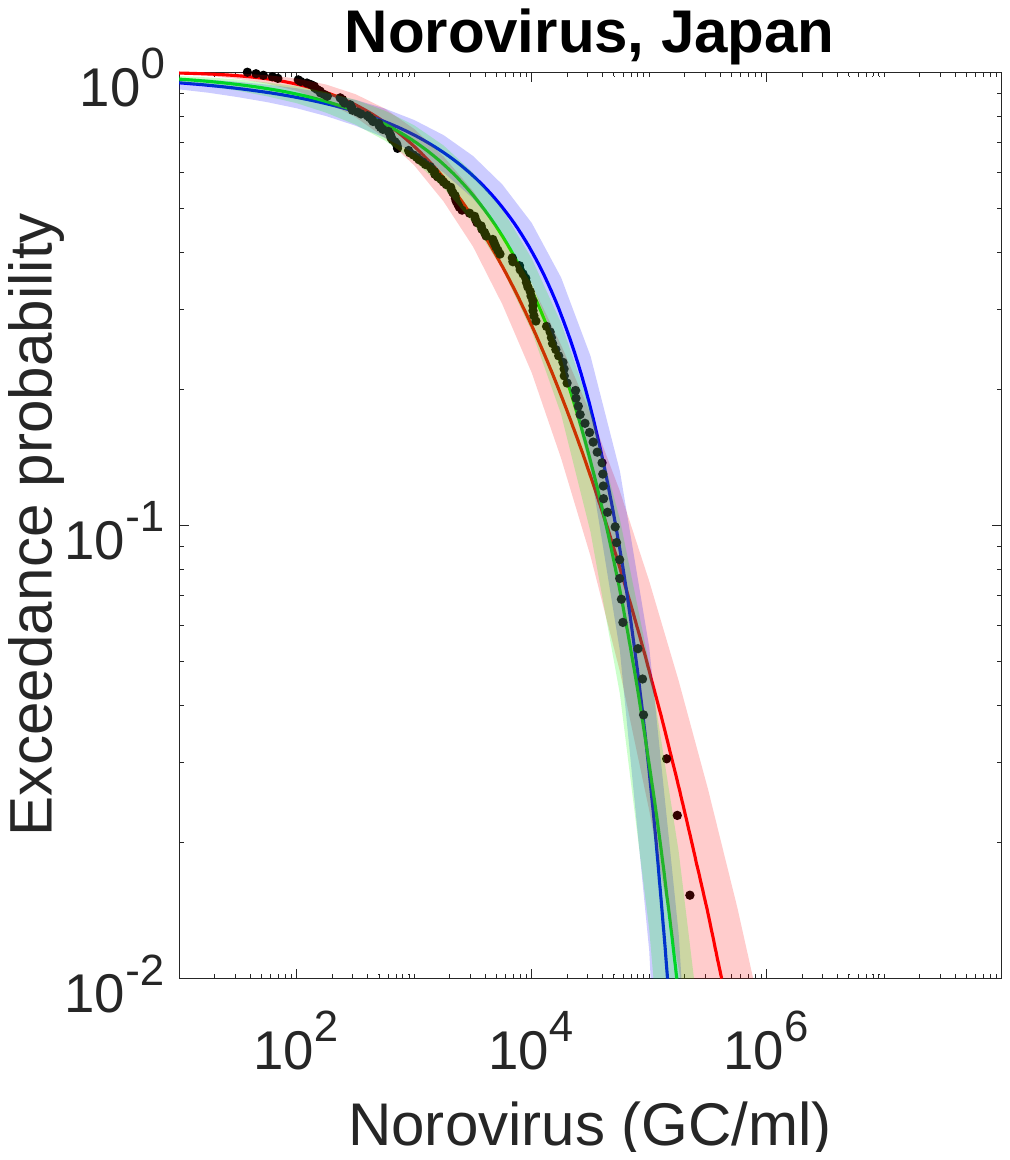

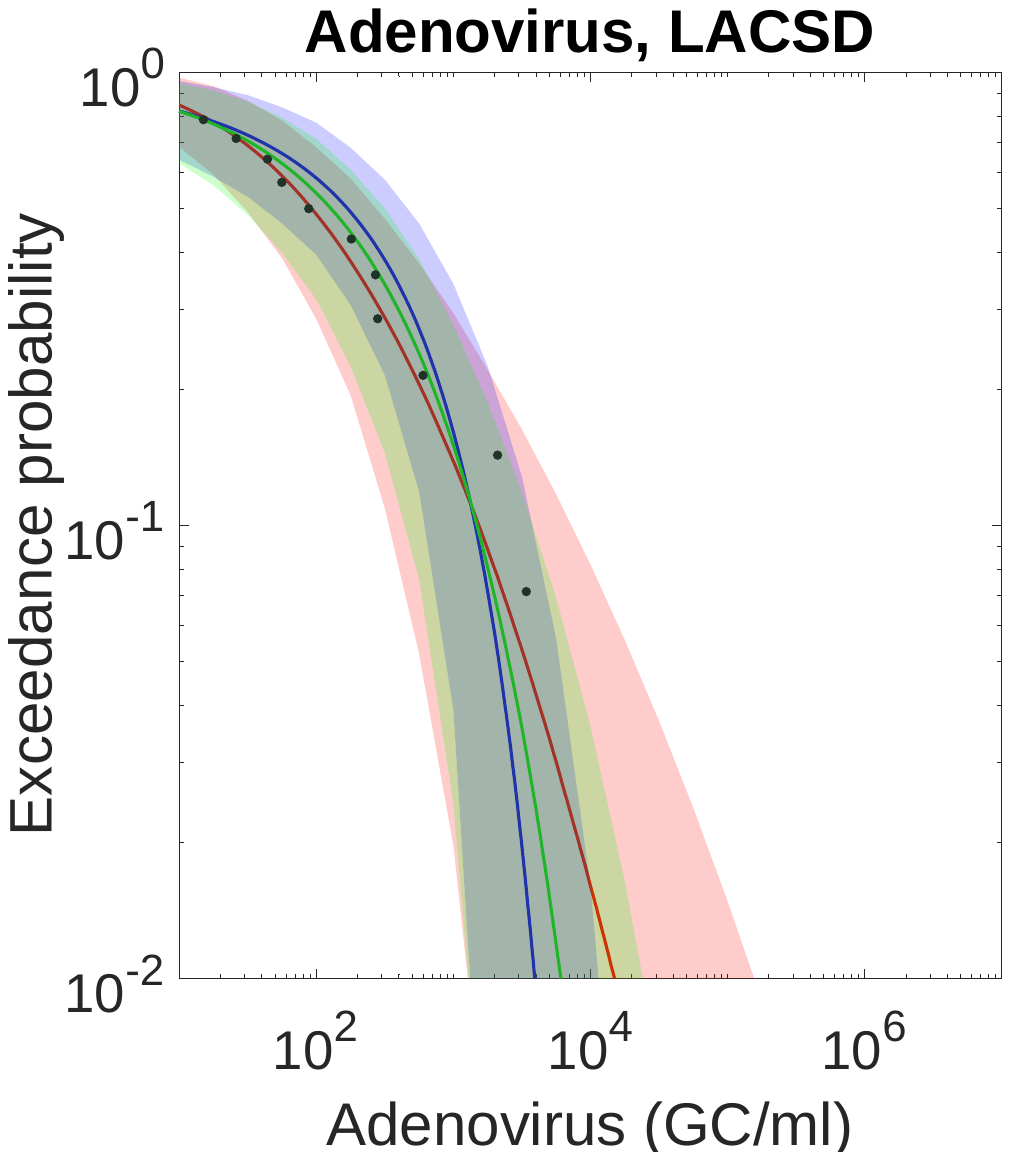

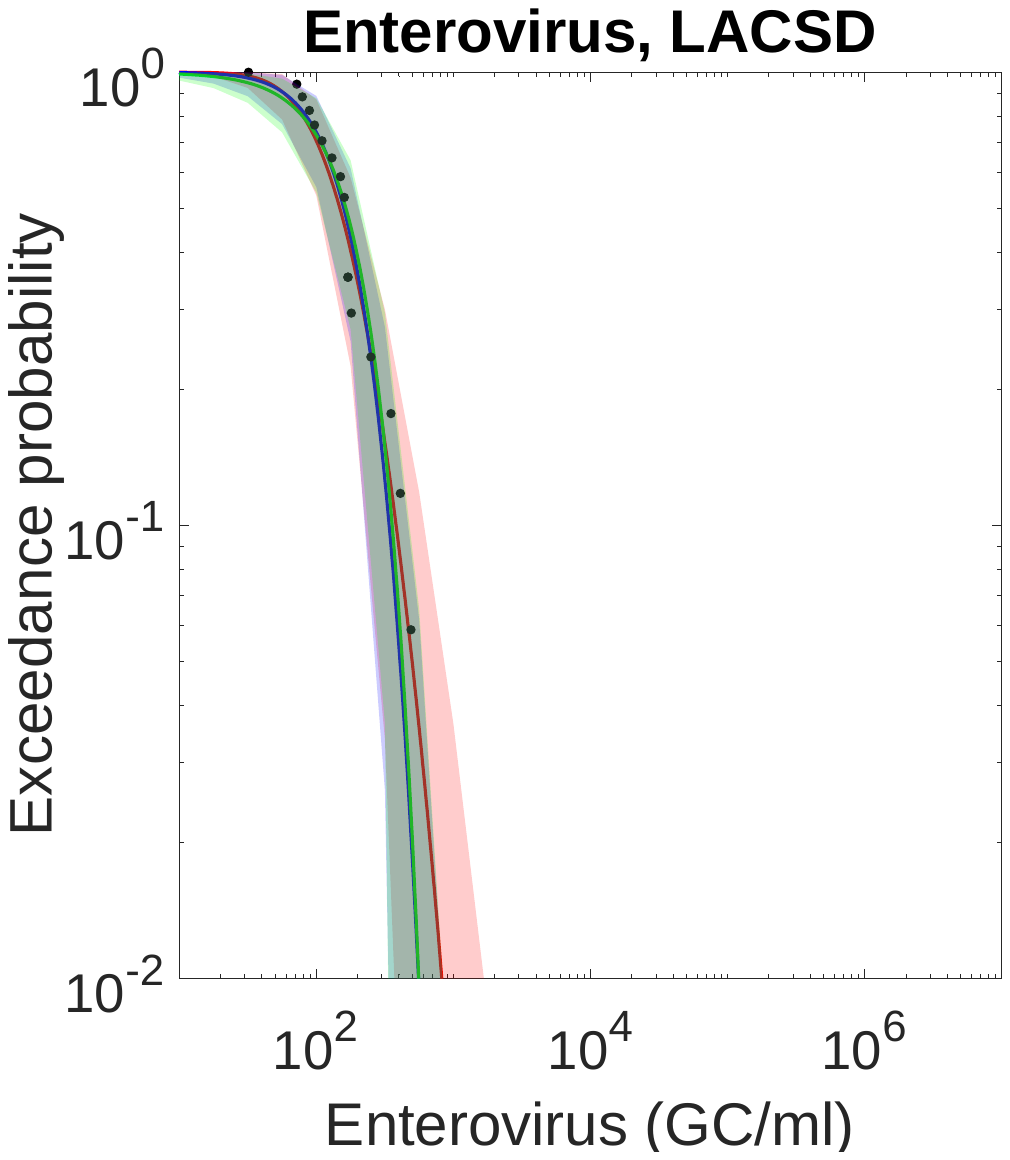

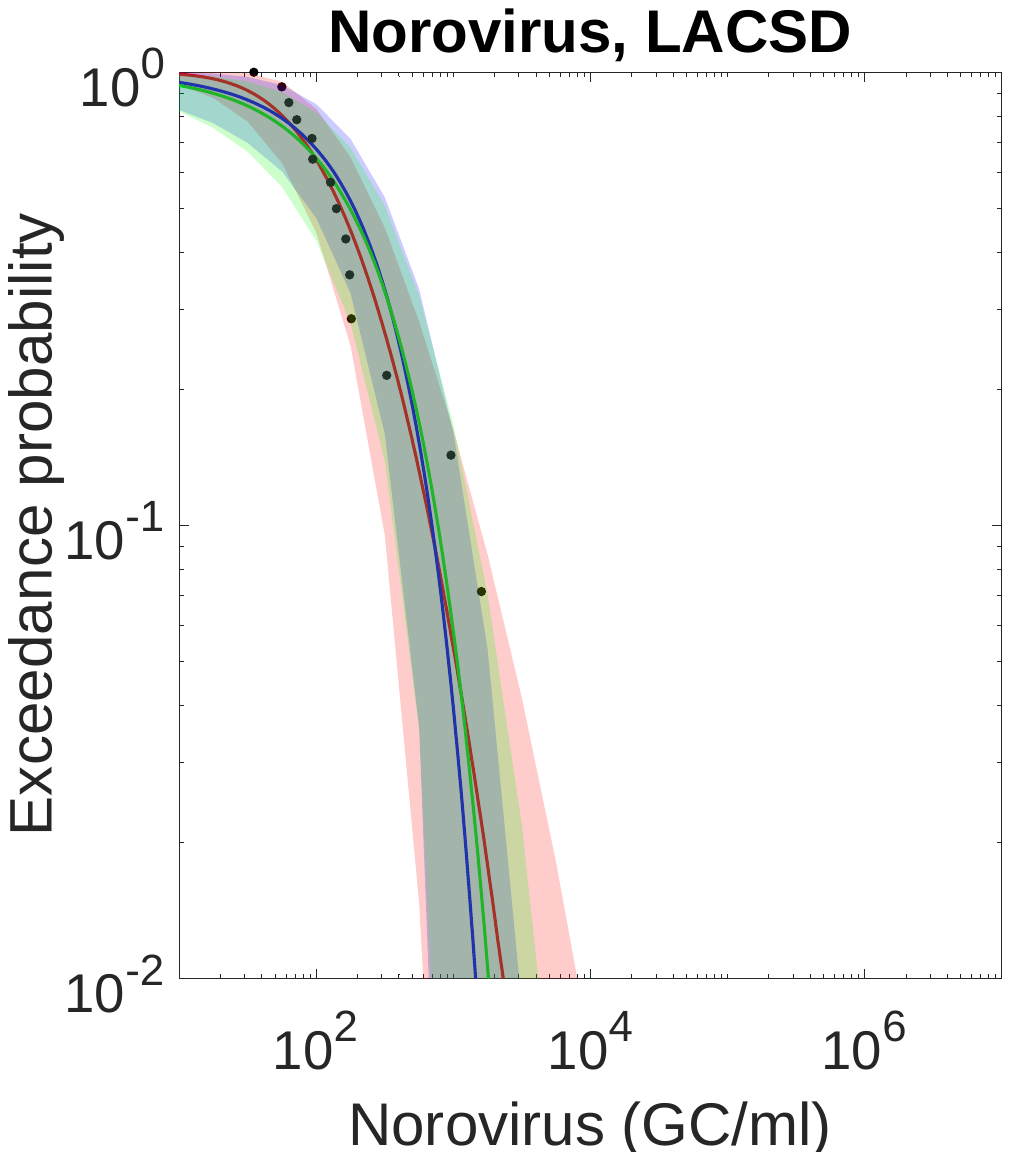

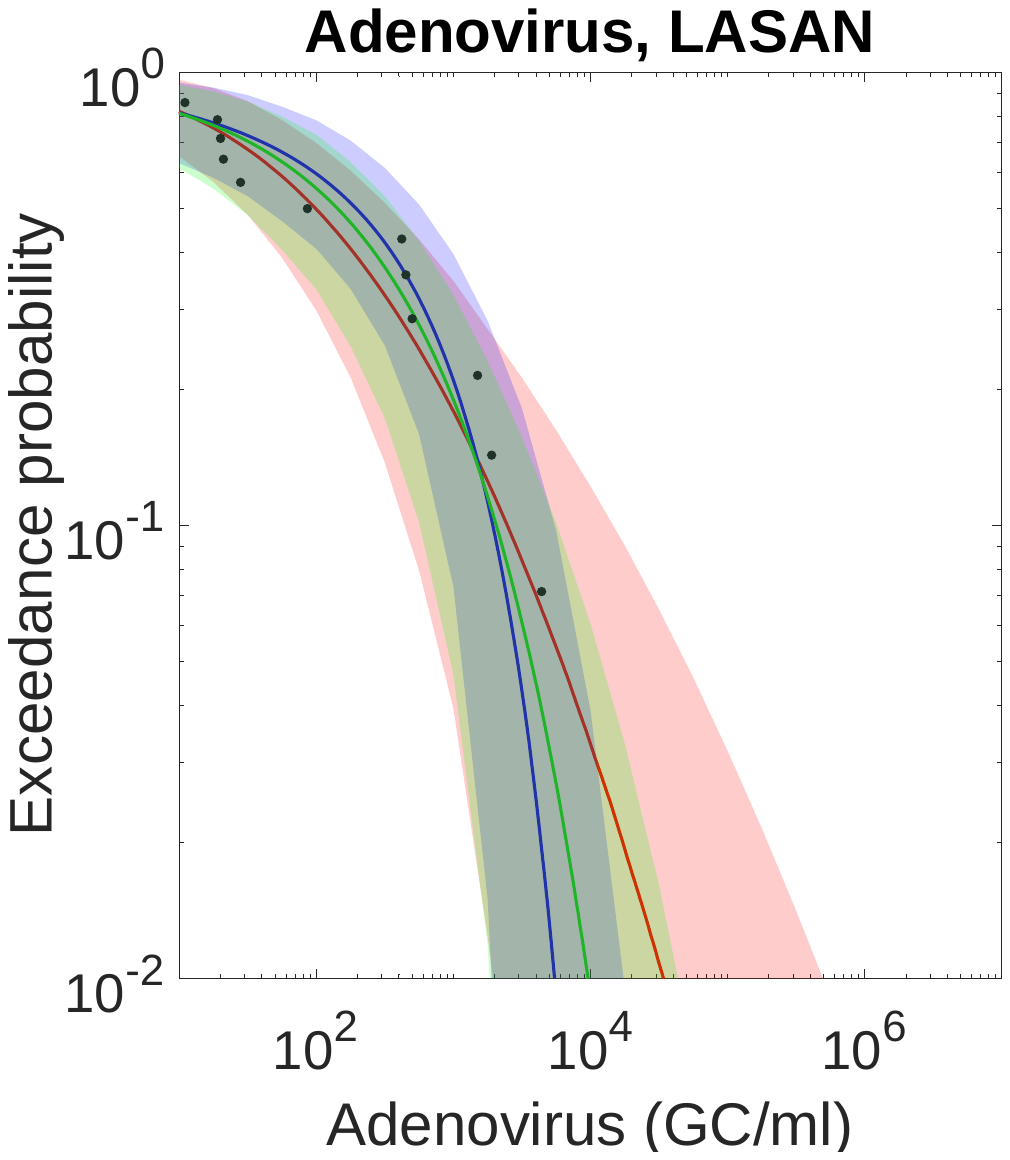

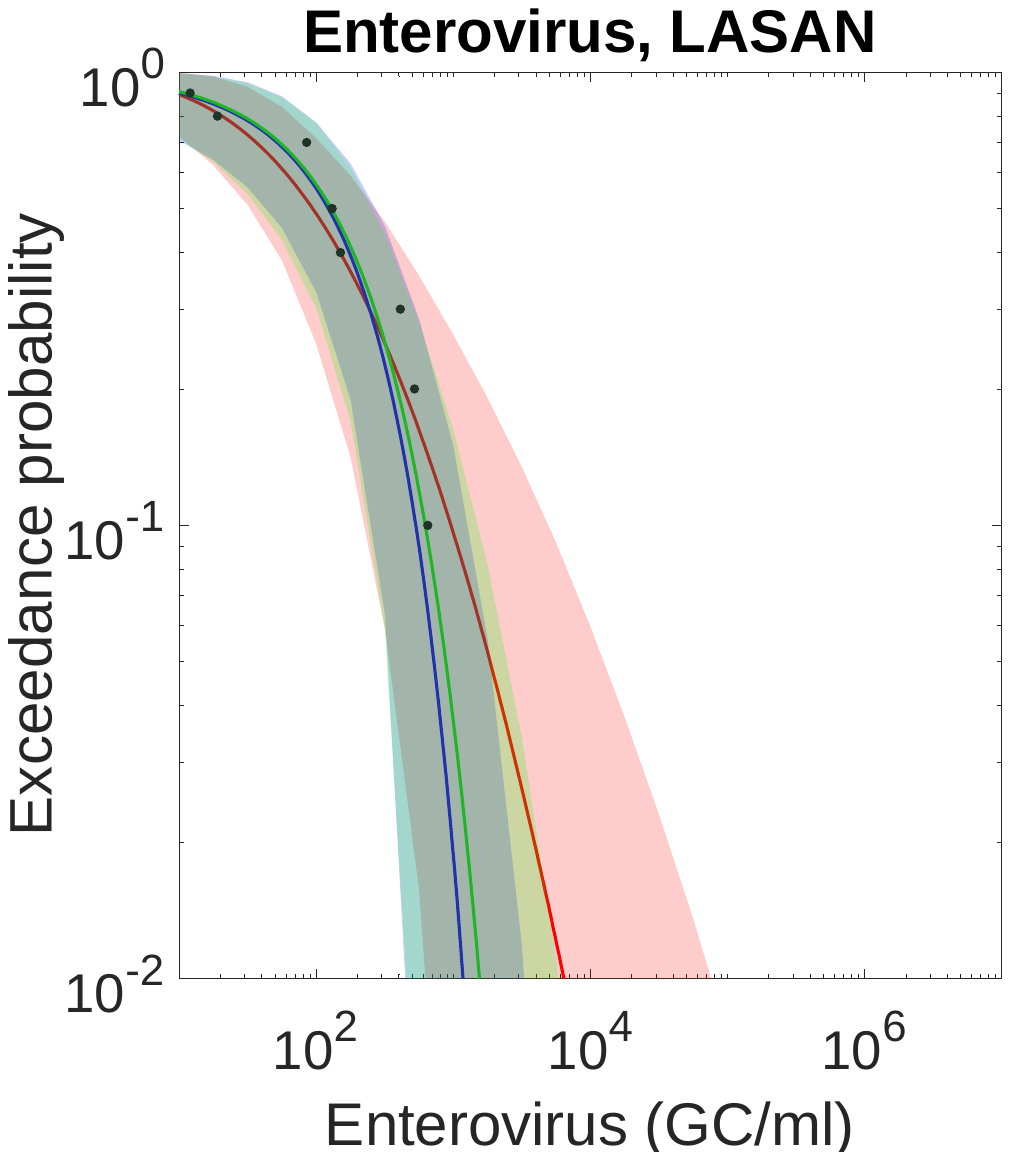

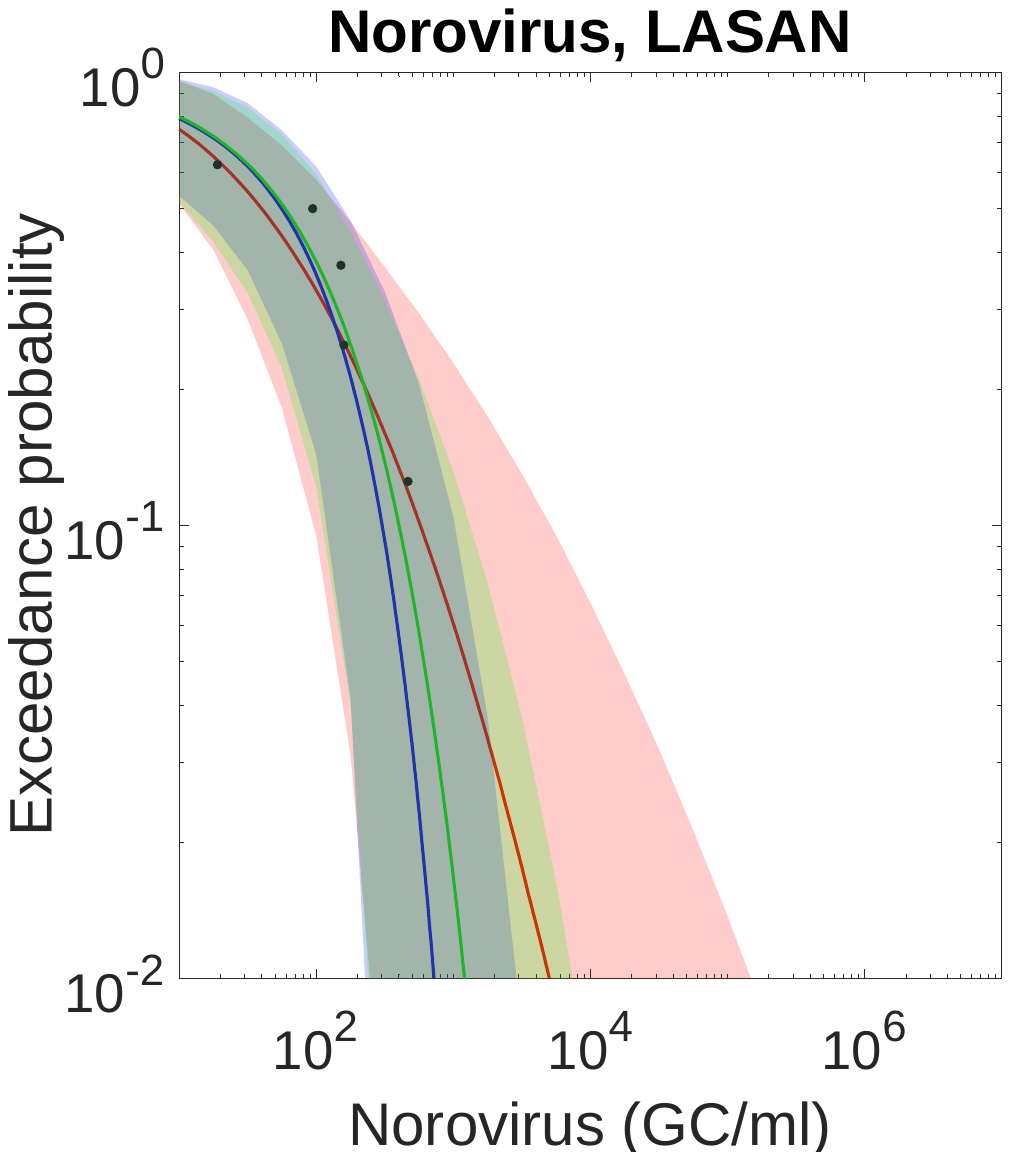

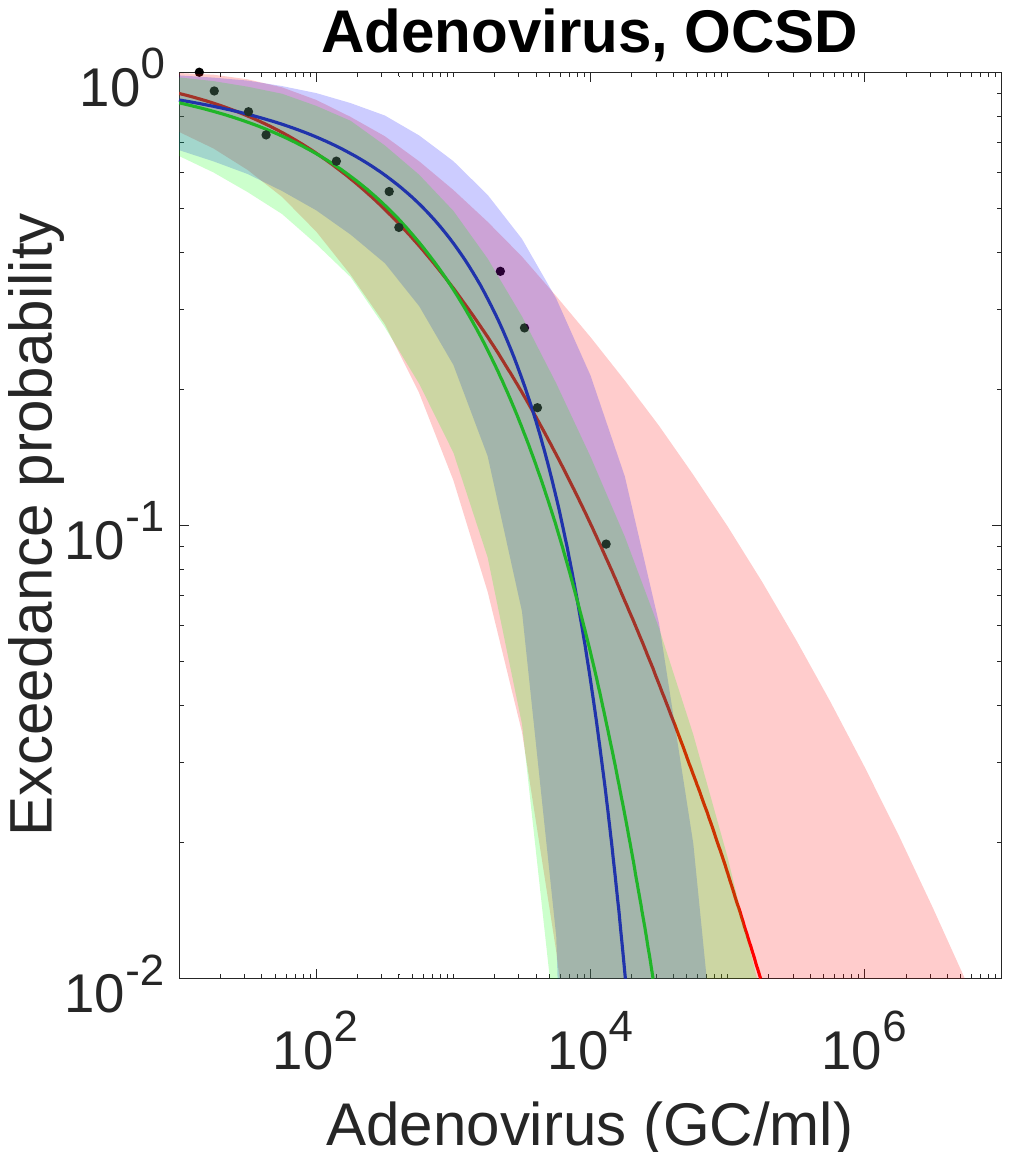

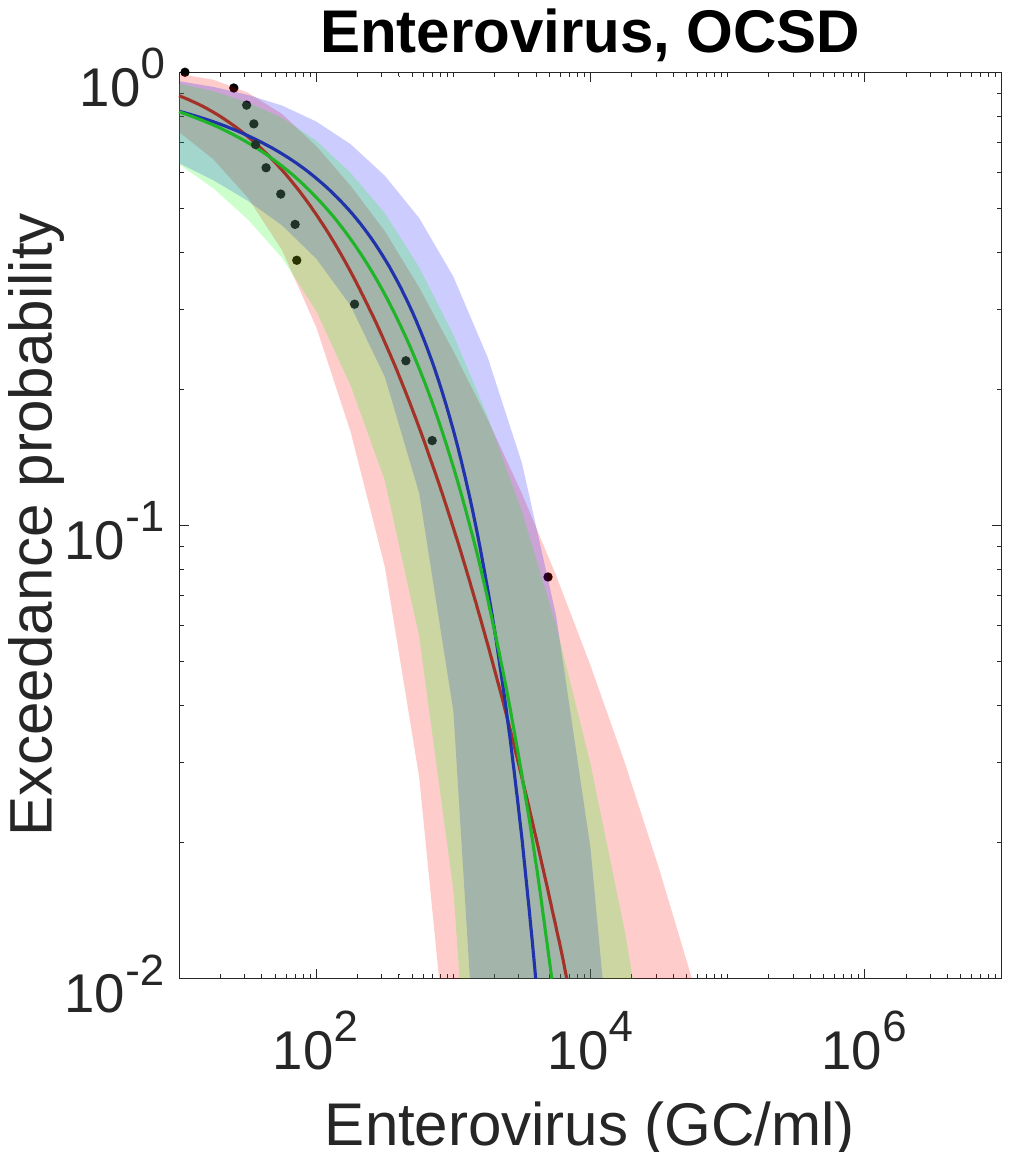

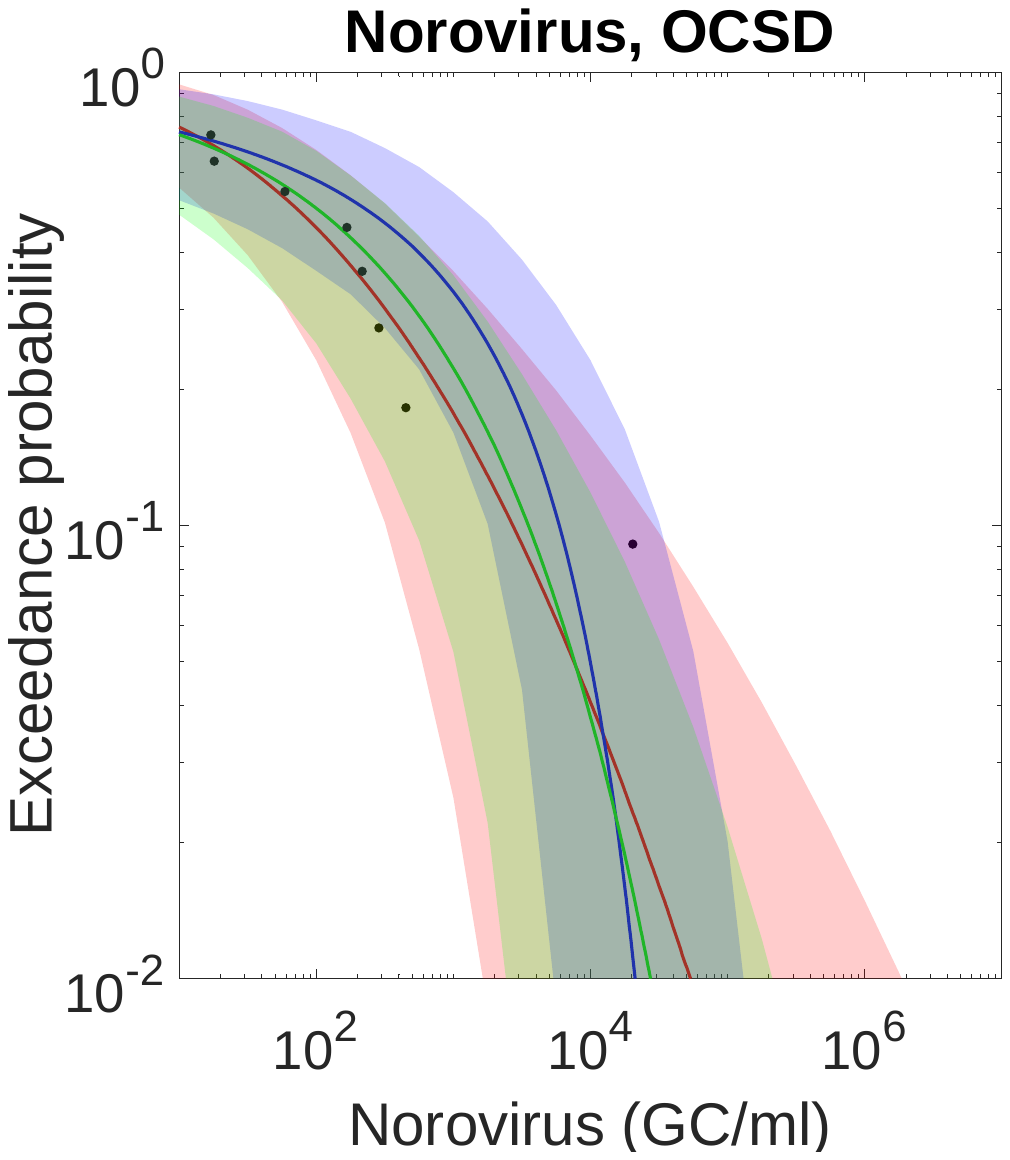

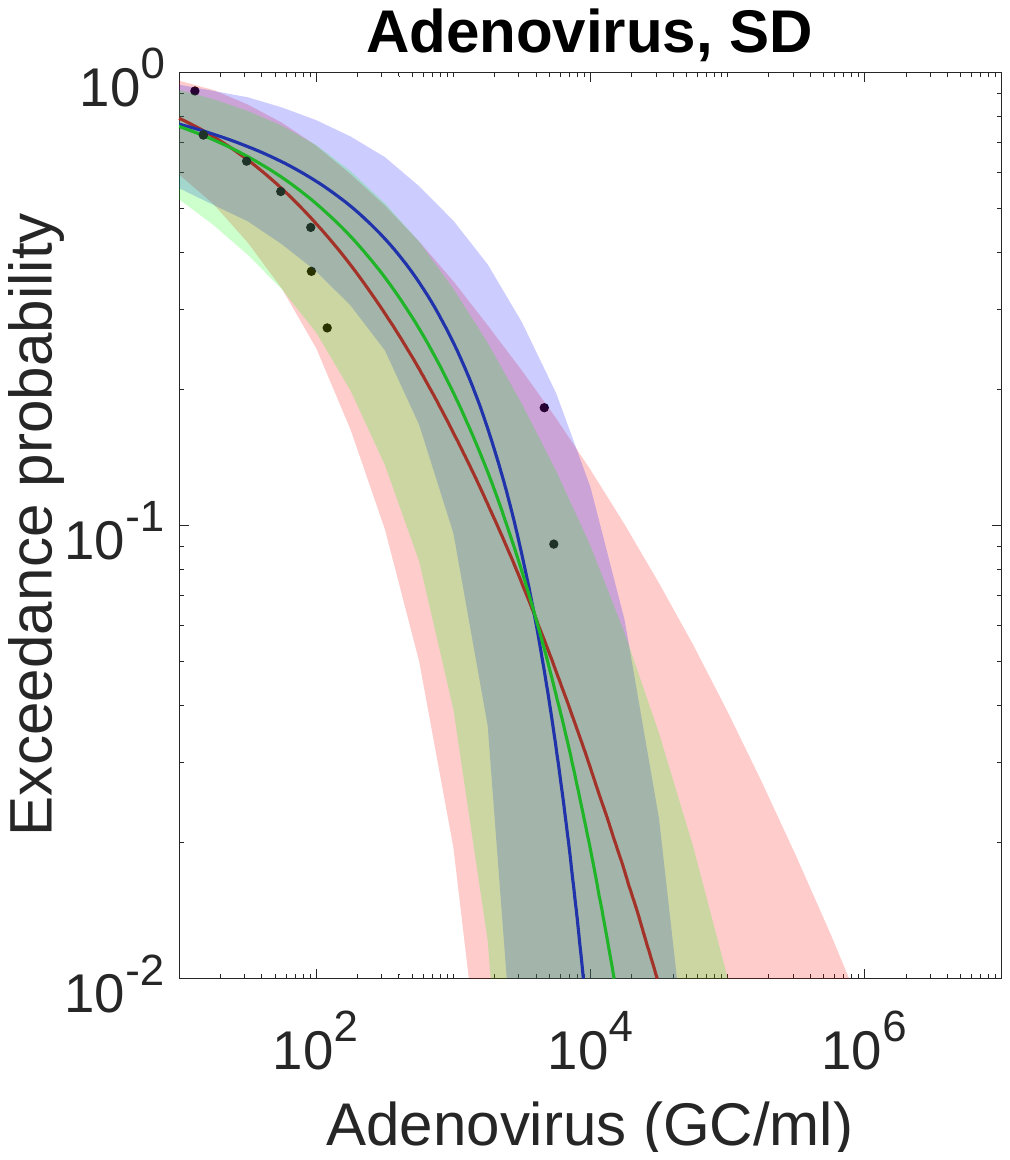

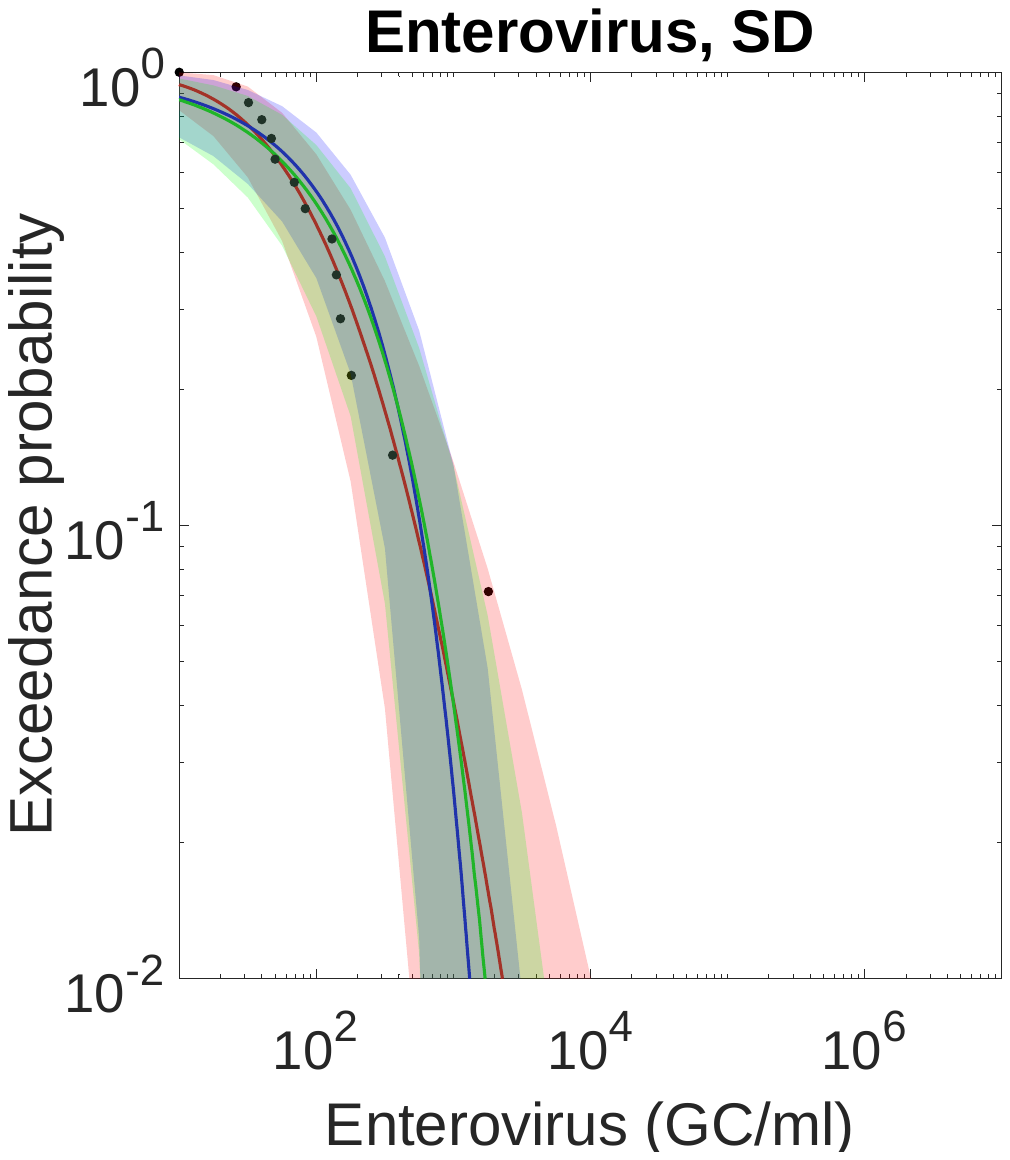

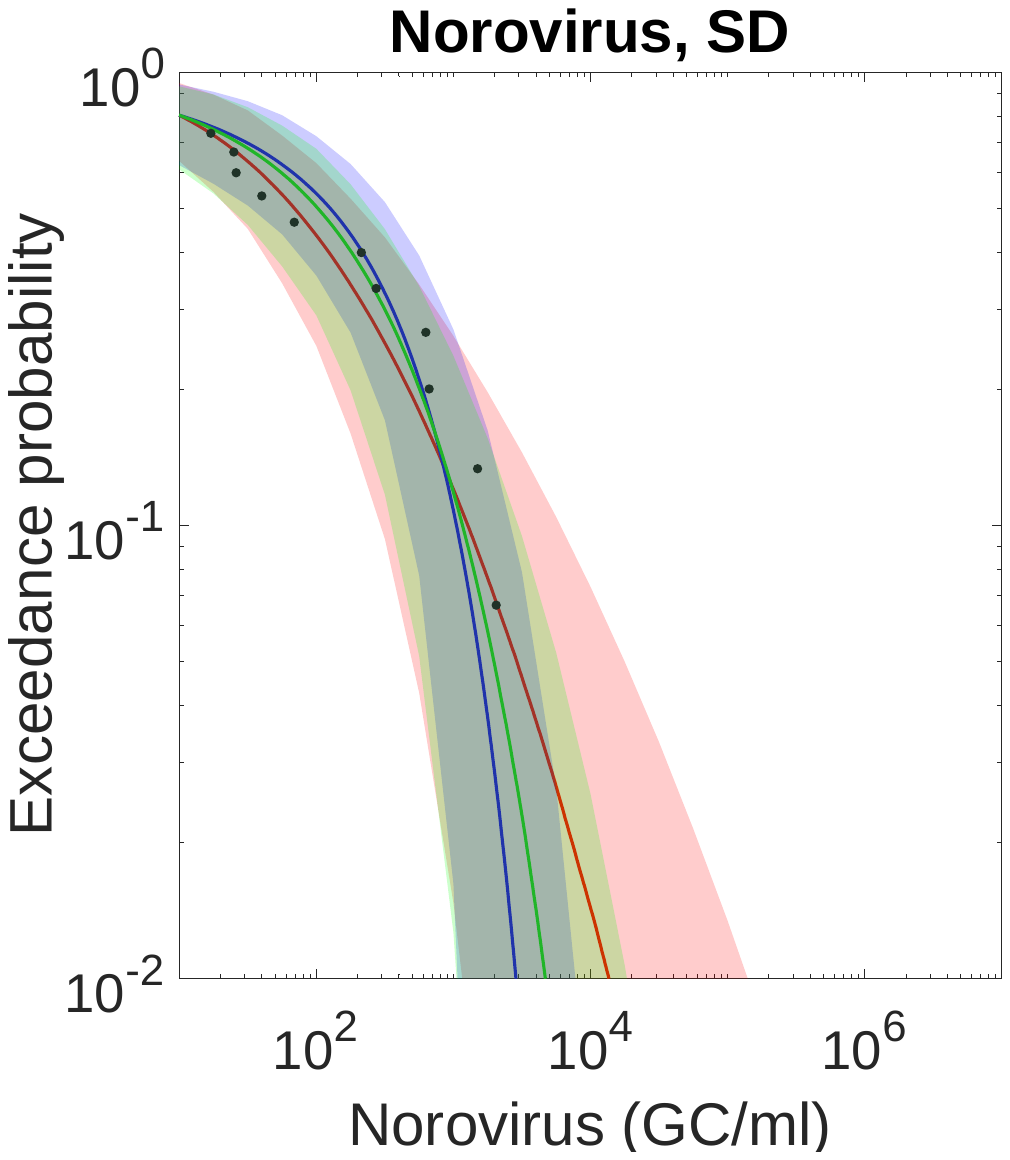

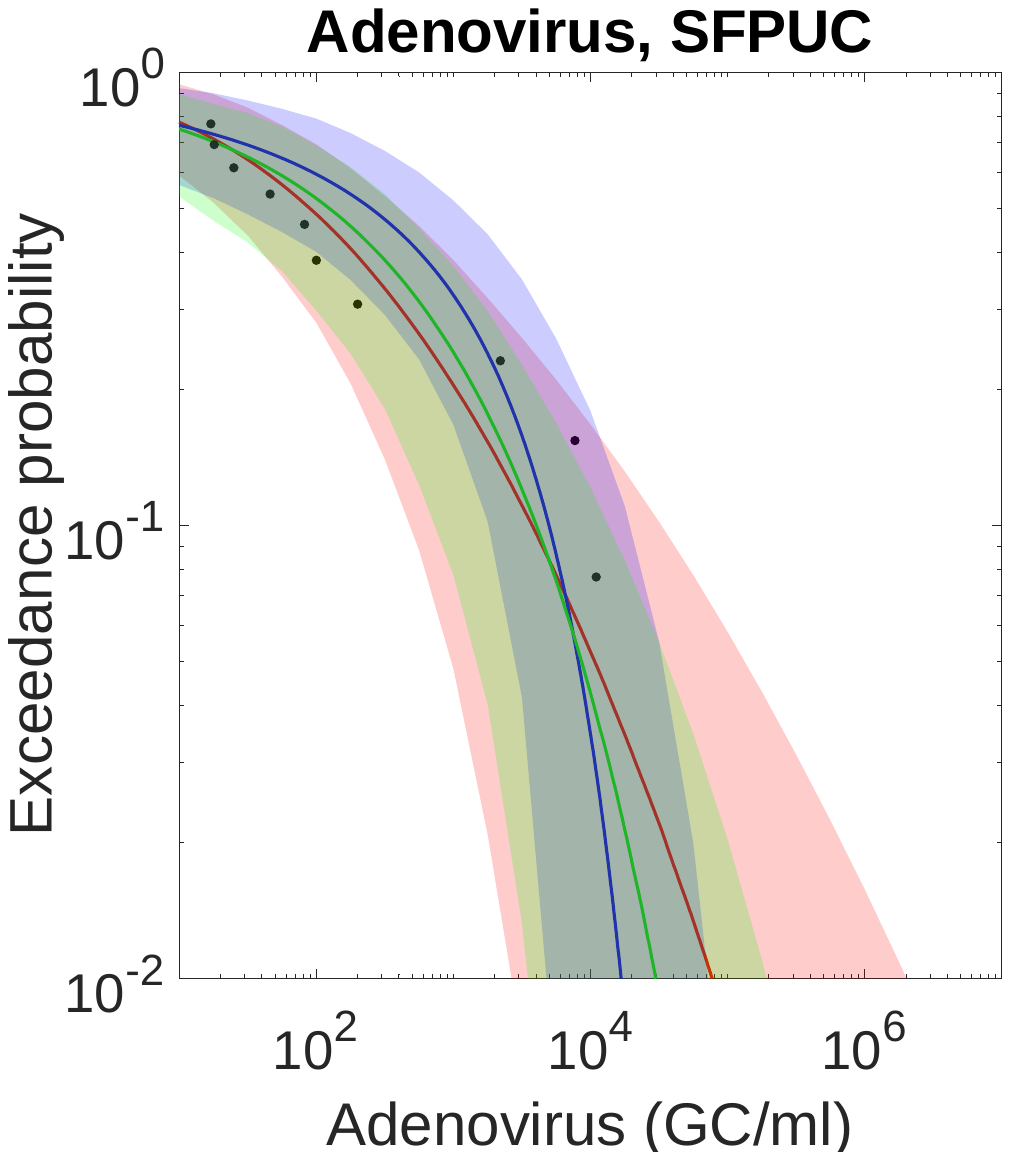

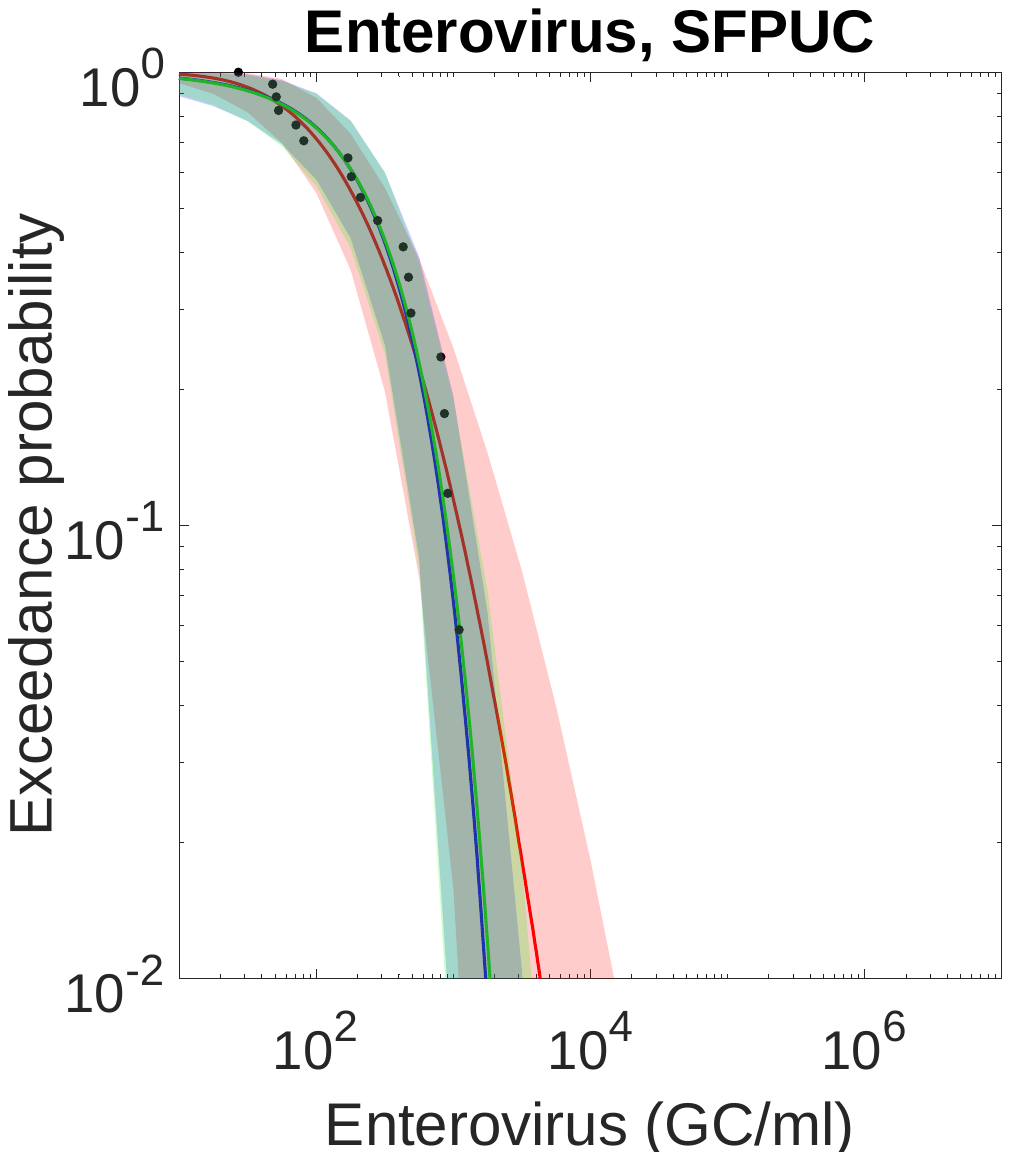

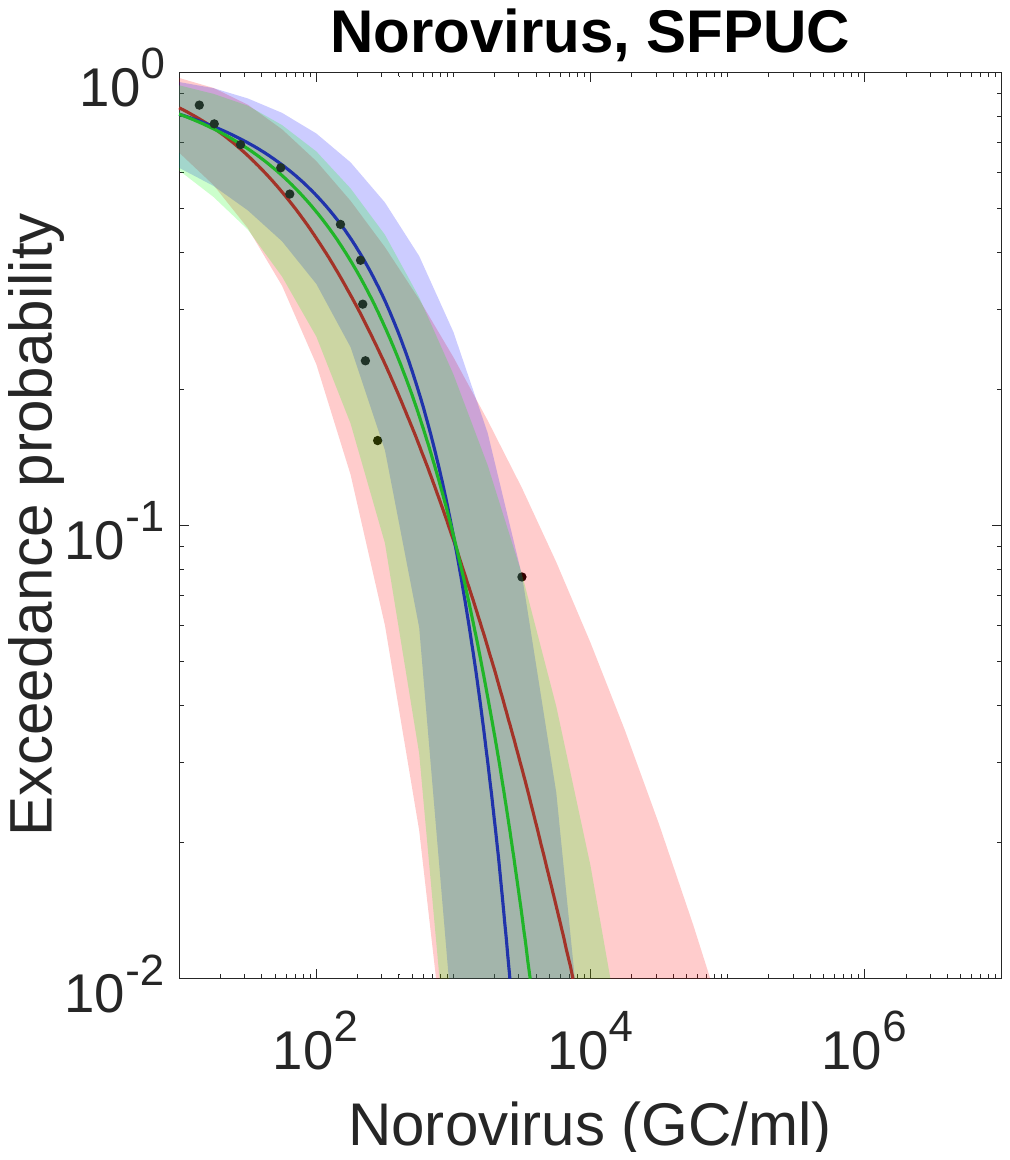

**Fig. S4**. Complementary cumulative distribution functions (CCDFs) of the continuous distributions of enteric virus concentrations for the eight wastewater treatment plants. Red curves represent lognormal distributions, blue curves represent gamma distributions, and green curves represent Weibull distributions.

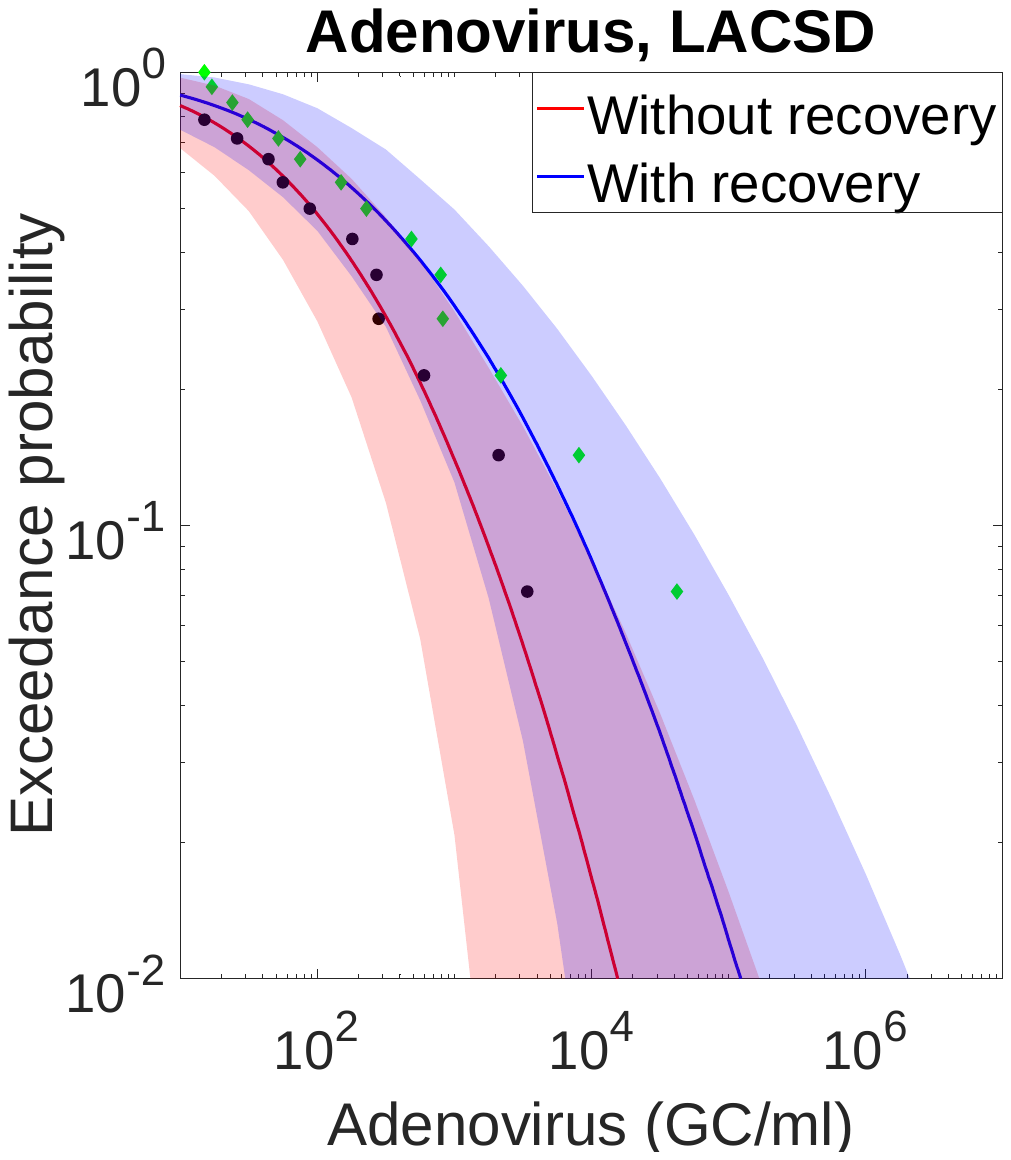

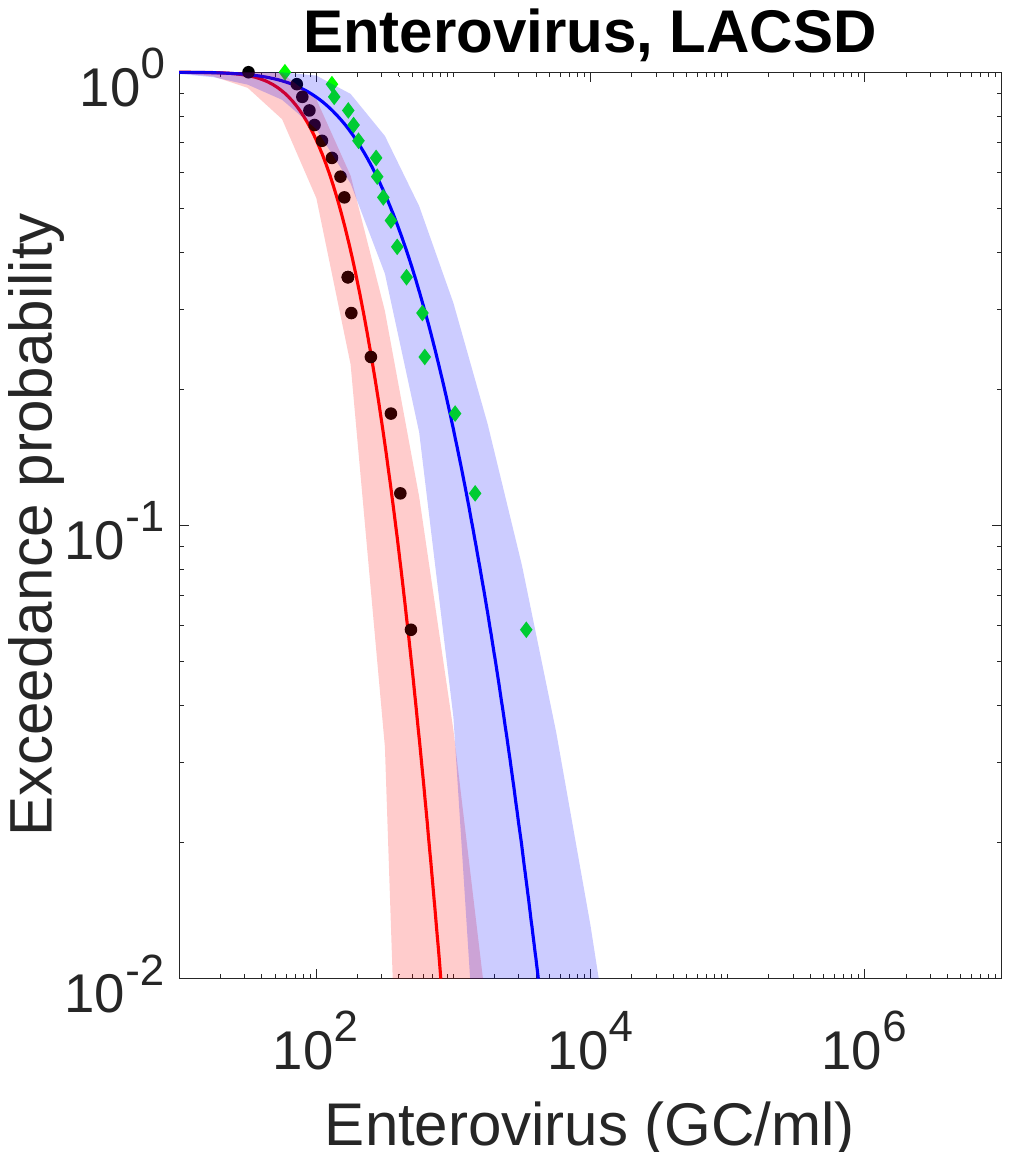

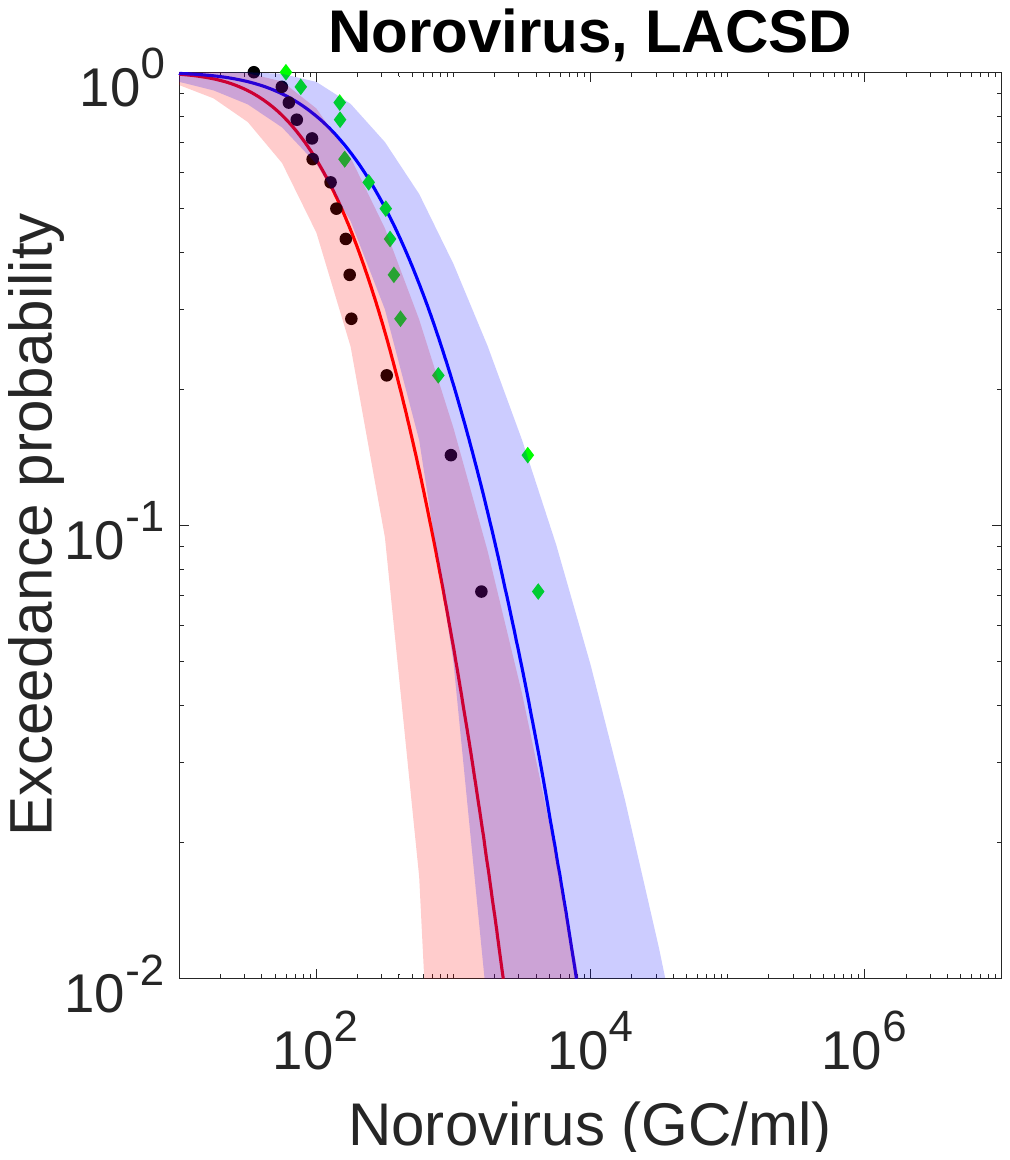

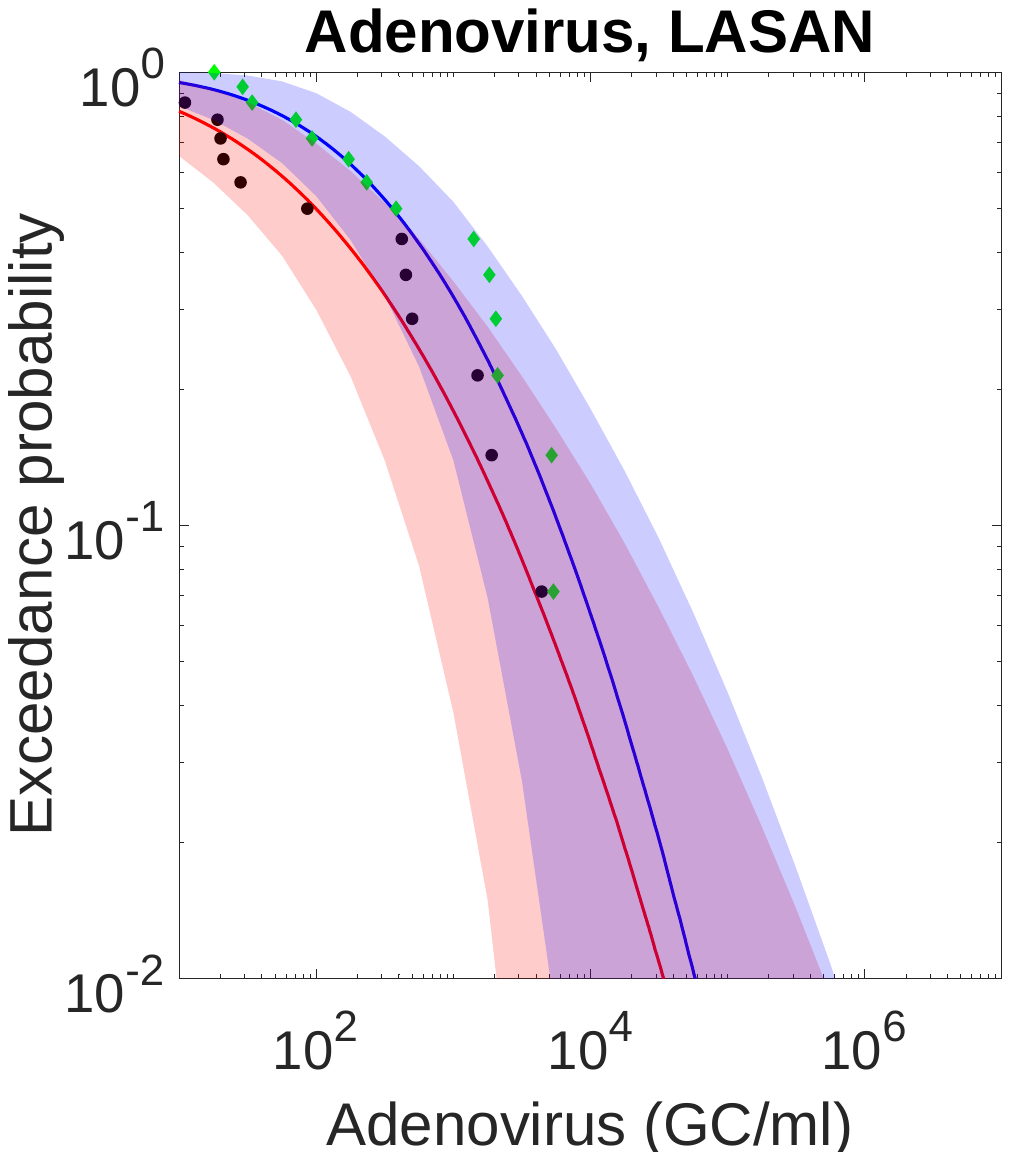

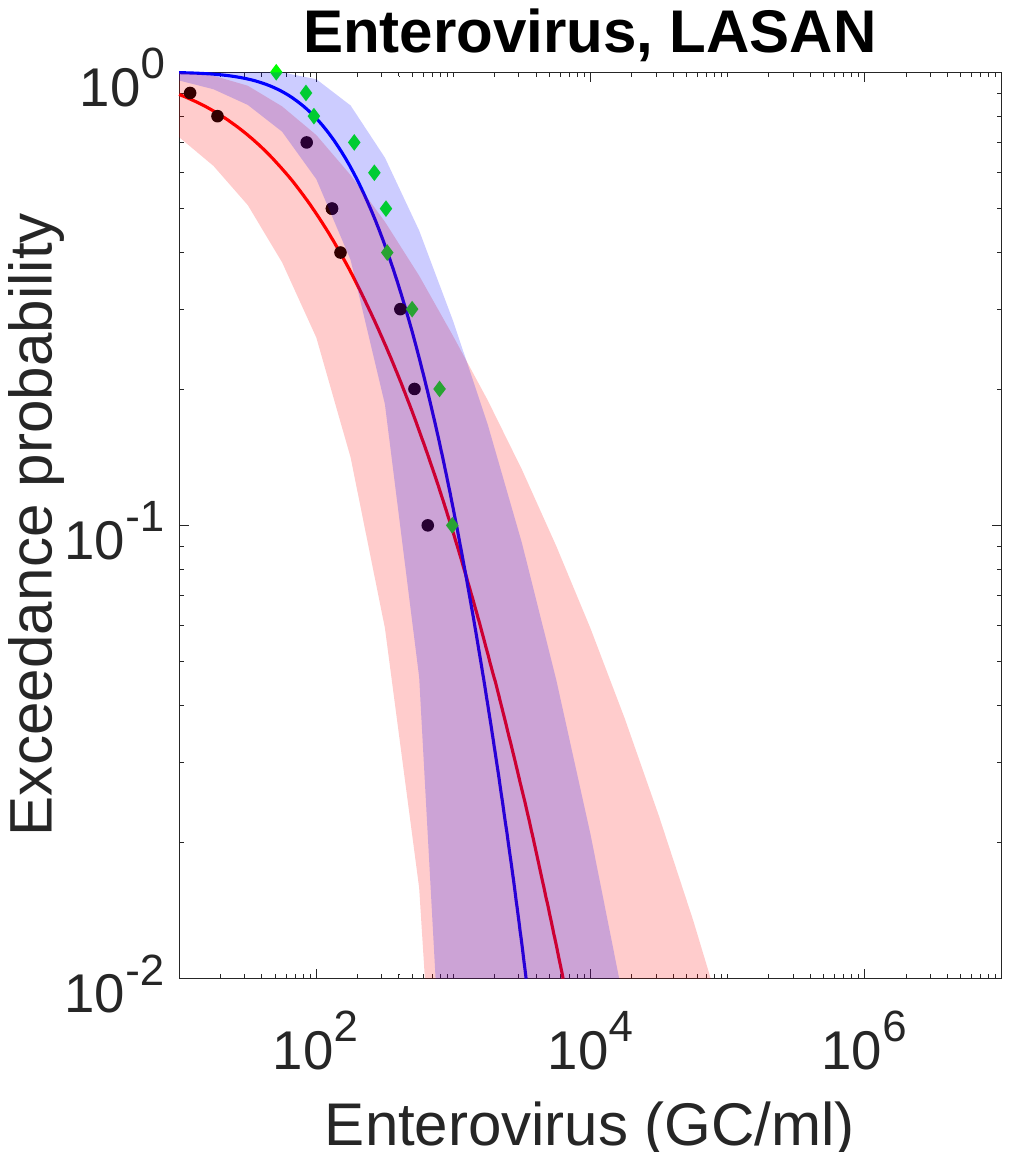

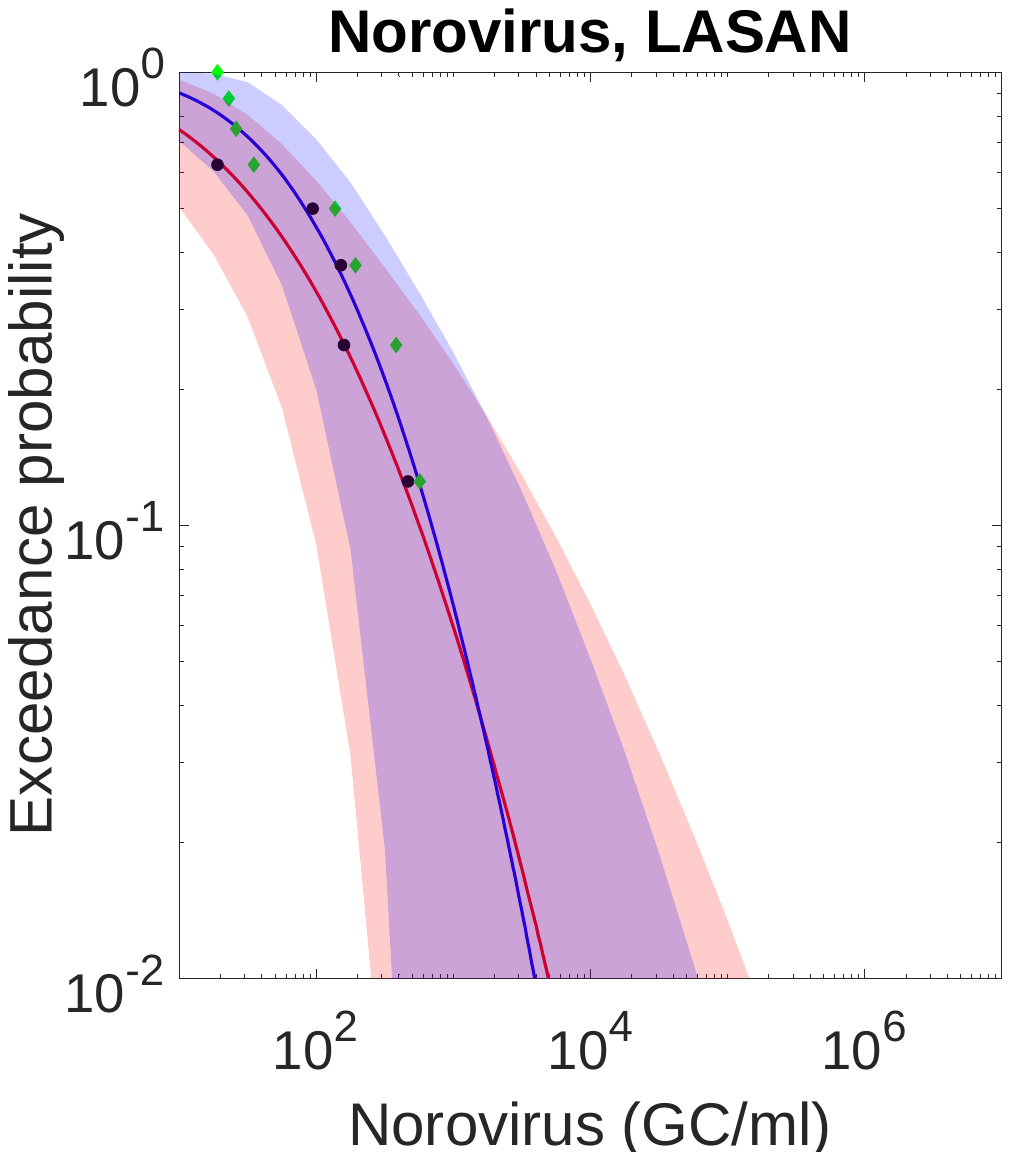

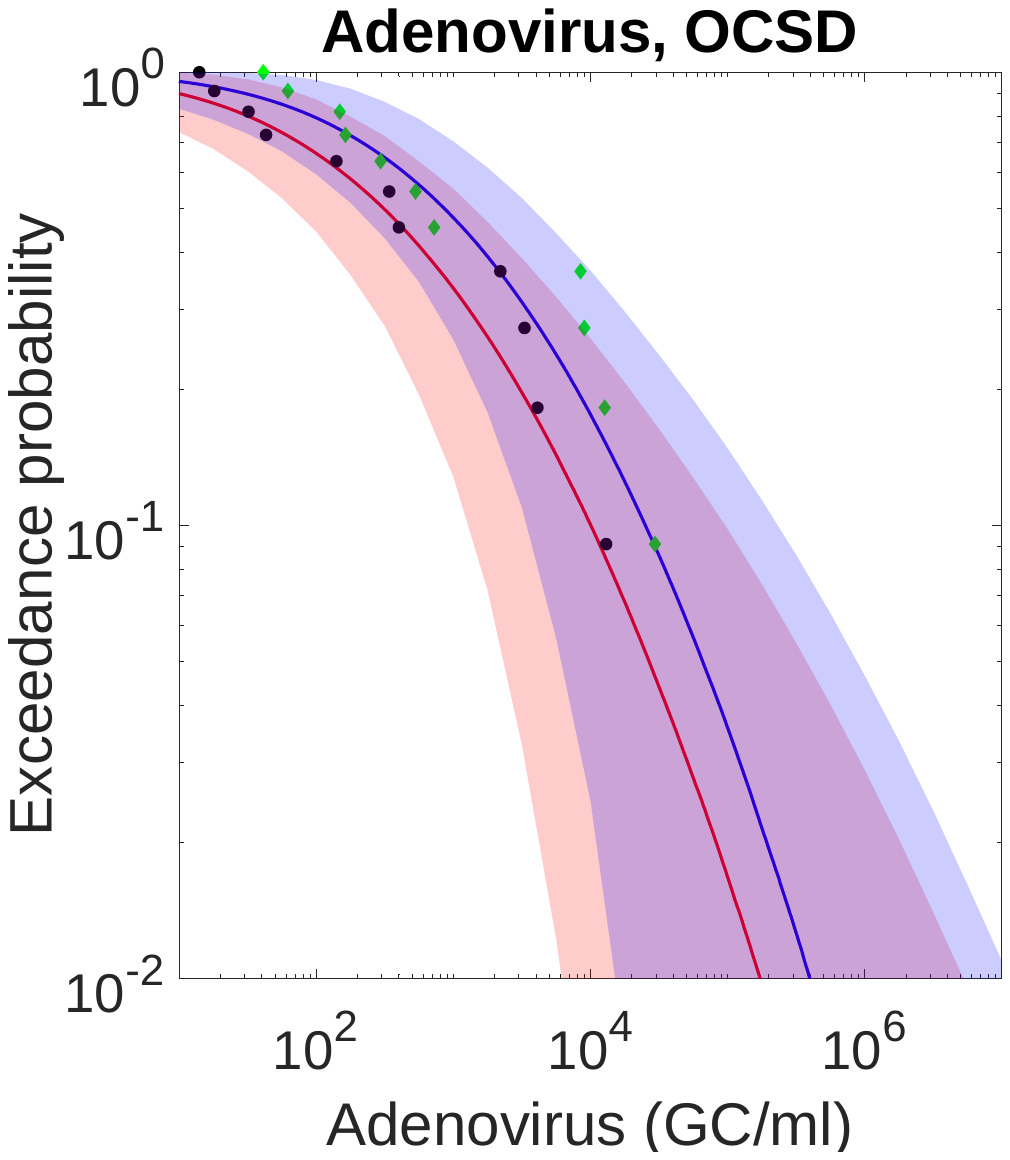

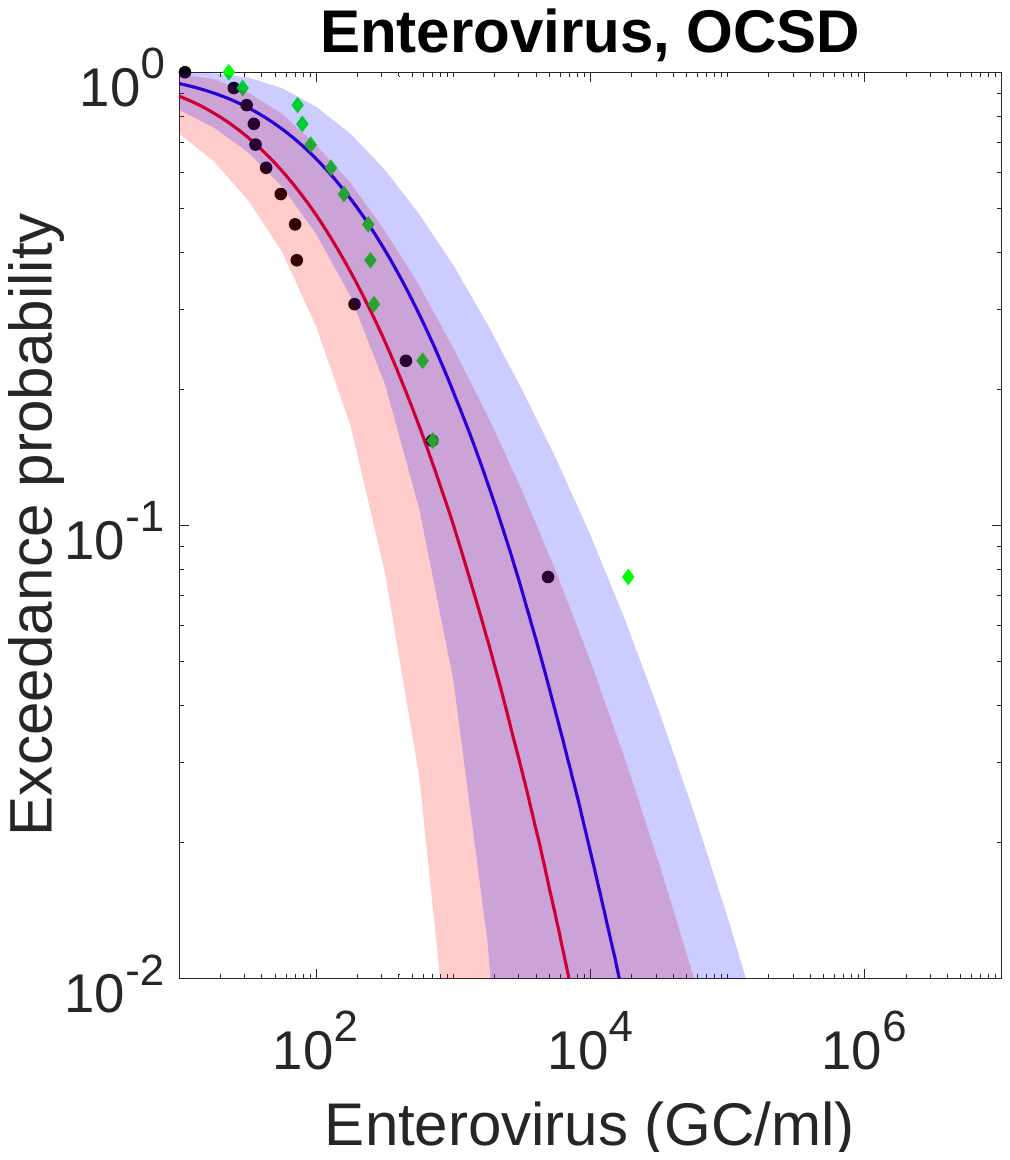

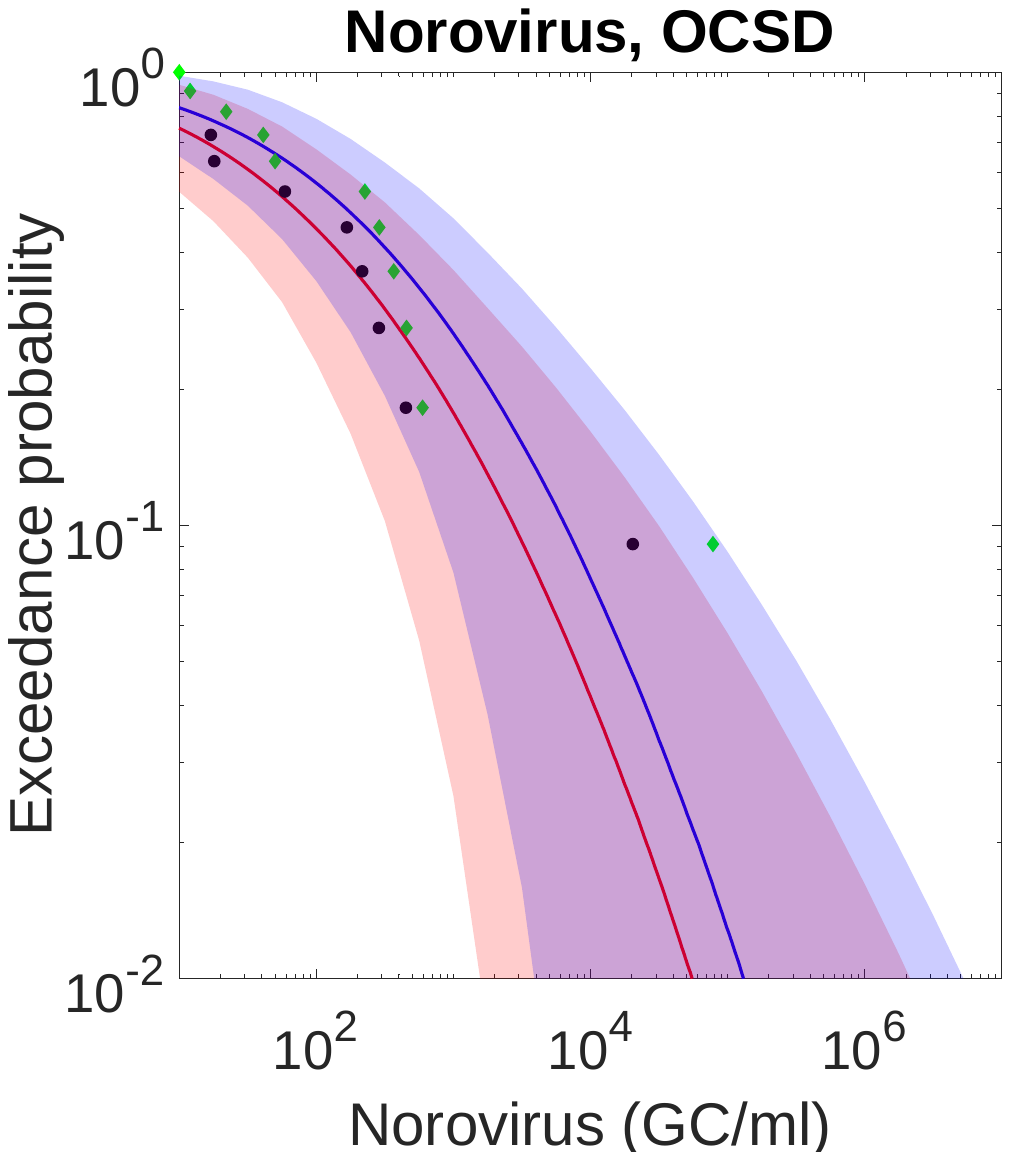

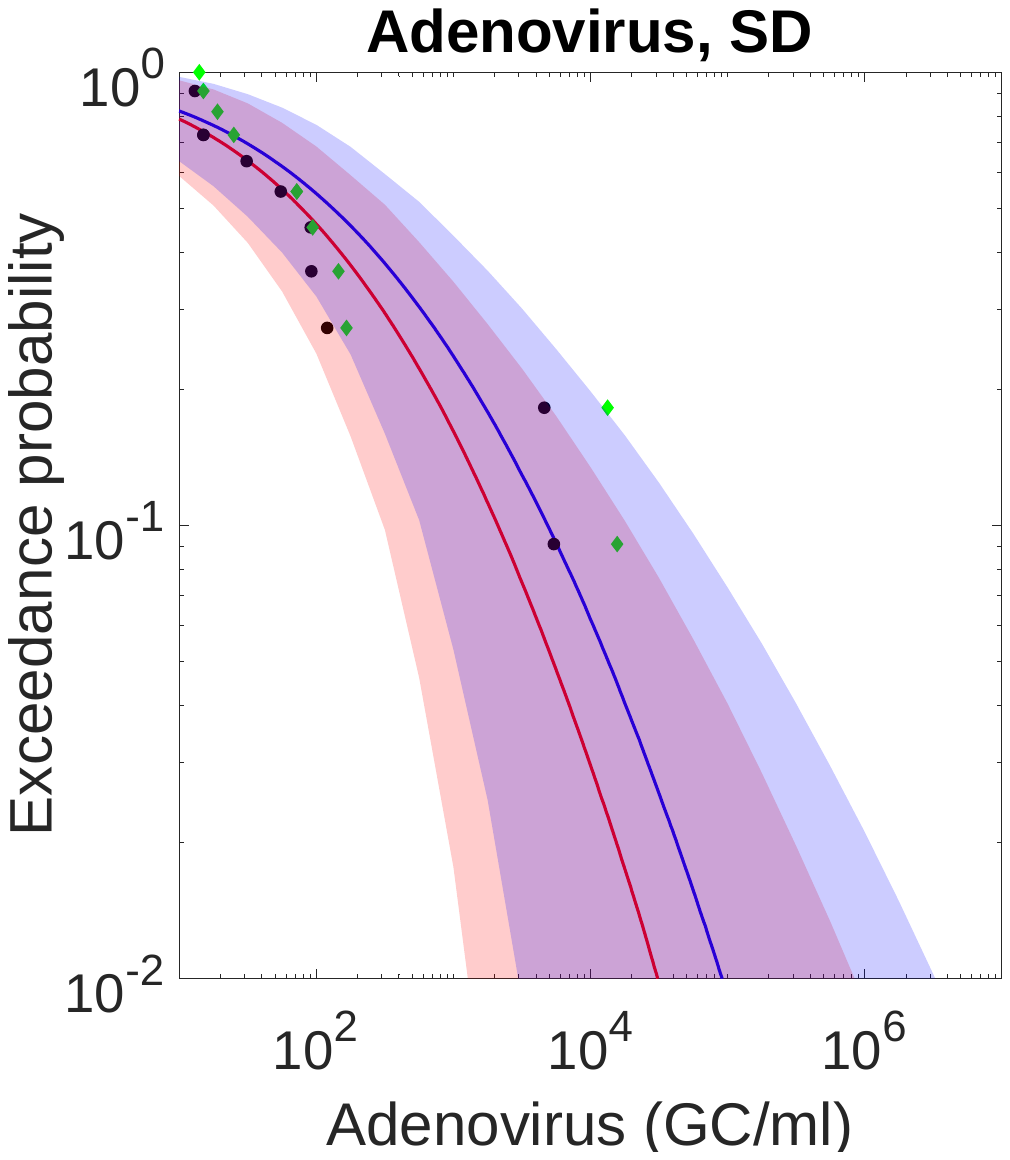

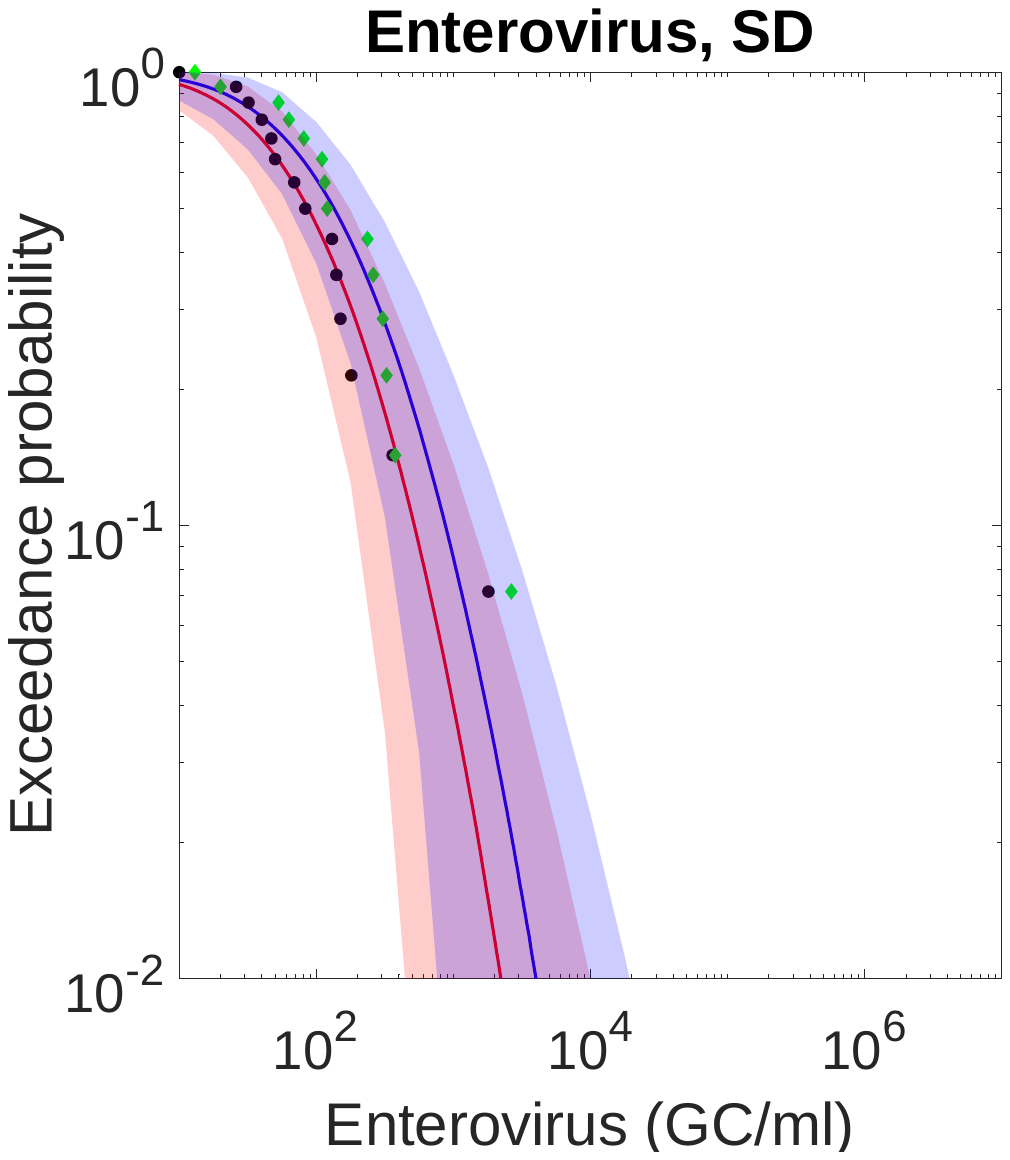

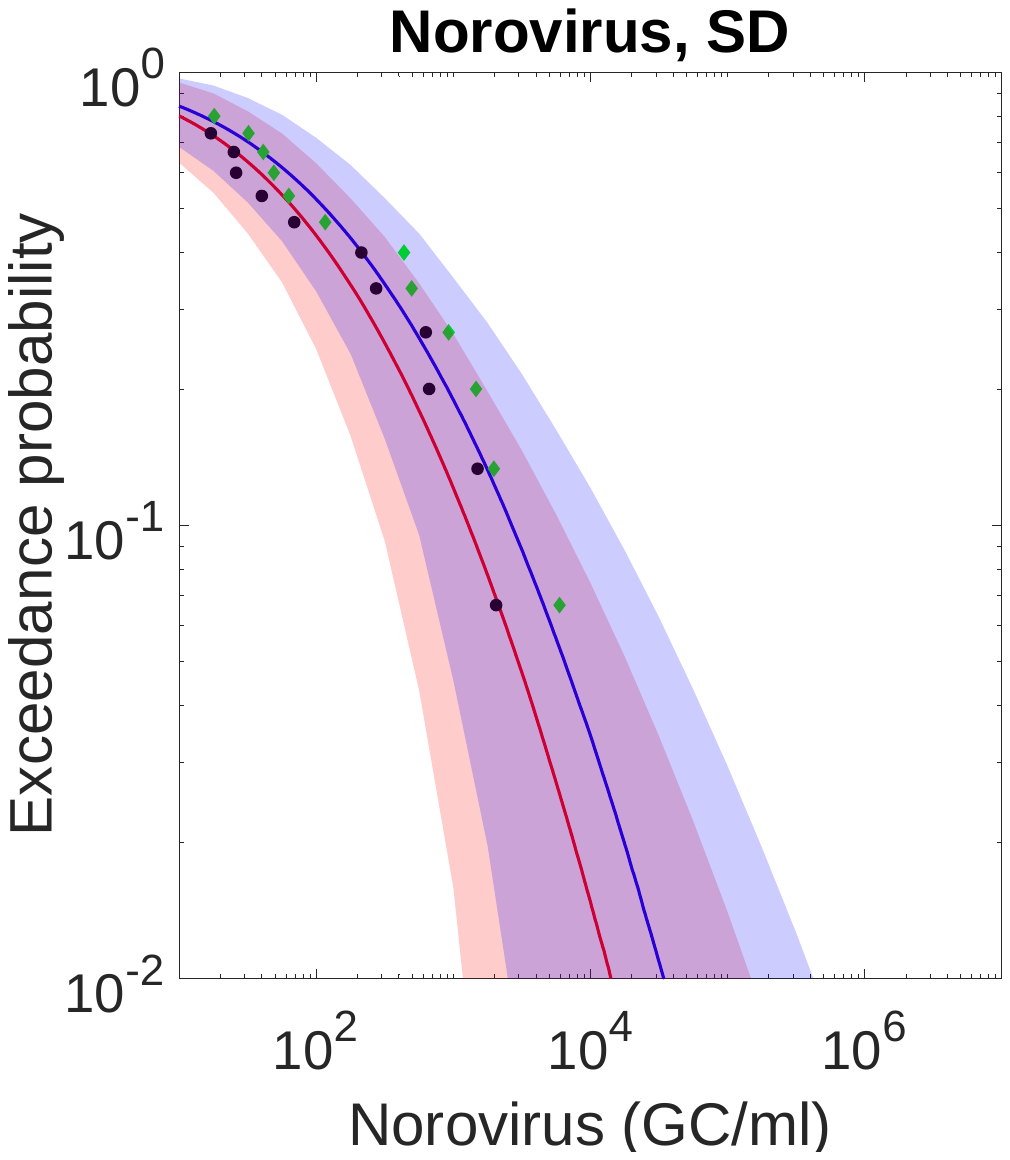

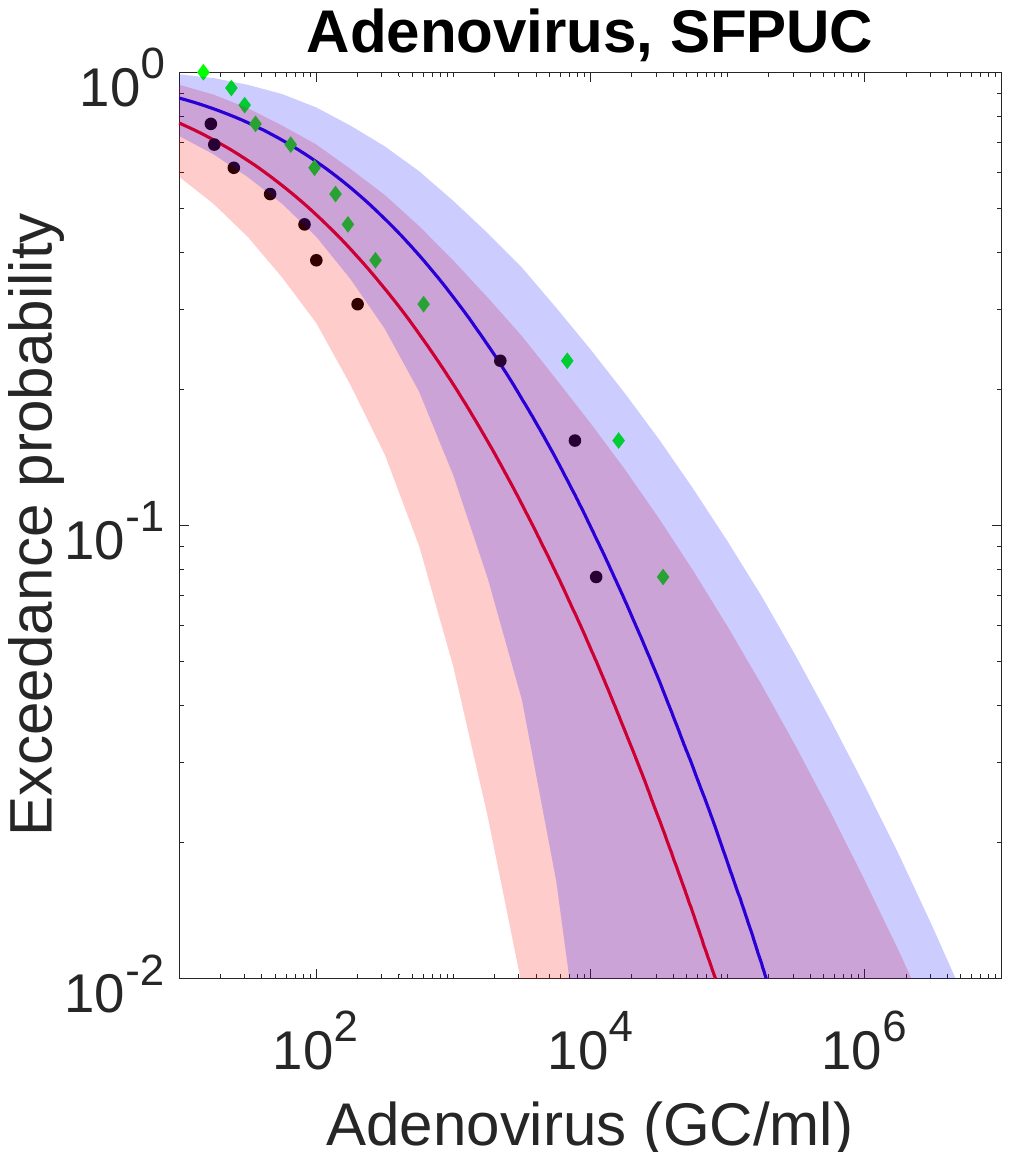

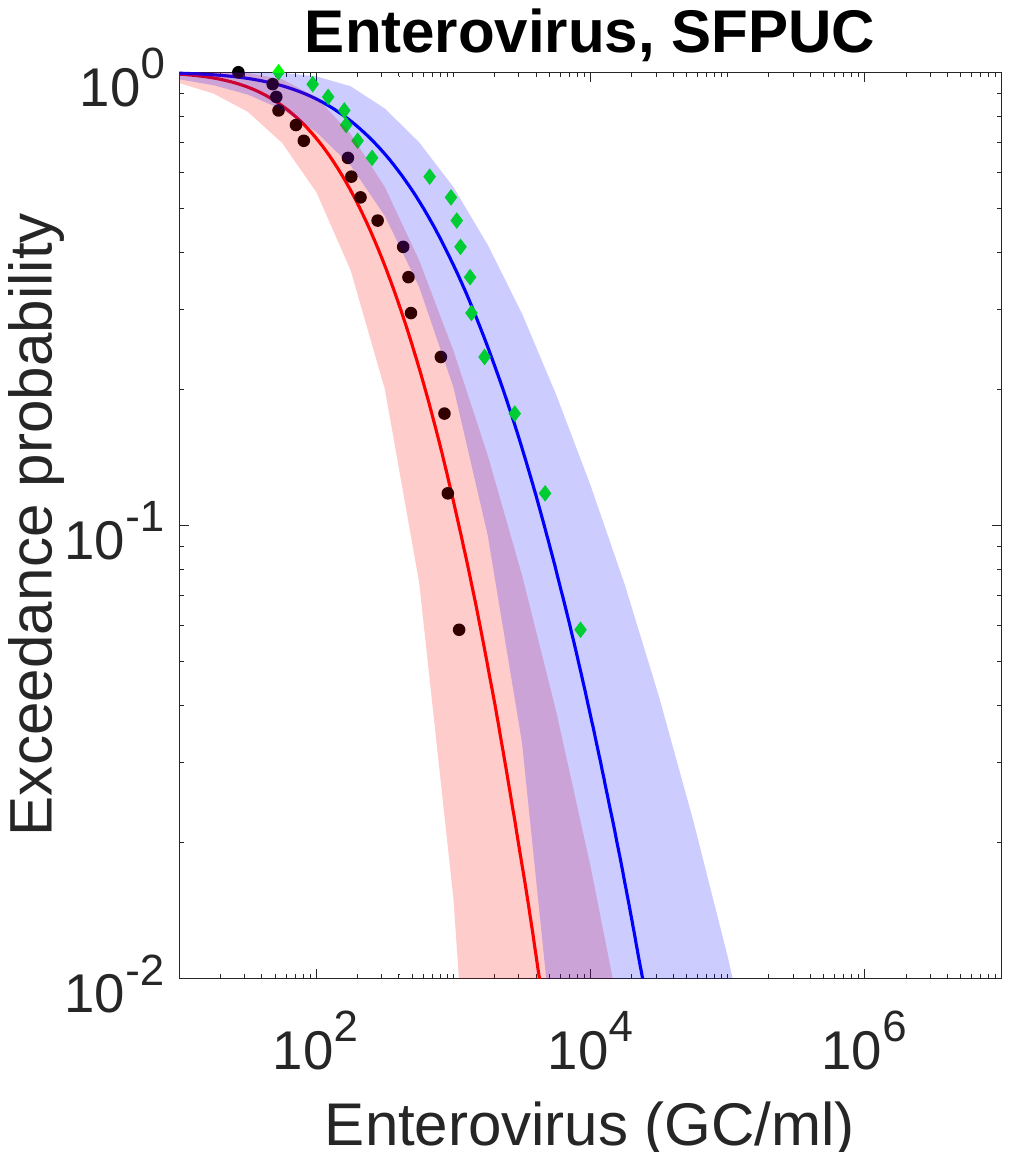

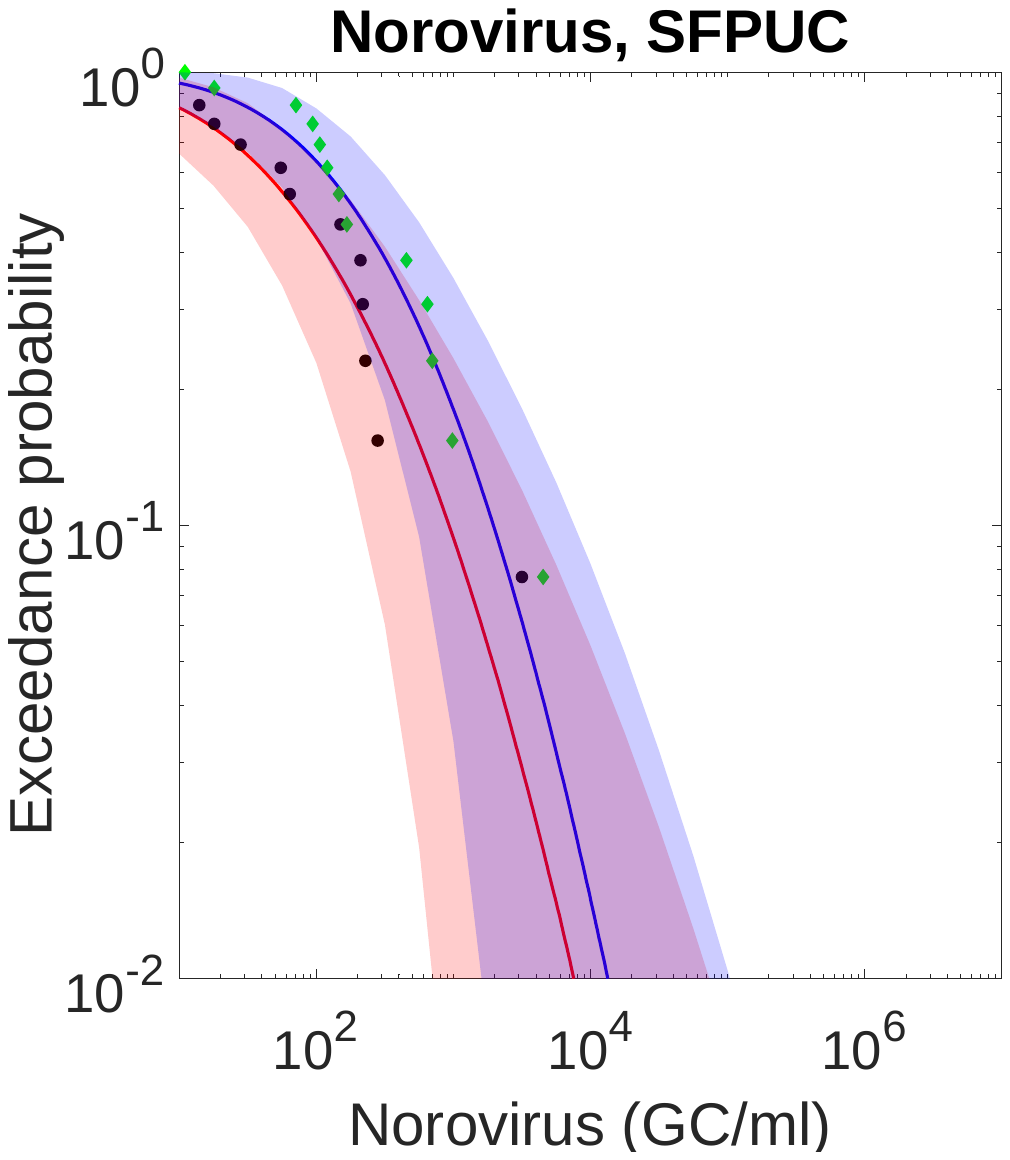

**Fig. S5**. Complementary cumulative distribution functions (CCDFs) of enteric virus concentrations with and without correcting for sample-specific recovery rates for the five wastewater treatment plants (WWTPs) from California, USA.

**Table S1.** mDIC of the mixed Poisson distributions and the DIC and AIC of the continuous distributions for wastewater treatment plants from Switzerland and Japan.

| Place | Lausanne, Switzerland | | | | Zürich, Switzerland Matsushima, Japan | | |
| --- | --- | --- | --- | --- | --- | --- | --- |
| DICm | AD | ET | NR | RT | ET | NR | NR |
| PLN | 179 | 176 | 253 | 269 | 1632 | 2348 | 851 |
| PGA | 181 | 182 | 247 | 280 | 1672 | 2352 | 922 |
| DIC | AD | ET | NR | RT | ET | NR | NR |
| Lognormal | 180 | 176 | 237 | 244 | 2968 | 2691 | 2648 |
| Gamma | 180 | 177 | 235 | 243 | 2969 | 3668 | 2684 |
| Weibull | 180 | 177 | 236 | 244 | 2977 | 3668 | 2663 |
| AIC | AD | ET | NR | RT | ET | NR | NR |
| Lognormal | 178 | 173 | 234 | 241 | 2967 | 3689 | 2651 |
| Gamma | 177 | 174 | 231 | 240 | 2974 | 3665 | 2686 |
| Weibull | 177 | 174 | 233 | 241 | 2976 | 3665 | 2664 |

**Table S2.** mDIC of the mixed Poisson distributions and the DIC and AIC of the continuous distributions for wastewater treatment plants from California, USA.

| Place | LACSD | | | LASAN | | | OCSD | | | | | | SD | | | | | | SFPUC | | | | | |
| --- | --- | --- | --- | --- | --- | --- | --- | --- | --- | --- | --- | --- | --- | --- | --- | --- | --- | --- | --- | --- | --- | --- | --- | --- |
| DICm | AD | ET | NR | AD | ET | NR | AD | | ET | | NR | | AD | | ET | | NR | | AD | | ET | | NR | |
| PLN | 271 | 207 | 209 | 223 | 130 | 93 | | 203 | | 205 | | 169 | | 173 | | 217 | | 200 | | 216 | | 237 | | 197 |
| PGA | 279 | 213 | 220 | 279 | 121 | 95 | 241 | | 247 | | 171 | | 236 | | 239 | | 211 | | 232 | | 236 | | 212 | |
| DIC | AD | ET | NR | AD | ET | NR | AD | | ET | | NR | | AD | | ET | | NR | | AD | | ET | | NR | |
| Lognormal | 382 | 444 | 379 | 389 | 270 | 205 | 332 | | 351 | | 301 | | 301 | | 369 | | 403 | | 363 | | 474 | | 346 | |
| Gamma | 387 | 443 | 385 | 392 | 269 | 206 | 334 | | 360 | | 312 | | 308 | | 375 | | 406 | | 370 | | 473 | | 352 | |
| Weibull | 419 | 509 | 423 | 414 | 295 | 211 | 347 | | 380 | | 321 | | 320 | | 404 | | 420 | | 384 | | 525 | | 372 | |
| AIC | AD | ET | NR | AD | ET | NR | AD | | ET | | NR | | AD | | ET | | NR | | AD | | ET | | NR | |
| Lognormal | 187 | 206 | 182 | 192 | 130 | 92 | 178 | | 168 | | 146 | | 146 | | 172 | | 194 | | 180 | | 236 | | 164 | |
| Gamma | 192 | 206 | 188 | 196 | 128 | 92 | 180 | | 178 | | 156 | | 154 | | 172 | | 196 | | 188 | | 236 | | 170 | |
| Weibull | 190 | 208 | 188 | 194 | 128 | 92 | 178 | | 174 | | 152 | | 150 | | 178 | | 196 | | 184 | | 236 | | 168 | |
